# Cerebellar transcranial alternating current stimulation in the theta band facilitates extinction of learned fear responses

**DOI:** 10.1101/2025.01.13.632735

**Authors:** Andreas Thieme, Zsofia Spisak, Philippe Zeidan, Michael Klein, Enzo Nio, Thomas M. Ernst, Nicolas Diekmann, Sophia Göricke, Sen Cheng, Christian J. Merz, Fatemeh Yavari, Michael A. Nitsche, Giorgi Batsikadze, Dagmar Timmann

## Abstract

Fear extinction is a major component of exposure therapy for anxiety disorders. There is initial evidence that the cerebellum contributes to fear extinction learning, i.e., the ability to learn that certain stimuli are no longer associated with an aversive outcome. So far, however, knowledge of the cerebellum’s role in extinction is scarce. In the present study, 6 Hz cerebellar transcranial alternating current stimulation (ctACS) was used to modulate cerebellar function during extinction learning in young and healthy human participants in an MRI study. A two-day differential fear conditioning paradigm was used with acquisition and extinction training being performed on day 1, and fear extinction recall being tested on day 2. 6 Hz ctACS reduced spontaneous recovery of the initial fear association during recall, stabilizing extinction effects compared to sham ctACS. fMRI data during recall revealed significantly reduced activation in cortical areas involved in initial fear acquisition, such as the anterior cingulate and insula, in the verum ctACS group compared to the sham group. During extinction training, on the other hand, the verum group exhibited more widespread cerebral activation compared to the sham group. Group differences were significant in occipital cortical areas. Although direct stimulation effects cannot be excluded, increased activation in the visual cortex may reflect enhanced encoding and processing of visual information during fear extinction learning. The findings suggest that theta-range oscillatory interactions between the cerebellum and cortical areas support extinction processes and provide causal evidence of the cerebellar role in the human fear extinction network.

**Significance Statement:** While the cerebellum is well known for its role in associative learning, its contribution to fear extinction learning remains underexplored. This study used 6 Hz cerebellar transcranial alternating current stimulation (ctACS) to modulate cerebellar function during extinction training in healthy participants. The present data demonstrate that ctACS enhances cerebral cortical activation during extinction training which is followed by enhanced recall of the extinction memory and subsequently reduced activation of cerebral cortical areas associated to spontaneous recovery of fear memory. These findings provide causal evidence for the cerebelluar involvement in the human fear extinction network and suggest that enhancing cerebellar theta oscillations may be useful to support exposure therapy.

## Introduction

The cerebellum is well known for its contribution to associative learning. The best-known example is eyeblink conditioning, a form of motor learning (De Zeeuw and Ten Brinke, 2015). The cerebellum is also involved in associative learning in the cognitive and emotional domains (Diedrichsen et al., 2019; Guell and Schmahmann, 2020; Schmahmann, 2019). One important emotion is fear. The cerebellum has been shown to contribute to learning of new fear associations (Hwang et al., 2022; Urrutia Desmaison et al., 2023). Recent studies suggest that the cerebellum also contributes to extinction learning (Doubliez et al., 2023). Deficient fear extinction is thought to contribute to a variety of anxiety disorders including posttraumatic stress disorder and others (Sep et al., 2023; VanElzakker et al., 2014).

The cerebellum is well suited to support extinction learning due to its anatomical connections with key regions of the fear extinction network, including the ventromedial prefrontal cortex (vmPFC), hippocampus, insula, striatum, amygdala, and midbrain periaqueductal gray (Boll et al., 2013; Li et al., 2011; Li and McNally, 2014). In functional brain imaging (fMRI) studies activations of the cerebellum are often found related to fear extinction learning (Batsikadze et al., 2022; Chang et al., 2015). There is also initial evidence that extinction learning is reduced in patients with cerebellar disease (Batsikadze et al., 2024; Maschke et al., 2002).

However, knowledge about how the cerebellum may contribute to extinction learning remains limited. Recent work in rodents shows that the cerebellum regulates fear extinction via monosynaptic projections from the fastigial nuclei to the periaqueductal gray (Frontera et al., 2020; Lawrenson et al., 2022), but also through thalamo-prefrontal interactions (Frontera et al., 2023).

Fear extinction learning is thought to be driven by prediction errors which are elicited by the unexpected omission of the aversive unconditioned stimulus (US) (Rescorla and Wagner, 1972). In healthy human participants, the unexpected omission of the aversive US results in strong cerebellar activation, suggesting that the cerebellum is involved in processing of prediction errors (Ernst et al., 2019). Batsikadze et al. (2024) tested cerebellar patients using fMRI. During extinction training and recall, controls showed activations related to the unexpected omission toward the CS+, whereas patients did not. Given that the cerebellar fMRI primarily reflects mossy fiber input (Diedrichsen et al., 2010), these findings suggest that cerebellar cortical degeneration alters input signals related to the unexpected omission of the aversive US.

In this study, cerebellar transcranial alternating current stimulation (ctACS) was used to modulate cerebellar function during extinction learning in healthy human participants in an MR scanner (Wessel et al., 2023). tACS, when administered at frequencies matching natural neuronal oscillations, can synchronize (“entrain”) intrinsic oscillations and enhance functional connectivity (Antal and Herrmann, 2016; Bachinger et al., 2017). Animal studies show that this is equally the case for the cerebellum (Asan et al., 2020; Kang et al., 2023).

We targeted cerebellar theta oscillations with 6 Hz ctACS, aligning with the response frequency of cerebellar granule and Golgi cells (D’Angelo et al., 2013; Gandolfi et al., 2013). Theta activity is well known to contribute to visuomotor adaptation and associative learning in humans and animals (Jonker et al., 2021; Tzvi et al., 2022). More specifically, a correlation between spontaneous cerebellar theta activity in the 4-7 Hz frequency range and successful extinction of conditioned eyeblink responses has been found in guinea pigs (Wang et al., 2014), Furthermore, a decrease in cerebellar theta activity following exposure to the conditioned stimulus has been linked to the spontaneous reappearance of previously extinguished eyeblink responses (Wang et al., 2019).

We found that theta ctACS during extinction training strengthened extinction recall, which was accompanied by significant changes of cerebral activations during extinction learning and recall. These findings provide further support that the cerebellum is involved in fear extinction learning, and modulates cerebral areas involved in fear extinction.

## Materials and Methods

### Subjects

A total of 55 young, healthy, right-handed, non-smoking participants (29 men / 26 women, 23.5 ± 3.5 years) was recruited to participate in the experiment. Forty participants (22 males / 18 females, 23.9 ± 3.5 years) were included in the final SCR data analysis, and 35 participants (19 men / 16 women, 23.7 ± 3.2 years) in the final fMRI data analysis. Six participants had to be excluded due to technical errors, e.g. volume orientation mix-ups or loss of adjustment volume settings, six participants due to moderate or higher depression, anxiety or stress scores based on the DASS-21 questionnaire (Lovibond et al., 1995), one participant was excluded for not attending the experiment on the second day, one was excluded due to an incidental finding (cavernoma in the temporal lobe) and one for taking oral contraceptives, which were not allowed as part of the study protocol. In addition, five participants had to be excluded due to constant motion and related artifacts throughout MRI acquisition. Their data was kept in the SCR analysis. All participants were right-handed based on the Edinburgh handedness inventory (Oldfield, 1971) and naïve to both brain stimulation and fear learning procedures. Each participant underwent neurological examination prior to the start of the experiment and their depression, anxiety and stress levels were assessed using the DASS-21 questionnaire (Henry and Crawford, 2005; Norton, 2007). Scores of the participants included in the final analysis were within the normal-to-mild range: depression score, median 2 (IQR [interquartile range] 0 – 4, range 0 - 10), anxiety score, median 2 (IQR 0 – 4, range 0 - 8), stress score, median 6 (IQR 2 – 10, range 0 - 12).

Participants who had a history of neurological disease, metallic head implants, present pregnancy or had taken central nervous system- active medication were excluded from the study. Women taking oral contraceptives were also excluded from the study to avoid any effects on fear-conditioning processes caused by changes in circulating sex hormones (Merz et al., 2018; Velasco et al., 2019). Additionally, participants were instructed to refrain from alcohol for at least 24 hours before the experiment. The study was approved by the Ethics Committee of the University Hospital Essen and conforms to the principles laid down in the Declaration of Helsinki. We obtained informed consent from all participants, and they were compensated with 80 Euros for their participation.

### Experimental procedures

The differential fear conditioning paradigm used in this study was based on the publication by Ernst et al. (2019). Differential fear conditioning was performed using two conditioned stimuli (CS): two pictures of black-and-white geometric figures (a square and a diamond shape) of identical brightness (**Figure 1A**). The CS+ was followed by an aversive US (paired CS+/US trial) during fear acquisition training in 62.5% of the trials. The CS- was never followed by the US. The use of the two CS figures was pseudorandomly counterbalanced across participants. The experiment was performed on two consecutive days (**Figure 1B**). Day 1 consisted of three phases: "habituation" (3 CS+ only trials, 3 CS- only trials), "acquisition training" (10 paired CS+/US trials, 6 CS+ only, 16 CS- only trials) and "extinction training " (16 CS+ only trials, 16 CS- only trials). Day 2 consisted of the recall phase (12 CS+ only trials, 12 CS- only trials). The presentation order of trial types in each phase was pseudorandomized, with two restrictions: firstly, the first two and last trials in the acquisition training were paired CS+/US trials, and secondly, the number of events of each kind was kept equal in the first and second halves of each phase. The order of events was identical for all participants during habituation, acquisition and extinction training. During recall, the initial event was counterbalanced between CS+ and CS- trials.

**Figure 1.**
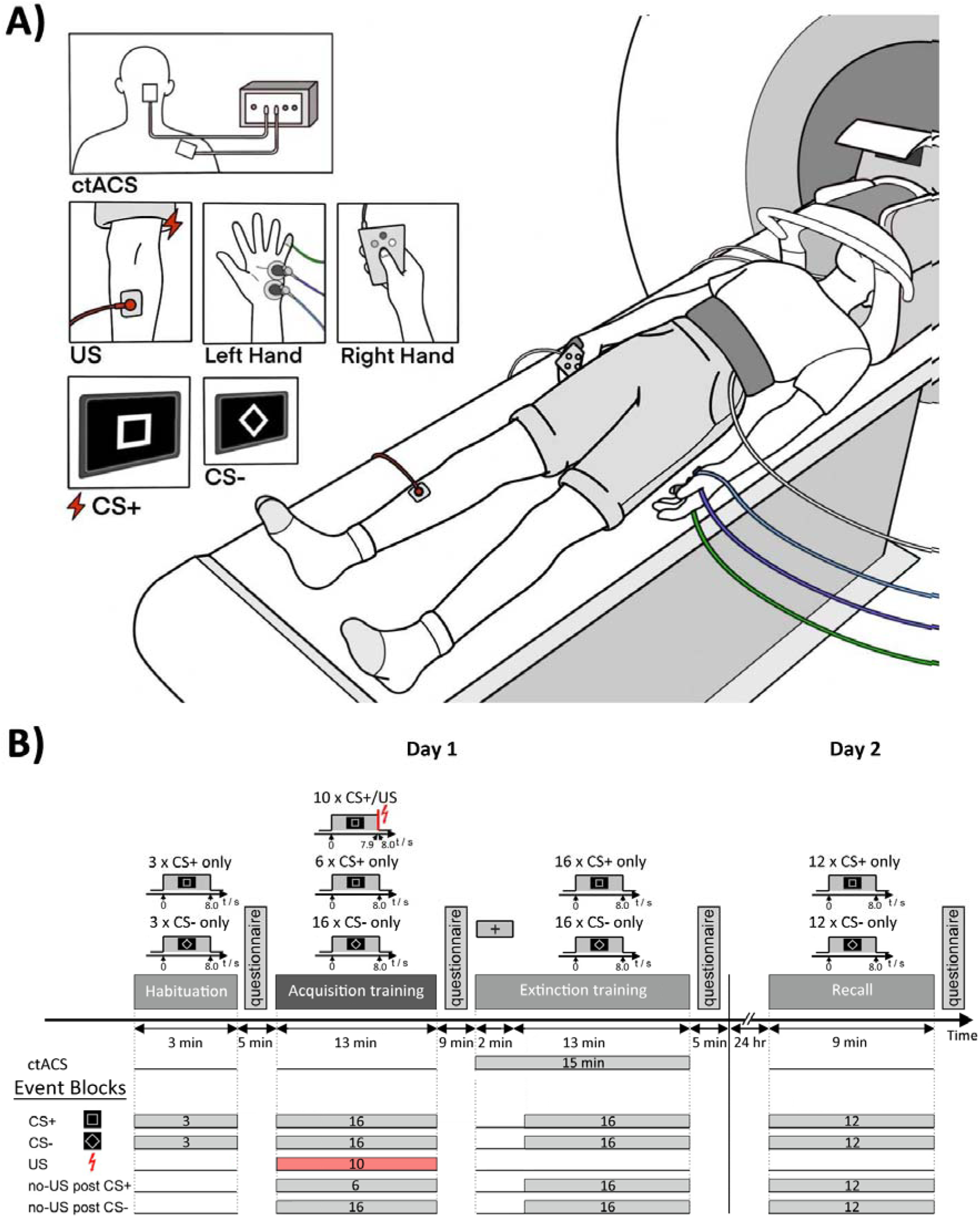
A) Experimental paradigm and **B)** event blocking scheme. During the experiment, two different geometrical figures, a diamond and a square, were presented as conditioned stimuli (CS), while an electric shock was used as an unconditioned stimulus (US). Additional information regarding the experiment can be found in the text.

The electric shock was generated by a constant current stimulator (DS7A, Digitimer Ltd., London, UK) and applied to the right shin via a concentric (ring-shaped) bipolar surface electrode with 6 mm conductive diameter and a central platinum pin (WASP electrode, Specialty Developments, Bexley, UK). The electrode position was marked with a permanent marker on day 1 to use the same electrode position on day 2. The 100 ms US consisted of a short train of four consecutive 500 µs current pulses (maximum output voltage: 400 V) with an interpulse interval of 33 ms. Before the start of MRI measurements, the US intensity for each participant was determined. The stimulation intensity was increased gradually, and participants were asked to rate the perceived sensation on a nine-point Likert scale ranging from "not unpleasant" to "very unpleasant" until a score of 8 out of 9 (i.e., “unpleasant but not painful”) was reached. To avoid a reduction in the conditioned responses caused by habituation to the US, the individual thresholds were increased by 20% (Batsikadze et al., 2022; Inoue et al., 2020). The average US intensity was 5.35 ± 5.87 mA (range 1 mA – 34.8 mA), and there were no significant differences in the applied current between the two groups (independent samples t-test: *t*_(38)_ = 0.714, *p* = 0.240).

Prior to each phase, the participants were provided with on-screen instructions, notifying them of the upcoming visual stimuli. Prior to habituation, they were informed that electric shocks would not be administered. Prior to the acquisition training, they were informed that electric shocks would be administered during this phase, and that if they detected a pattern between stimuli, it would remain consistent throughout the experiment. On the second day, participants were again reminded that any pattern perceived during day 1 would remain the same on day 2. All participants had to confirm that they read and comprehended the instructions.

Each trial consisted of an 8 s CS presentation. In the case of reinforced trials, a 100 ms aversive US was presented after 7.9 s and coterminated with the CS. A black cross image ("fixation cross") was displayed against a neutral gray background before the first CS picture onset and during the first two minutes of ctACS prior to the extinction phase. Intertrial intervals were randomized between 14.3 s and 17.9 s. Each experimental phase was performed within a separate session of fMRI data acquisition.

Participants had to answer a questionnaire following each phase of the experiment. As outlined in more detail in Batsikadze and colleagues (Batsikadze et al., 2022), they were asked to rate the hedonic valence, emotional arousal, fear, and expectancy of an US on a nine-step Likert scale. The scale ranged from "*very pleasant*" to "*very unpleasant*," "*very calm*" to "*very nervous*," "*not afraid*" to "*very afraid*," and "*US not expected* " to "*US expected*" respectively. After undergoing fear acquisition training, participants were asked to rate the unpleasantness of the US on a Likert scale ranging from 1 ("not unpleasant") to 9 ("very unpleasant") and to estimate the mean probability (in %) that a US occurred following the presentation of the CS (CS/US contingency).

Additionally, before and after the ctACS stimulation session, participants had to respond to questions about possible stimulation side effects (headache, neck pain, back pain, blurred vision, scalp irritation, scalp tingling, scalp itching, accelerated heartbeat, burning sensation, hot flashes, vertigo, sudden mood change, fatigue, phosphenes) and rate them on a Likert scale from 1 (“absent”) to 9 (“strong”) (adapted from Brunoni et al., 2011). Participants used an MRI-compatible button box with their right hand to give answers to questions that were projected onto the screen located inside the MRI scanner.

### Physiological data acquisition

Throughout the experiment, skin conductance responses (SCRs), pulse and breathing rate were acquired using MRI-compatible skin conductance, pulse oximetry and differential air pressure modules (MP160, BIOPAC Systems Inc., Goleta, CA). The sampling rate was set at 2 kHz. Two skin conductance electrodes were attached to the participant’s left hypothenar, approximately 20 mm apart. The pulse oximetry sensor was clipped to the participant’s left index finger. A respiratory belt was attached to the participant’s lower abdomen using a hook-and-loop belt.

### Modeling of prediction and prediction error values

An artificial agent was trained to predict the likelihood of a shock for a given visual input in a virtual version of the experiment as described in Batsikadze et al. (2022) and summarized in the following. The model was based on reinforcement learning (Sutton and Barto, 2018) and consisted of a deep neural network (DNN). The model hyper-parameters were fit to SCRs recorded in the experiment, which served as a read-out of the participants’ expectation of an US. The resulting model was used to derive predictions for the likelihood of a shock and the prediction error, which were then used as predictors in the fMRI data analysis.

Specifically, we used the same simplified visual stimuli as in Batsikadze et al. (2022). Reinforcement signals for paired and non-paired trials were coded as 1 and 0, respectively. The network architecture used to represent the agent’s value function comprised two hidden fully connected layers with 64 units each and an output layer to represent the probability of the shock. For each of the two trial sequences experienced by the participants, 25 randomly initialized agents were trained. On each trial *t*, the agent predicted the probability of the US *v_t_* and a prediction error Δ_*t*_, given *S*_*t*_ the current stimulus and a reinforcement signal. At the end of each trial, an experience tuple 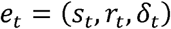 was stored in the agent’s memory for later replay (Lin, 1992). The agents were trained using the backpropagation algorithm (Rumelhart et al., 1986) on batches of experiences of size b, which were sampled randomly from memory with a probability that was proportional to a priority score :

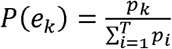

Priority scores depended on the experiences’ λ^𝜏^ recency where is the time passed since the experience and is a decay factor. Optionally, the priority could additionally depend on the magnitude of the US prediction error, i.e.,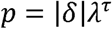(Schaul et al., 2015). The parameter RPE ∊ {Yes, No} indicates whether the replay priorities also depended on the magnitude of the US prediction error. The number of replays was varied to control for the degree of learning in each trial.

While the previous model could account for ABA renewal (Batsikadze et al., 2022), it cannot account for spontaneous recovery in the AAA paradigm, because, in the model, extinction in the same context overwrites the association formed during acquisition. Hence, we extended the model by an additional replay phase, which takes place between day 1 and day 2 trials and serves to recover the initially acquired association. The sleep replay phase prioritized experiences according to the reinforcement received in a trial and consisted of a total of 100 replays of batches of size 64. Reactivation probabilities for sleep replays were computed as follows:

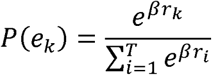

where denotes the inverse temperature and controls the relative difference of reactivation probabilities.

### Hyper-parameter fitting

We averaged SCRs from CS+ and CS- trials separately for each trial sequence. The averaged SCRs were accordingly defined as 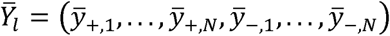 where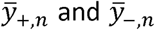 are the averaged SCRs for the n-th CS+ and n-th CS- presentations across all participants who completed a given trial sequence, respectively. Analogously, the averaged US predictions of the model were defined as 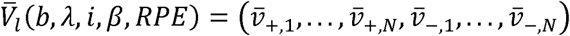 where 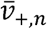 and 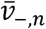 are the averaged US predictions for the n-th CS+ and CS- presentations across all model instances who were trained on a given trial sequence, respectively. The goodness of fit was defined as:

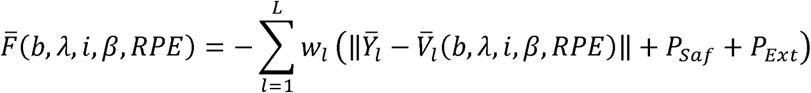

where is the number of participants who experienced trial sequence . To ensure that the overall learning curve in the model resembled that of the participants, we added the following penalty terms, if the model failed to

1. learn that CS- is not followed by the US: where is the average US prediction over the last 4 CS- presentations of acquisition training.
2. successfully extinguish the CR: where is the

average US prediction over the last 4 CS+ presentations of extinction training.

A grid search was conducted over the hyper-parameter sets shown in **Table 1**. The model with the best goodness-of-fit was then chosen to generate predictions that were used as parametric modulations in the fMRI data analysis.

**Table 1.** Modeling. Parameter values the grid search was run for.

| Grid Search Parameters |  |
| --- | --- |
| $b$ (batch size) | {1, 16, 32, 64, 96, 128} |
| $\lambda$ (decay factor) | {0.05, 0.4, 0.5, 0.6, 0.7, 0.8, 0.825, 0.85, 0.895, 1} |
| $i$ (training repeats) | {1, 2, 3, 4, 5, 6, 7, 8, 9, 10, 11, 12} |
| $\beta$ (inverse temperature) | {1, 2, 3, 4, 5} |
| $RPE$ | {Yes, No} |

### Cerebellar transcranial alternating current stimulation (ctACS)

Participants were randomly assigned to either a verum or a sham stimulation group and received 2 mA (peak-to-peak) 6 Hz ctACS for 15 minutes or sham ctACS for 30 seconds. The final data analysis included SCR data from 22 participants [10F/12M] in the verum group and 18 participants [8F/10M] in the sham group, and, for reasons given above, the final fMRI analysis included data from 19 participants [9F/10M] in the verum group and 16 participants [7F/9M] in the sham group. ctACS was started two minutes before the start of the extinction phase and lasted until its end. To ensure good contact with the scalp and provide a conductive medium, a thin layer of Ten20 paste (Ten20®, Weaver) was applied to each rubber electrode. To maintain consistency in experimental conditions, minimize stimulation-related potential discomfort and ensure effective blinding, a topical anesthetic cream containing 10% lidocaine (EMLA®, AstraZeneca, UK) was administered to the electrodes on top of the Ten20 paste approximately 10-15 minutes before participants entered the MRI scanner on both days of the study (McFadden et al., 2011). The target electrode (5 cm x 7 cm) was placed vertically over the right cerebellar cortex. The non-target electrode (5 cm x 7 cm) was placed over the right deltoid muscle. The current was ramped up and down for 15 s at the beginning and the end of the stimulation. The battery-driven MRI-compatible DC-Stimulator Plus (neuroConn GmbH, Ilmenau, Germany) was used to deliver the stimulation. Both the experimenter and the participants were blinded to the type of stimulation. The study achieved double blinding by utilizing the stimulator’s study mode, where pre-assigned five-digit codes were inputted into the device to initiate either the active or sham protocol.

We found in a prior study that cerebellar tDCS effects on neurophysiological measures (i.e., cerebellar-brain inhibition, CBI) were not significantly different using the more standard position of the return electrode on the buccinator muscle compared to the less commonly used position on the deltoid muscle (Batsikadze et al., 2019). Thus, the deltoid muscle was used as position for the return electrode, which is preferred in the MR scanner as compared to the face reducing the risk of possible side effects. In addition, computational modeling of transcranial electric stimulation (tES) induced electric fields was performed to test which cerebellar electrode positions result in the most selective stimulation of the cerebellar target areas. Target areas were determined based on fMRI data from Ernst et al. (2019). Simulations were performed using SimNIBS (Thielscher et al., 2015) to choose a montage which maximizes the average electric field (EF) value in the right Crus I with the least possible average EF values in the vermis, left crus I, and extracerebellar areas involved in the fear network including R/L DLPFC, hippocampus, and amygdala. This resulted in a montage with the cerebellar electrode centered on PO10 (according to the 10-20 EEG system, (Homan, 1988)) with the return electrode over the neck – as a proxy for the deltoid muscle (**Figure 2**). The cerebellar electrode was placed accordingly. Note that although electrode position was optimized for stimulation of Crus I, maximum stimulation is still in Crus II, which is a characteristic finding in cerebellar tDCS (Batsikadze et al., 2019; D’Mello et al., 2017).

**Figure 2.**
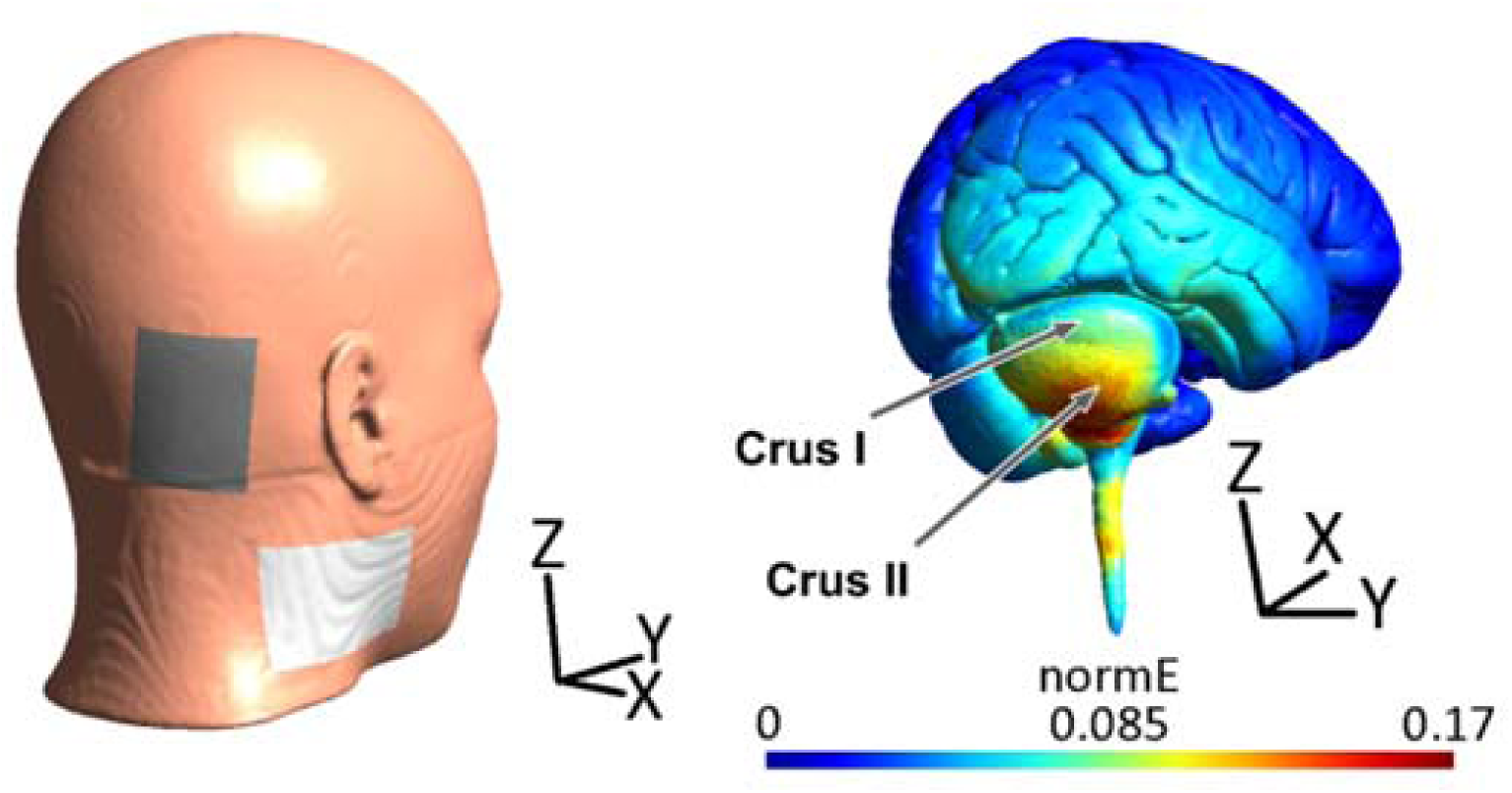
Transcranial electric stimulation (tES) montage resulting in most selective stimulation of target area, the right Crus I (neck position corresponds to shoulder position). normE – the magnitude of stimulation-induced electric field.

### MRI acquisition

All MR images were acquired with the participants lying headfirst supine inside a clinical 3 T scanner (VIDA, Siemens Healthineers, Erlangen, Germany) equipped with a 1-channel transmit/ 64-channel receive array head coil. Electrodes for ctACS were attached to the participant’s head before placement in the scanner and fixed in position with ample bandages. As needed, inflatable cushions were used to fix the head position within the coil. To minimize inter-subject placement variance, the auto-alignment option was used, and manually corrected if needed.

Isotropic 1 mm resolution anatomic T1-weighted images were acquired using the MPRAGE sequence immediately before functional MRI acquisition on day 1. Sequence parameters were selected as follows: TR/TE, 2530/2.27 ms, TI, 1100 ms, flip angle, 7°, parallel acceleration factor, 2, acquisition matrix, 256 × 256, number of slices, 176, TA, 6:03 min. Additionally, isotropic 1-mm resolution MP2RAGE was acquired after fMRI acquisition on day 1. Sequence parameters were selected as follows: TR/TE, 5000/2.98 ms, TI1/TI2, 700/2500 ms, flip angles 1/2, 4°/5°, parallel acceleration factor, 3, acquisition matrix, 256 × 256, number of slices, 176, TA, 7:19 min. No anatomic images were acquired on day 2.

Functional MRI acquisition was performed to cover the whole brain with an isotropic voxel size of 2.5 mm^3^ using a fat-saturated, two-dimensional simultaneous multi slice echo planar image (SMS-EPI) sequence. Further imaging parameters were selected as follows: TR/TE, 1790/30.0 ms, flip angle, 60°, parallel acceleration factor (GRAPPA), 2, SMS factor, 2, acquisition matrix, 96 × 96, number of slices, 60, bandwidth, 2,170 Hz/pixel, phase encoding direction anterior to posterior. To correct for distortion artifacts before functional acquisition, two brief EPI sequences were acquired: (a) five frames of opposed-phase reference, i.e., with opposite phase encoding direction (posterior to anterior), and (b) one frame as single band reference, i.e., without SMS acceleration, and otherwise identical parameter settings as the actual fMRI sequence.

Additionally, resting state and diffusion images were acquired immediately after fMRI acquisition on day 2. However, resting state and diffusion data is not presented in this work.

### Image preprocessing

DICOM data was converted to NIFTI format using dcm2niix software (version 1.0.20201102; Li et al., 2016) and all data was structured to comply with brain imaging data structure (BIDS; Poldrack et al., 2024). All evaluation steps were then performed on a laptop running 64-bit Linux (distribution Ubuntu 22.04). Data preprocessing was carried out using the fMRIPrep Docker container (version 23.1.4), a robust preprocessing pipeline that standardizes functional MRI preprocessing (Esteban et al., 2019), with default settings and with Freesurfer surface processing disabled (--fs-no-reconall). The preprocessing pipeline involved several key steps, including skull stripping, head motion correction, susceptibility distortion correction, and spatial normalization to the MNI152NLin2009cAsym template with an isotropic resolution of 2 mm. Please refer to supplements for further details.

Finally, the functional BOLD volume images were smoothed using a smoothing kernel of 6 mm, and voxel timeseries were low-pass filtered with a Gaussian-weighted least-squares straight line fitting, with sigma = 2.548, as implemented in FSL Feat.

### Analysis and statistics

#### Skin conductance responses

Skin conductance data was low-pass filtered with a 10 Hz cutoff using a hardware filter (EDA100C-MRI module, BIOPAC Systems Inc., Goleta, CA). Offline data processing was performed using the MATLAB-based (Release 2019a, RRID: SCR_001622, The MathWorks) software EDA-Analysis App (Otto et al., 2023). SCRs were identified as the highest peaks that met specific requirements, including a minimum amplitude of 0.01 μS and a minimum rise time of 500 ms (Boucsein et al., 2012), starting within a time window of 1 to 6 s after the onset of the CS. Trials not meeting the criteria were scored as zero and included in subsequent analysis (Pineles et al., 2009).

The resulting raw SCRs were averaged into blocks and normalized through a logarithmic [LN(1+SCR)] transformation (Boucsein, 2012; Venables and Christie, 1980). Three habituation trials of the same CS were combined to form single blocks. In the subsequent phases, the trials of the same CS were divided into early and late blocks. Specifically, in the acquisition and extinction training, the averaging included the first and last six trials, while in the recall, the averaging included the first and last three trials. The Shapiro-Wilk-test was used to test the normalized data and the distribution of residuals for normality. Since the normality test revealed a non-normal distribution of SCRs and the residuals (*p* < 0.05), data were analyzed with non-parametric statistical analysis for repeated measures using rank-based F-tests (ANOVAF option in the PROC MIXED method in SAS, SAS Studio 3.8, SAS Institute Inc, Cary, NC, USA) and the nparLD R package (http://www.R-project.org/), which is recommended for dealing with skewed distributions, outliers or small sample sizes (Brunner et al., 2002). These methods use an ANOVA-type statistic with the denominator degrees of freedom set to infinity (Brunner et al., 2002; Noguchi et al., 2012) to enhance the reliability of the ANOVA-type statistic. Using finite denominator degrees of freedom can lead to increased type I errors (Bathke et al., 2009).

Non-parametric ANOVA-type statistics for repeated measures (ATS) were used separately for each phase with SCRs as dependent variable and stimulus (CS+, CS-) and block (early, late) as within-subjects factors and group (verum, sham) as between-subjects factors as well as their interactions. In addition, late acquisition training and early extinction training were compared. In case of significant results of ATS, *post hoc* comparisons were performed using least square means tests and were adjusted for multiple comparisons using the Tukey-Kramer method.

To address the individual variability of SCR amplitudes, we computed the differential skin conductance responses (SCR_diff_) by subtracting the SCR to CS- from the SCR to CS+ for each block (Schellen et al., 2023). This enabled us to quantify the differential reaction to the conditioned stimuli. ATS was used separately for each phase, with SCR_diff_ as the dependent variable, block (early, late) as the within-subjects factor, and group (verum, sham) as the between-subject factor, in addition to their interactions.

To quantify the effect sizes, we used a metric called relative treatment effects (RTE). The RTE represents the probability that a randomly selected value from one specific factor level of interest (*X*) is greater than, less than, or equal to the mean value (*Y*) of a fixed reference distribution, expressed as *px = P(X > Y*,*) px = P(X < Y*,) or *px = P(X = Y*.) If *p_X_* is lower than *p_Z_*, it suggests that measurements taken under condition *X* are generally smaller than those under condition *Z*. Conversely, *p_X_*= *p_Z_* indicates no consistent difference between the data from conditions *X* and *Z*. For example, a *p_X_* value of 0.25 means there is approximately a 25% chance of randomly selecting a participant from the dataset who would score lower than a randomly chosen participant from condition *X* (Rubarth et al., 2022).

#### Questionnaires

Questionnaires were analyzed using ATS with the respective rating as dependent variable and stimulus (CS+, CS-) and time (prior to, post fear acquisition training, post extinction training and post recall) as within-subjects factor and group (verum, sham) as between-subjects factor as well as their interactions.

Side effect ratings were analyzed using ATS with the respective rating as dependent variable, time (prior and post ctACS) as within-subjects factor and group (verum, sham) as between-subjects factor as well as their interaction.

#### fMRI analysis

The first-level analysis was performed on the preprocessed functional data, independently for all runs using FSL FEAT (FMRI Expert Analysis Tool) Version 6.0.1, part of FSL (FMRIB’s Software Library, www.fmrib.ox.ac.uk/fsl). Feat routines were invoked from Python using Nipype (version 1.8.6). In-scanner motion parameters estimated by fMRIprep (i.e., 3 rotations and 3 translations) were used as first-level nuisance regressors.

The first-level analysis was modeled as an event related-design for the entire experiment, i.e., all event durations were set to 0 s. Onsets of presentations of the CS+, CS-, and US (including the corresponding point in time for unpaired trials, i.e., US omission after CS presentation, further referred to as no-US) were modeled as individual events. Individual events were blocked as shown in **Figure 1**. First level main effect contrasts against baseline and appropriate differential first level contrasts were generated.

Finally, a second separate first level analysis was performed on the preprocessed and smoothed functional data. For each experimental phase, all events for each event type (CS, US and no-US) were grouped irrespective of CS trial type (CS+/CS-). Trial-by-trial parameters derived from the learning model were applied as parametric modulations, i.e., the mean prediction parameters for the CS events and the mean absolute prediction error parameters for US and no-US events.

For second level analysis, first-level contrasts were tested with a fixed-effect analysis within, across and between groups. Cluster-wise correction for multiple comparisons was applied using a Z-threshold of 3.1 and a cluster significance threshold of *p* < 0.05 (Worsley, 2001). The anatomical brain locations of significant clusters were determined using the Harvard-Oxford cortical and subcortical structural atlases (https://identifiers.org/neurovault.collection:262) and Cerebellum atlas (the version normalized with FLIRT). When the Harvard-Oxford atlas was insufficient for identification of subcortical structures, manual identification was conducted using the multi-contrast anatomical subcortical structures (MASSP) atlas (Bazin et al., 2020). The correspondence between anatomical and functional areas was verified by author G.B.. To display results, cerebellar (SUIT space) activation maps were plotted on cerebellar flatmaps (Diedrichsen, 2006; Diedrichsen and Zotow, 2015).

## Results

### Skin conductance responses (SCRs)

*<u>Habituation phase (day 1):</u>* There was no significant difference in the mean SCR amplitudes between the CS+ and CS-. Additionally, there was no group difference in differential SCRs (SCR_diff_). ATS did not reveal any significant effects or interactions (all *p* values ≥ 0.108, **Table 3- 1**).

*<u>Acquisition training (day 1):</u>* Both groups of participants showed higher SCRs in response to the CS+ than to the CS-, and the responses were larger during the early block compared to the late block. The differences between the CS+ and CS- responses were significant for both groups (**Figure 3**). ATS revealed significant main effects of Block (*F*_1_ = 57.62, *p* < 0.001), Stimulus type (*F*_1_ = 56.75, *p* < 0.001) and a significant Block x Stimulus type interaction (*F*_1_ = 4.69, *p* = 0.03; **Table 3-1**). No significant group differences and interactions were revealed (all *p* values ≥ 0.117). *Post hoc* exploratory analysis of the Block × Stimulus differences revealed significantly higher SCRs in the early vs. late acquisition blocks in both groups towards both CSs, and significantly higher SCRs towards CS+ vs. CS- in both early and late blocks (all *p* values < 0.001, least squares means test).

**Figure 3.**
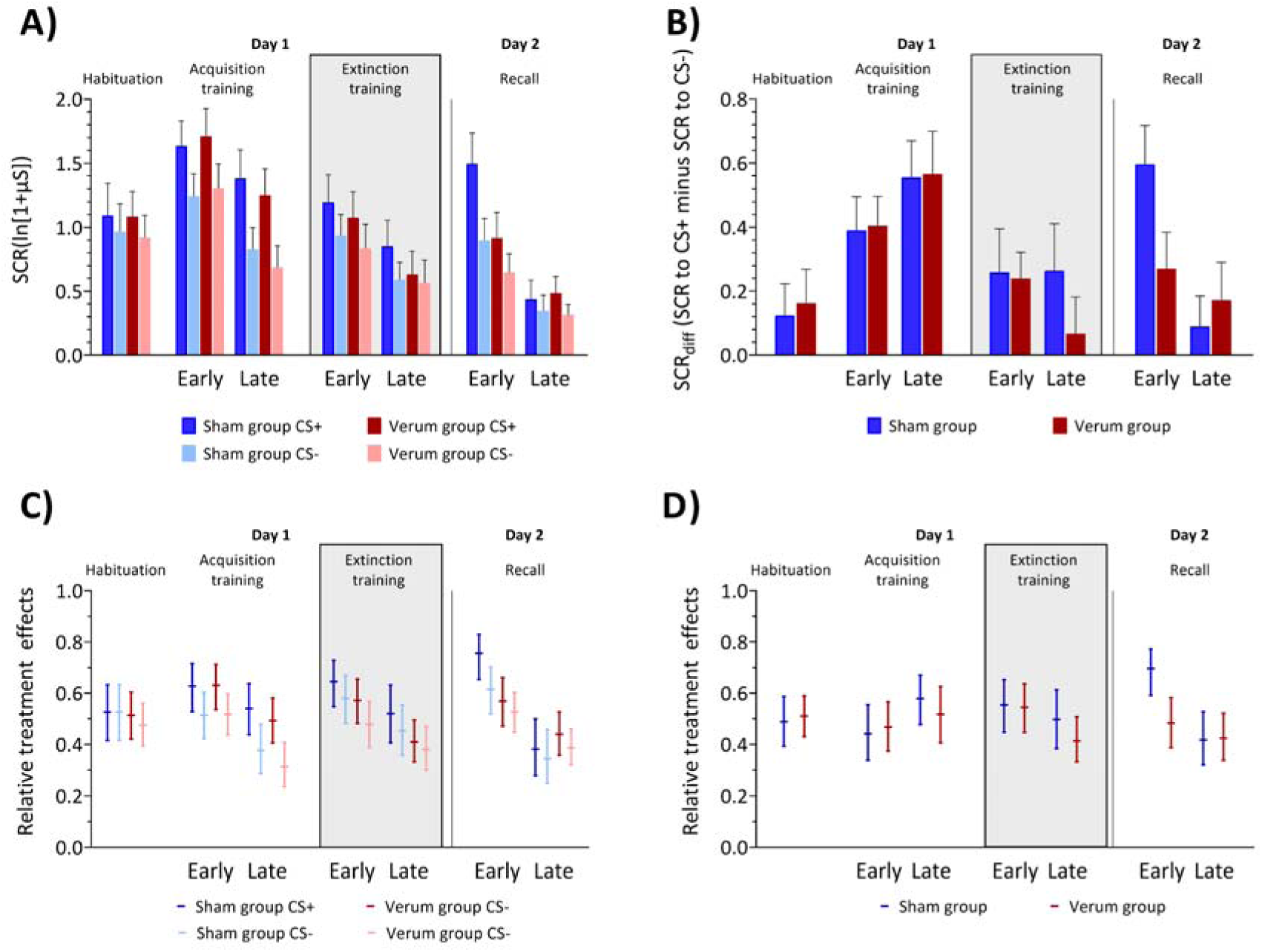
A) Skin conductance response (SCR) amplitudes, **B)** differential skin conductance responses (SCR_diff_) and **C, D)** respective relative treatment effect (RTE) estimates during fear acquisition training, extinction training and recall. **A,B)** Colored bars represent group mean (log-transformed) values for early and late blocks of fear acquisition training, extinction training and recall. Error bars indicate S.E.M. **C,D)** Horizontal lines denote median RTEs and whiskers denote 95% confidence intervals. Blue colors = sham group, red colors = verum group. Dark colors: CS+, light colors: CS-. The grey background in the figure represents the time during which transcranial alternating current stimulation (ctACS) was administered.

The SCR_diff_ analysis did not show any significant effects of Block, Group, or their interaction (all *p* values ≥ 0.066).

*<u>Extinction training (day 1)</u>* : Both groups showed a difference of SCR amplitudes between the CS+ and CS-, with higher SCRs elicited by the CS+. Larger SCRs were observed in the early block compared to the late block (**Figure 3**). ATS revealed a significant main effect of Block (*F*_1_ = 32.25, *p* < 0.001) and Stimulus type (*F*_1_ = 12.93, *p* < 0.001). No significant group differences and interactions were revealed (all *p* values ≥ 0.247; **Table 3-1**). Exploratory ATS between late acquisition training and early extinction training blocks showed a significant main effect of CS (*F*_1_ = 43.24, *p* < 0.001) and CS x Block interaction (*F*_1_ = 8.11, *p* = 0.004, **Table 3-1**). Pairwise comparisons revealed significant differences between CS+ and CS- both in late acquisition training and early extinction training blocks (both *p* values ≤ 0.003), but not between phases (both *p* values ≥ 0.305).

The SCR_diff_ analysis showed a close to significant Block effect *(p* = 0.056). The group effect was not significant (*p* = 0.559) and there was no significant Block x Group interaction (*p* = 0.445; **Table 3-1**). Exploratory analysis between late acquisition and early extinction training blocks showed that SCR_diff_ was significantly larger in late acquisition training compared to early extinction training. ATS revealed a significant main effect of Block (late acquisition training vs early extinction training: *F*_1_ = 8.97, *p* = 0.003, **Table 3-1**), but no significant main effect of Group (*p* = 0.583) or Group x Block interaction (*p* = 0.654) were revealed.

Visual inspection of **Fig. 3A** and **3B** suggest that, in late extinction, extinction effects were stronger in the verum group compared to the sham group. This was, however, not reflected in the statistical analysis (that is, no significant Group effects, and none of the interactions involving Group coming even close to significance). Exploratory analysis of the late extinction block showed no significant Group effect (*p* = 0.233) and no significant Stimulus x Group interaction (*p* = 0.668). For differential SCRs, the Group effect in late extinction was also not significant (*p* = 0.433).

*<u>Recall (day 2):</u>* In both groups, the SCRs were higher in the early compared to the late block. In the early recall phase, the SCRs related to the CS+ were larger than to the CS- in the sham group, but this was not observed in the verum group (**Figure 3**). In the late recall phase, both the sham and verum groups responded to both the CS+ and CS- in a similar manner. ATS revealed significant main effects of Block (*F*_1_ = 74.03, *p* < 0.001), Stimulus type (*F*_1_ = 10.02, *p* = 0.001) and a significant Block x Group (*F*_1_ = 12.59, *p* < 0.001) interaction. No other significant main effects or interactions were revealed (all *p* values ≥ 0.179; **Table 3-1**). The significant interaction between Block and Group was driven by a disordinal interaction, with the SCRs in the early sham block being larger than those in the late verum block (*p* = 0.003, least square means test).

Exploratory *post-hoc* ATS on the first recall block revealed a significant main effect of Stimulus type (*F*_1_ = 20.78, *p* < 0.001) and Group x Stimulus type interaction (*F*_1_ = 4.27, *p* = 0.039). The effect of Group was not significant (*F*_1_ = 3.00, *p* = 0.083). The sham group showed significantly higher SCRs related to the CS+ compared to the CS- in pairwise comparisons (least square means test, *p* < 0.001), while the verum group did not (least square means test, *p* = 0.443; **Figure 3**).

Regarding SCR_diff_, ATS revealed a significant main effect of Block (F_1_ = 11.30, *p* < 0.001) and a significant interaction between Block and Group (F_1_ = 4.82, *p* = 0.028; **Table 3-1**). SCR_diff_ values were significantly higher in the early block compared to the late block (*p* = 0.002, least square means test), however, this effect (early vs. late block) was particularly present in the sham (*p* = 0.001, least square means test), but not in the verum group (p = 0.853, least squares means test). Additionally, exploratory ATS on the first recall block showed a significant main effect of Group (F_1_ = 5.36, *p* = 0.021), with the sham group exhibiting significantly higher differential SCRs.

### Questionnaires

*US unpleasantness, CS-US contingency* . Median US unpleasantness was rated 7 (interquartile range, IQR 6-7) post acquisition on a Likert scale ranging from 1 (“not unpleasant”) to 9 (“very unpleasant”). No significant group differences occurred between reported mean US unpleasantness (Mann-Whitney U test, *z* = 0.041, *p* = 0.968). The mean reported probability of an US occurring after the presentation of the CS+ was estimated to be 63.6% ± 19.5%, while the mean reported probability of an US occurring after the presentation of the CS- was 2.5% ± 7.2% (0% probability by 36 out of 40 [95%] participants). There were no significant differences between the reported mean probabilities in the two groups (Mann-Whitney U tests, all *p* values ≥ 0.222).

Before acquisition training, there were no significant differences in valence, arousal, and fear ratings between the CS+ and CS-. After acquisition training, the CS+ was rated as significantly less pleasant, with higher arousal and fear compared to the CS-, and these differences persisted through extinction training and recall. Similarly, before acquisition training, participants had similar expectation for the US after both stimuli, but after acquisition training, US expectation was significantly higher after the CS+ than the CS-, and this difference also persisted until the end of the experiment. The detailed results of the questionnaire analysis are presented in Extended data **Figure 3-1**, **Tables 3-1 and 3-2.**

*<u>Stimulation side effects.</u>* The verum group reported significantly higher scalp tingling. ATS revealed a significant main effect of Group (*F*_1_ = 4.22, *p* = 0.040), but not Time (*p* = 0.286) and Group x Time interaction (*p* = 0.665). However, it is worth noting that this significant effect was driven by one participant in the verum group who reported strong sensations (skin irritation, skin itching, skin tingling) both before and after stimulation, which could be attributed to discomfort caused by the electrode and skin contact in this individual. When this participant was excluded from the analysis, the significant group effect was no longer observed (*p* = 0.332). Other stimulation side effect ratings were not significantly different between groups (all *p* values ≥ 0.284). Both groups reported significantly higher post-ctACS increased headache (*p* = 0.012), fatigue (*p* = 0.013) and a close-to-significantly increased scalp irritation (*p* = 0.058) compared to pre-ctACS. Responses to the items "increased heartbeat" (*p* = 0.020) and “sudden mood change” (*p* = 0.005) were significantly lower post-stimulation compared to the pre-stimulation values. Other ratings were not significant (all *p* values ≥ 0.107; **Table 3-1**). The questionnaire results suggest that blinding was successful, as the ratings of sensations reported by participants who received verum stimulation were not significantly different from those who received sham (**Table 3-2**).

### fMRI data

As stated above, the anatomical brain locations were determined using the Harvard-Oxford atlas for cortical and subcortical structures (https://identifiers.org/neurovault.collection:262), the Cerebellum atlas (the version normalized with FLIRT) and the MASSP atlas (Bazin et al., 2020). Corresponding functional brain subregions are provided in square brackets.

### Activations during fear acquisition training

Because tACS was administered during extinction training, no significant differences were expected between the sham and verum group during acquisition training. As anticipated, activation patterns showed similar results for each group when analyzed separately (see Supplements). No significant between-group differences [contrasts ’verum > sham’ and ’verum < sham’] were found (**Table 4-1** and **Figure 4**), therefore findings are reported by combining participants from both groups.

**Figure 4.**
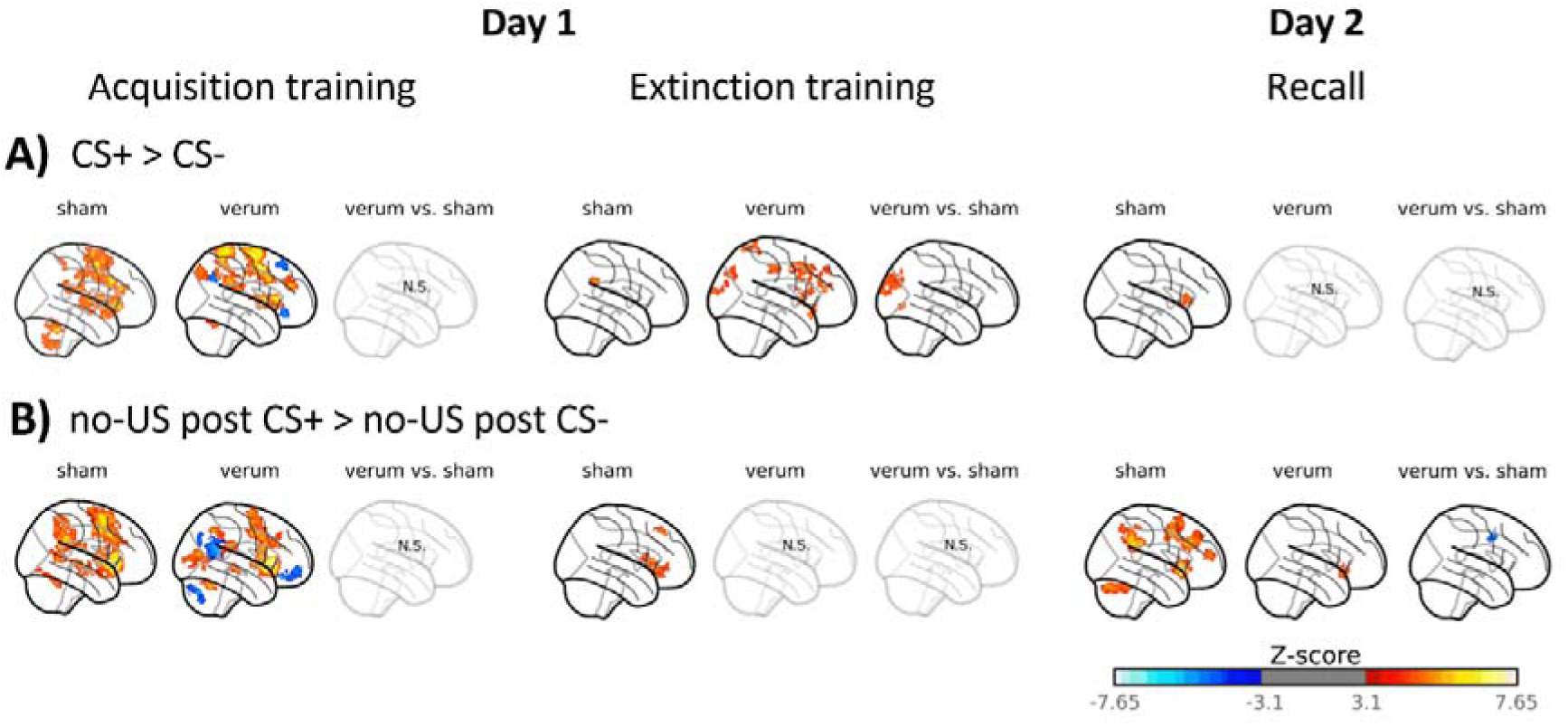
The glass brain visualization showing cerebral and cerebellar activations during fear acquisition training, extinction training and recall related to the **A)** prediction of the CS (contrast ‘CS+ > CS-’) and **B)** omission of the aversive US (contrast ‘no-US post CS+ > no-US post CS-’) provided separately for verum and sham groups alongside a differential contrast (contrast ‘verum vs sham’). In the individual group contrast increases are shown in red and decreases in blue. In the differential contrasts ‘verum vs sham’, shades of blue denote higher activation in the sham group, while shades of red denote higher activation in the verum group. CS = conditioned stimulus; US – unconditioned stimulus; N.S. – no significant clusters. Results of fMRI analysis are provided in **Tables 4-1**, **4-2** and **4-3.**

*<u>Activation related to the prediction of the aversive stimulus [contrast ‘CS+ > CS-’]</u> .* Significantly higher brain activity was observed related to the CS+ compared to the CS- in brain areas well known to show increased differential activations in fear acquisition training (e.g., Fullana et al., 2016). These areas included the insular cortex, anterior cingulate cortex (ACC), frontal cortical areas (middle frontal gyrus [corresponds to dlPFC], frontal pole, precentral gyrus [corresp. primary motor cortex – M1], and supplementary motor area - SMA), parietal cortical areas (postcentral gyrus [corresp. primary somatosensory cortex – S1], anterior and posterior supramarginal gyri [corresp. secondary somatosensory cortex – S2], parietal operculum, precuneal cortex, and superior parietal lobule), occipital cortical areas (lingual gyrus), as well as subcortical areas (basal ganglia, thalamus), midbrain (red nucleus) and the cerebellum (bilaterally in cerebellar lobules VI (including vermis) and Crus I, as well as VII and VIII) (**Table S3**, **Figures 2** and **S2**, shades of red). The contrast CS- > CS+ revealed no significant differences, that is, no brain areas showed significantly higher activations towards the CS-.

*<u>Activation related to the omission of the aversive stimulus [contrast ‘no-US post CS+ > no US post CS-’].</u>* The pattern of differential activations related to the (unexpected) omission of the US showed overlap with areas related to the prediction of the US. Overall, activated cortical areas were more extended, and in addition areas within the occipital lobe showed increased activations likely because attention is redirected to the CS+. Areas included the insular cortex, anterior cingulate cortex (ACC), frontal cortical areas (middle frontal gyrus [corresp. dlPFC], inferior frontal gyrus [corresp. ventrolateral prefrontal cortex – vlPFC], frontal operculum and orbitofrontal cortex), parietal cortical areas (postcentral gyrus [corresp. S1], anterior and posterior supramarginal gyri [corresp. S2], angular gyrus, parietal operculum and superior parietal lobule) and occipital cortical areas (lingual and occipital fusiform gyri), subcortical areas (basal ganglia, thalamus, VTA), and left cerebellar lobules VI-VII, left Crus I and II, right VI, and right Crus I (**Table S3**, **Figures 2** and **S2**, shades of red).

Higher activations towards the omission of the CS- [contrast ’no-US post CS+ < no-US post CS- ’] were observed in the posterior cingulate cortex (PCC), precuneal cortex, frontal cortical areas (frontal pole [corresp. dlPFC], frontal medial cortex [corresp. ventromedial prefrontal cortex - vmPFC], precentral gyrus [corresp. M1]), postcentral gyrus [corresp. S1], anterior and posterior divisions of the middle temporal gyrus, superior division of lateral occipital cortex, left amygdala, left hippocampus as well as the right cerebellar Crus I and II (**Table 4-1**, **Figures 2** and **4-1**, shades of blue).

*Activations during fear extinction training*.

*<u>Activation related to the prediction of the aversive stimulus [contrast ‘CS+ > CS-’]</u> .* In the sham group, activations were observed in the supramarginal gyrus of the parietal lobe [corresp. S2] (**Table 4-2**, **Figure 4**). In the verum group, activations during extinction training were more widespread. Significant activations were detected in insular cortex, cingulate cortex (ACC and PCC), frontal cortical areas (inferior frontal gyrus [corresp. vlPFC], orbitofrontal cortex, frontal operculum, precentral gyrus [corresp. M1], and SMA), occipital cortical areas (superior division of the lateral occipital cortex and occipital pole) and superior parietal lobule (**Table 4-2**, **Figure 4**, shades of red).

Between-group comparisons [contrasts ’verum > sham’ and ’verum < sham’] revealed significantly higher activation in the cuneal cortex, occipital pole and lingual gyrus in the verum compared to the sham group (**Table 4-2**, **Figure 4**, shades of red).

*<u>Activation related to the omission of the aversive stimulus [contrast ‘no-US post CS+ > no US post CS-’]</u>.* During fear extinction training, activations were observed in the sham group in the frontal cortical areas (inferior frontal gyrus [corresp. vlPFC], superior frontal gyrus [corresp. dlPFC], and orbitofrontal cortex) (**Table 4-2**, **Figure 4**, shades of red). No significant activations were observed in the verum group. Between-group differences were not significant (contrasts ’verum > sham’ and ’verum < sham’).

### Activations during recall

*<u>Activation related to the prediction of the aversive stimulus [contrast ‘CS+ > CS-’]</u> .* In the sham group, activations were observed in the insular cortex and frontal operculum (**Table 4-3**, **Figures 4** and **4- 2**, shades of red). No significant activations were observed in the verum group. Between-group differences were not significant [contrasts ’verum > sham’ and ’verum < sham’].

*Activation related to the omission of the aversive stimulus [contrast ‘no-US post CS+ > no US post CS-’].* During recall, widespread activations (shades of red) were observed in the sham group, including the insular cortex, cingulate cortex (ACC and PCC), frontal cortical areas (inferior frontal gyrus [corresp. vlPFC], middle frontal gyrus [corresp. dlPFC], frontal pole, orbitofrontal cortex and frontal operculum), parietal cortical areas (supramarginal gyrus [corresp. S2] and parietal operculum), occipital cortex (cuneal cortex, occipital pole, lingual gyrus, superior division of lateral occipital cortex), basal ganglia, as well as bilaterally in cerebellar lobules VI and Crus I (**Table 4-3**, **Figures 4** and **4-2**, shades of red).

In the verum group, activations were detected in the insular cortex and frontal cortical areas (inferior frontal gyrus [corresp. vlPFC] and orbitofrontal cortex) (**Table 4-3**, **Figures 4** and **4-2**, shades of red).

Between-group comparisons [contrasts ’verum > sham’ and ’verum < sham’] revealed significantly higher activation in M1 [precentral gyrus] in the sham compared to the verum group (**Table 4-3**, **Figures 4** and **4-2**, shades of red).

### Parametric modulation with model predictions for shock probability and prediction errors

Parametric modulation was used to test whether learning-model derived predictions and prediction errors were associated with fMRI signal alterations during CS and no-US events. CS events denote the onset of conditioned stimuli, while no-US events represent instances where the US was anticipated in reinforced CS+ trials but was absent in unreinforced CS+ and CS- trials. Thus, the modulation is determined by trial-by-trial model-derived predictions for CS events and prediction errors for no-US events.

### Fear acquisition training

No significant between-group differences for parametric modulation effects for CS and no-US events [contrasts ’verum > sham’ and ’verum < sham’] were found (**Table 5-1** and **Figure 5**). Therefore, findings are reported by combining participants from both groups.

**Figure 5.**
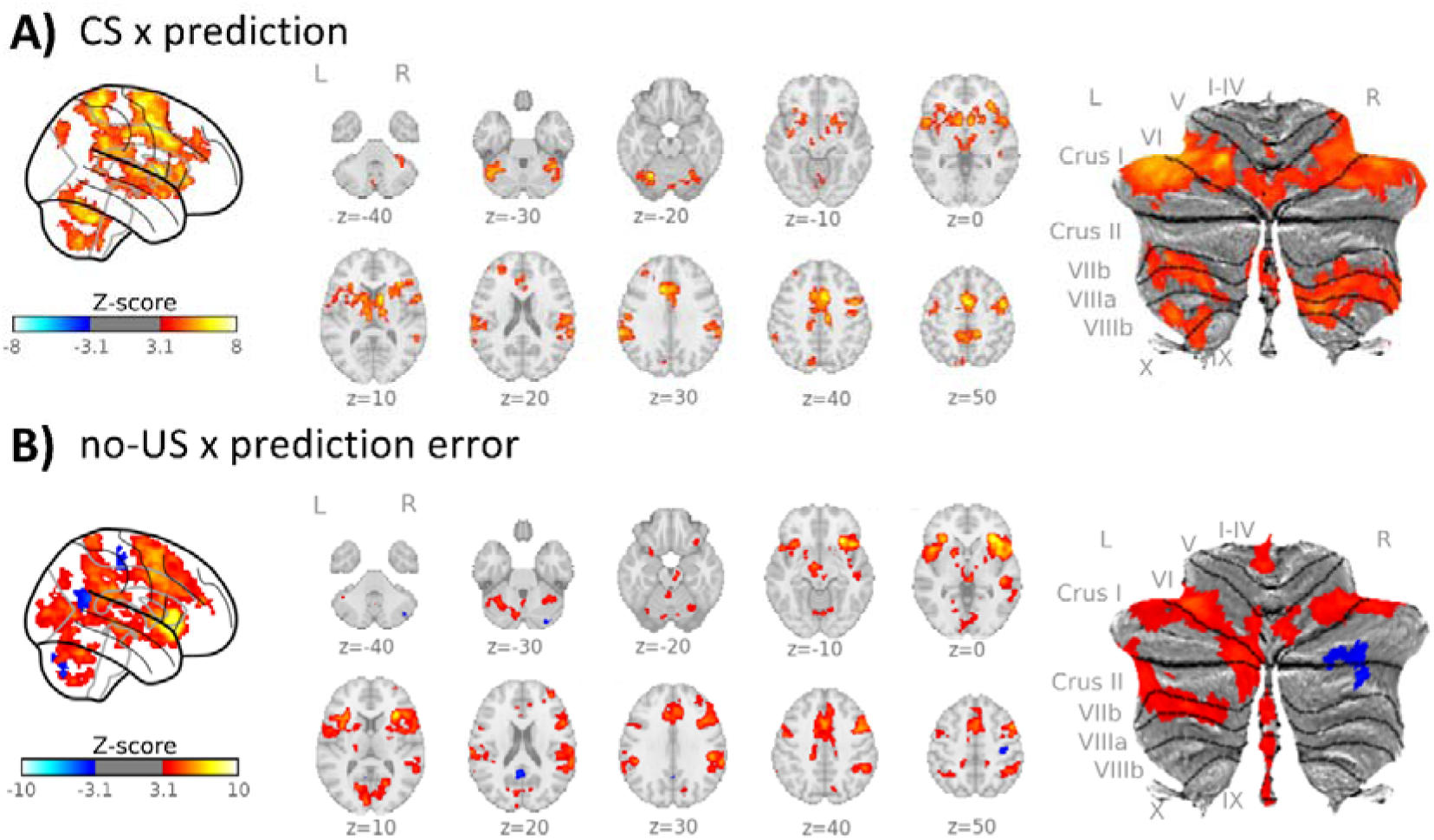
Parametric modulation with learning model-derived data during fear acquisition training in both verum and sham groups. **A)** Parametric modulation of CS events with individual mean prediction values (CS x prediction), **B**) parametric modulation of omission of US events at CS termination with individual absolute mean prediction error values (no-US x prediction error). Increases are shown in red, decreases in blue. CS = conditioned stimulus; L = left; R = right; SUIT = spatially unbiased atlas template of the cerebellum; results of fMRI analysis are provided in **Table 5-1.**

During fear acquisition training, parametric modulation effects for CS and prediction values revealed significant clusters in the insular cortex, ACC, frontal cortical areas (middle frontal gyrus [corresp. dlPFC], inferior frontal gyrus [corresp. vlPFC], orbitofrontal cortex, precentral gyrus [corresp. M1] and SMA), parietal cortical areas (postcentral gyrus [corresp. S1], anterior and posterior supramarginal gyri [corresp. S2], precuneal cortex, angular gyrus and superior parietal lobule), occipital cortical areas (lateral occipital cortex), temporal cortical areas (middle and superior temporal gyri, lingual gyrus, planum temporale), subcortical areas (basal ganglia, thalamus) and the cerebellum (bilaterally in lobules VI-IX and Crus I-II; **Table 5-1**, **Figures 5** and **6**, shades of red).

**Figure 6.**
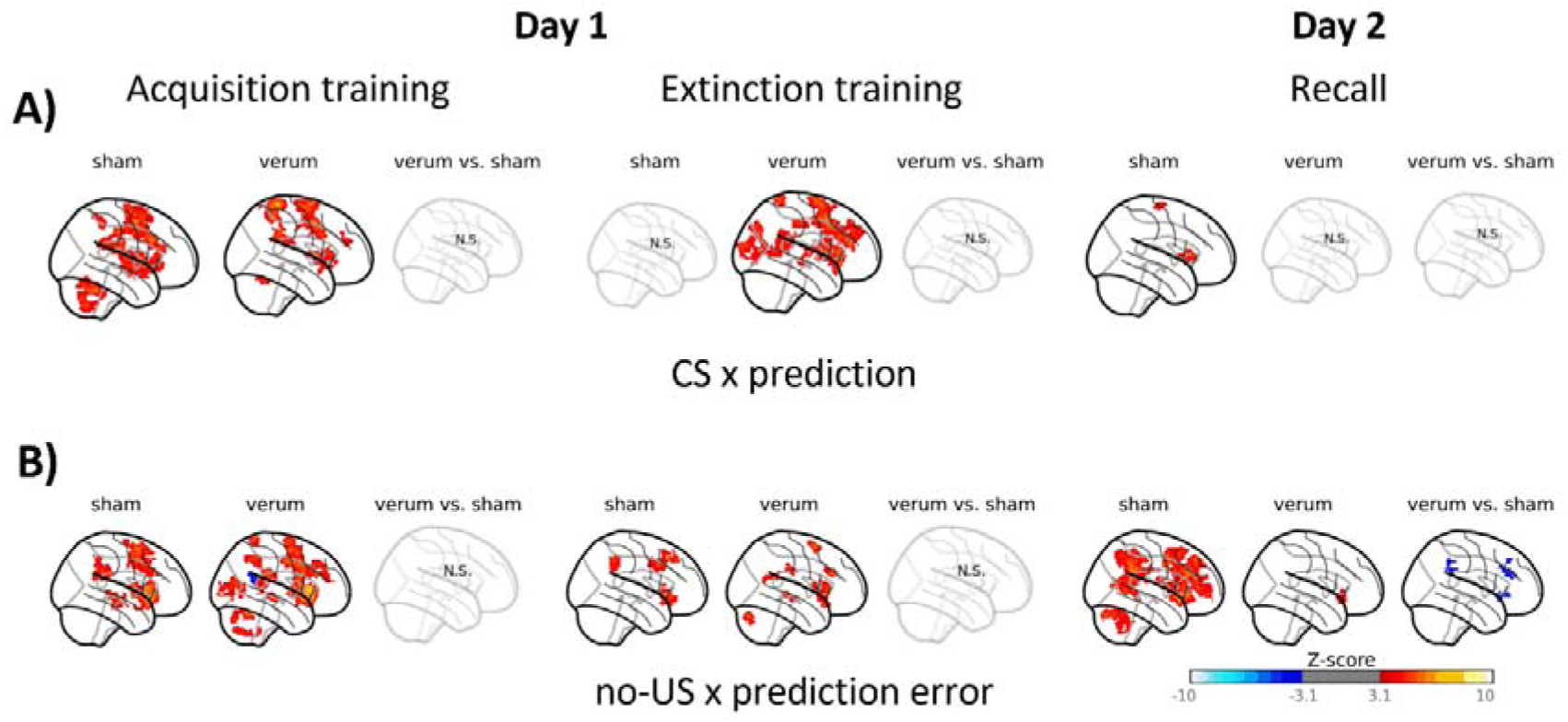
The glass brain visualization shows parametric modulation with learning model- derived data during recall. **A)** Parametric modulation of CS events with individual mean prediction values (CS x prediction), **B**) parametric modulation of omission of US events at CS termination with individual absolute mean prediction error values (no-US x prediction error). In the individual group contrasts increases are shown in red and decreases in blue. For the differential contrast ‘verum vs sham’, shades of blue denote higher activation in the sham group, while shades of red denote higher activation in the verum group. CS = conditioned stimulus; US = unconditioned stimulus; N.S. – no significant clusters. Results of fMRI analysis are provided in **Tables 5-1, 6-1** and **6-2.**

Parametric modulation effects for no-US and prediction error values revealed significant activation clusters in the ACC, insular cortex, frontal cortical areas (superior frontal gyrus [corresp. dlPFC], inferior frontal gyrus [corresp. vlPFC], frontal pole, orbitofrontal cortex, precentral gyrus [corresp. M1] and SMA), parietal cortical areas (supramarginal gyrus [corresp. S2], precuneal cortex, parietal operculum and angular gyrus) and occipital cortical areas (inferior and superior lateral occipital cortices, cuneal and intracalcarine cortices), temporal cortical areas (superior and middle temporal gyri, lingual gyrus), subcortical areas (thalamus, hippocampus, amygdala, periaqueductal gray [PAG]) and cerebellar areas (bilaterally in lobules I-IV and Crus I, left VII-VIII and Crus II, vermal VIII-IX; **Table 5-1**, **Figures 5** and **6**, shades of red). Reduced activation clusters were found in PCC, precentral gyrus [corresp. M1], postcentral gyrus [corresp. S1], as well as right cerebellar Crus I and II (**Table 5-1**, **Figures 5** and **6**, shades of blue).

### Fear extinction training

In fear extinction training, parametric modulation effects for CS and prediction values revealed no significant clusters for the sham group. However, in the verum group, significant clusters were revealed in the insular cortex, ACC, frontal cortical areas (inferior frontal gyrus [corresp. vlPFC], middle frontal gyrus [corresp. dlPFC], frontal and central operculi, orbitofrontal cortex, frontal pole, SMA), parietal cortical areas (supramarginal gyrus [corresp. S2], precuneal cortex, superior parietal lobule and parietal operculum), occipital cortex (cuneal and lateral occipital cortices, lingual gyrus, and occipital pole), temporal cortical areas (middle and superior temporal gyri) and subcortical areas (thalamus, basal ganglia) (**Table 6-1**, **Figure 6**, shades of red).

Parametric modulation effects for no-US and prediction error values revealed significant activation clusters for the sham group in frontal cortical areas (inferior frontal gyrus [corresp. vlPFC], middle and superior frontal gyri [corresp. dlPFC], frontal operculum and orbitofrontal cortex), and parietal cortex (supramarginal [corresp. S2] and angular gyri) (**Table 6-1**, **Figure 6**, shades of red). In the verum group, significant activation clusters were found in the insular cortex, cingulate cortex (ACC and PCC), frontal cortex (inferior frontal gyrus [corresp. vlPFC], frontal operculum, orbitofrontal cortex and SMA), parietal cortex (supramarginal [corresp. S2] and angular gyri), temporal cortex (superior and middle temporal gyri) (**Table 6-1**, **Figure 6**, shades of red).

No significant between-group differences for parametric modulation effects for CS and no-US events (contrasts ’verum > sham’ and ’verum < sham’) were found.

### Recall

In the recall condition, parametric modulation effects for CS and prediction values revealed significant clusters for the sham group in the insular cortex, frontal cortical areas (inferior frontal gyrus [corresp. vlPFC], superior frontal gyrus [corresp. dlPFC], frontal operculum and precentral gyrus [corresp. M1]), whereas no significant clusters were revealed in the verum group (**Table 6-2**, **Figures 6** and **6-1**, shades of red).

Parametric modulation effects for no-US and prediction error values revealed significant activation clusters for the sham group in the insular cortex, ACC, frontal cortical areas (precentral gyrus [corresp. M1], frontal operculum, orbitofrontal cortex and frontal pole), parietal cortex (supramarginal [corresp. S2] and angular gyri), lateral occipital cortex, as well as bilaterally in cerebellar lobules VI-VII and Crus I-II, and right VIII (**Table 6-2**, **Figures 6** and **6- 1**, shades of red).

In the verum group, significant activation clusters were found in the frontal cortex (inferior frontal gyrus [corresp. vlPFC], frontal operculum and orbitofrontal cortex) and temporal pole (**Table 6-2**, **Figures 6** and **6-1**, shades of red).

No significant between-group differences for parametric modulation effects for CS events (contrasts ’verum > sham’ and ’verum < sham’) were found.

Between-group comparisons for no-US events (contrasts ’verum > sham’ and ’verum < sham’) revealed significantly higher activation in the ACC, frontal cortical areas (inferior frontal gyrus [corresp. vlPFC], frontal operculum and orbitofrontal cortex) and parietal cortical (supramarginal [corresp. S2] and angular gyri), in the sham compared to the verum group (**Table 6-2**, **Figures 6** and **6-1**, shades of blue).

## Discussion

Cerebellar tACS at 6 Hz during extinction training reduced spontaneous fear memory recovery in recall. Specifically, after extinction training on day 1, testing recall of the extinguished fear association showed that participants who had received sham ctACS exhibited a return of differential SCRs when comparing the CS+ and CS-. In contrast, this return was absent in the verum group. Thus, the application of ctACS during extinction training stabilized extinction effects. fMRI data showed that activations of cerebral cortical areas associated with the spontaneous recovery of previously extinguished conditioned fear responses were significantly higher in the sham ctACS group compared to the verum group, whereas cerebral activations were enhanced during extinction training in the verum group. These findings suggest that oscillatory interactions between the cerebellum and cerebral cortical areas in the theta range contribute to extinction-related processes.

The present results in humans are further supported by findings in animal literature. Correlations between spontaneous cerebellar and medial prefrontal cortex theta activity in the 4-7 Hz frequency range and successful extinction of conditioned eyeblink responses have been found in guinea pigs (Wang et al., 2014). Similarly, significant activations of frontal cortical areas were observed during extinction learning in the verum group, but not in the sham group. Furthermore, a decrease in cerebellar theta activity following exposure to the conditioned stimulus has been linked to the spontaneous reappearance of previously extinguished eyeblink responses (Wang et al., 2019). Thus, increased cerebellar theta activity seems to support extinction learning and prevent the recall of the initial fear association after successful extinction training. Wang and colleagues have studied eyeblink conditioning, which involves the conditioning of a specific aversive response (Thompson et al., 1987). The present findings extend their observation to the extinction learning of unspecific aversive fear responses in humans. Increased cerebellar theta activity seems to stabilize extinction memory and prevent the recall of the initial fear association after successful extinction training.

Our fMRI data suggest that the activation of cerebral areas related to the return of fear is better suppressed when cerebellar tACS at 6 Hz is applied during extinction. This is in very good accordance with a recent study in mice showing that the cerebellum modulates the extinction of conditioned fear responses via its connections with the dorsomedial prefrontal cortex (dmPFC; Frontera et al., 2023). Frontera et al. (2023) found that the fastigial nucleus (FN) in mice projects via the thalamus preferentially to the dmPFC rather than the ventromedial prefrontal cortex (vmPFC), which is commonly associated with extinction learning (Dunsmoor et al., 2019; Milad et al., 2007). When FN output is inhibited, dmPFC 2-6 Hz oscillations are high and extinction learning is impaired. Thus, the cerebellum may play an important role in extinction learning by modulating the activation of cerebral areas involved in the acquisition and expression of learned fear responses.

In the present study 6 Hz ctACS applied during extinction training led to reduced spontaneous recovery, that is it reduced the return of learned fear during recall. This was accompanied by significantly less activation of cerebellar and cerebral areas which are known to contribute to fear conditioning during recall. Because of spontaneous recovery of fear memory in the sham group, we assume that the US was expected in the initial recall trial. In accordance, the strongest difference between the sham and verum group was observed at the time of the (initially unexpectedly) omitted US in parametric modulation analysis using prediction error values. Brain activation related to the unexpected US omission have been related to prediction error processing by our group and others (Ernst et al., 2019). Moreover, significantly higher activation in sham compared to verum stimulation was found in prefrontal areas known to contribute to the acquisition of learned fear including the ACC and insula region (Buchel et al., 1999; Buchel et al., 1998; Fullana et al., 2016).

Based on the findings by Frontera et al. (2023) given above one might have expected that 6 Hz cerebellar tACS would reduce extinction learning. Cerebellar granule (and Golgi) cells have been shown to oscillate in the theta frequency range (Dugue et al., 2009; Hoffmann and Berry, 2009). Theta ctACS may therefore lead to increased recruitment of cerebellar granule (and Golgi) cells. The consequences this may have on oscillations in cerebral areas are hard to predict, given that granule cells are connected to the (inhibitory) Purkinje cells, which project onto cerebellar nuclei that inform cortical areas via the thalamus (Popa et al., 2019). EEG recordings could help determine whether cerebellar tACS modulates theta oscillations in the prefrontal cortex and in which direction (increase or decrease). This will be an important area of interest for future research.

During extinction training, the verum group exhibited more widespread cerebral activations compared to the sham group. The group differences were significant in occipital cortical areas. Cortical plasticity in the visual cortex has been demonstrated to contribute to both the acquisition and extinction of conditioned fear (Petro et al., 2017; Xie et al., 2023). Cerebello-frontal connections are well known, particularly with the dlPFC, which plays a crucial role in attentional processes related to fear conditioning (Lissek et al., 2017; Middleton and Strick, 2001). An alternative explanation could be that the cerebellum contributes to enhancing selective attention towards the CS. Additionally, oscillations in the 4-8 Hz range within visual cortical areas have been described in relation to the processing of visual stimuli (Gao et al., 2021; Levy et al., 2017; Tang et al., 2023). Thus, activations in the visual cortex may reflect mechanisms by which visual information is encoded and processed during fear extinction learning, potentially influenced by interactions with the cerebellum. Note that extinction effects appeared to be stronger in the verum compared to the sham group in late extinction, although this was not reflected by the results of the statistical analysis. One cannot exclude, however, that direct occipital stimulation effects have at least partially contributed.

The present findings may also have clinical implications. Extinction learning forms the basis of exposure therapy, a widely used technique for addressing fear memories associated with conditions such as posttraumatic stress disorder (PTSD) (Milad et al., 2008). Our findings suggest that 6 Hz ctACS may be a way to enhance the efficacy of exposure therapy. Most studies aiming to enhance extinction have administered transcranial direct current stimulation (tDCS) directed to cerebral cortical areas, particularly parts of prefrontal cortex. Findings, however, are heterogeneous, which is a common observation in many tDCS studies (Adams et al., 2020; Markovic et al., 2021, for review). tDCS is direction-dependent which makes it difficult to predict the outcome, particularly in highly convoluted structures like the cerebellum (Oldrati and Schutter, 2018). tACS uses an oscillating current that is able to entrain neural oscillations at specific frequencies and is direction-independent. Thus, cerebellar tACS may reduce variability in outcomes and offer better reproducibility compared to cerebellar tDCS. Further studies, however, are needed to validate the present findings, including testing different timings of application and stimulation frequencies. Furthermore, findings need to be replicated using a three-day design, with acquisition and extinction learning being performed on separate days. Of note in a previous study of our group we did not see ctACS effects on fear extinction learning (Schellen et al., 2023). The paradigm used, however, was different, including changes of context, and there were flaws in the design (reinforcement of the CS- which was not intended).

In sum, the present data demonstrates that ctACS enhances cerebral cortical activation during extinction which is followed by enhanced recall of the extinction memory and subsequently reduced activation of cerebral cortical areas associated to spontaneous recovery of fear memory. The present study offers causal evidence that the cerebellum is involved in the fear extinction network in humans. The downregulation of cerebral areas involved in acquisition and expression of fear associations highlights the significant contribution of the cerebellum to extinction processes.

## Conflicts of interest

The authors report no conflict of interest.

## Funding sources

This work was supported by a grant from the German Research Foundation (DFG; project number 316803389 – SFB 1280) to D.T. (subproject A05), C.J.M. (subproject A09), M.N. (subproject A06) and S.C. (subproject F01). At the time of this study A.T. held a position that was funded in part by the University Medicine Essen Clinician Scientist Academy (UMEA) and the German Research Foundation (DFG; grant no. FU356/12-2).

## Acknowledgments

The authors would like to express their gratitude to G. Wippich for creating the experimental paradigm figure.

## CRediT authorship contribution statement

**Andreas Thieme:** Investigation, Formal Analysis, Writing – Original Draft, Writing – Review & Editing.

**Zsofia Spisak:** Formal Analysis, Software, Writing – Review & Editing.

**Philippe Zeidan:** Investigation.

**Michael Klein:** Investigation, Formal Analysis.

**Enzo Nio:** Formal Analysis, Software.

**Thomas M. Ernst:** Methodology, Software, Validation, Resources, Data Curation, Writing – Review & Editing.

**Nicolas Diekmann**: Methodology, Formmal analysis, Software, Writing – review and editing.

**Sophia Göricke:** Resources.

**Sen Cheng:** Conceptualization, Methodology, Writing – review and editing.

**Christian J. Merz:** Conceptualization, Methodology, Writing – Review & Editing. **Fatemeh Yavari:** Conceptualization, Methodology, Software, Writing – Review & Editing. **Michael A. Nitsche:** Conceptualization, Methodology, Writing – Review & Editing.

**Giorgi Batsikadze:** Investigation, Conceptualization, Methodology, Software, Formal Analysis, Visualization, Project Administration, Supervision, Writing – Original Draft, Writing – Review & Editing. **Dagmar Timmann:** Conceptualization, Methodology, Supervision, Project Administration, Funding Acquisition, Writing – Original Draft, Writing – Review & Editing.

## Notes

### Competing Interest Statement

The authors have declared no competing interest.

