## Extended Data for "Cerebellar transcranial alternating current stimulation in the theta band facilitates extinction of learned fear responses"

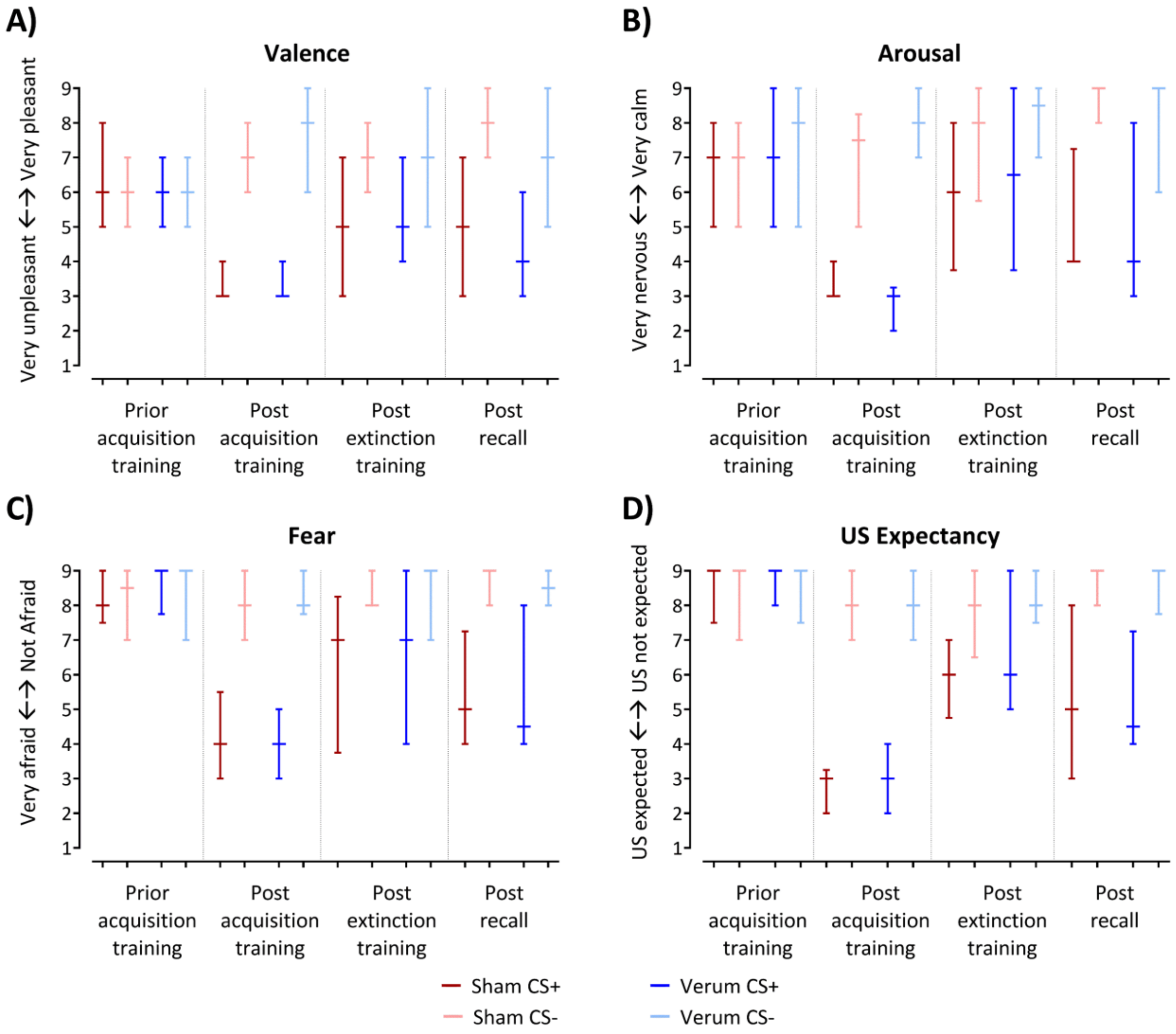


**Figure 3-1**. Median ratings regarding **A)** valence, **B)** arousal, **C)** fear and **D)** US expectancy obtained using a Likert-scale ranging from 1 (*"very pleasant" / "very calm" / "not afraid", "US not expected"*, respectively) to 9 (*"very unpleasant" / "very nervous" / "very afraid", "US expected"*, respectively). The median values are represented by horizontal lines, and the whiskers extend from the first to the third quartile. The sham group is represented by red colors, the verum group by blue colors. Dark colors: CS+, light colors: CS-.

Prior acquisition training, valence, arousal and fear ratings of the CS+ and CS- were not significantly different. Post acquisition training, the CS+ was rated as significantly less pleasant compared to the CS-, arousal and fear towards the CS+ were rated significantly higher compared to the CS- and these differences remained until the end of the recall phase (least square means test, all $p < 0.001$). ATS revealed a significant main effect of Time (all $p$ values $\leq 0.005$), Stimulus (all $p$ values $< 0.001$) and a significant Stimulus x Time interaction (all $p$ values $< 0.001$). *Post hoc* tests showed significant difference between stimuli post-acquisition training, extinction training and recall (least square means tests, all $p$ values $< 0.001$), but not prior acquisition training (least square means test, all $p \geq 0.313$).

Prior acquisition training, the reported US expectation after CS+ and CS- was not significantly different. Post acquisition training, participants reported significantly higher US expectation after the CS+ compared to the CS- and this difference remained until the end of the recall phase. ATS revealed a significant main effect of Time ($F_{2.73} = 27.73$, $p < 0.001$), Stimulus ($F_1 = 109.09$, $p < 0.001$), and a significant Stimulus x Time ($F_{2.79} = 42.16$, $p < 0.001$) interaction. *Post hoc* tests showed a significant difference between stimuli post-acquisition, extinction and recall phases (least square means tests, all $p$ values < 0.001), but not prior acquisition training ($p = 0.485$).

**Table 3-1.** Results of the non-parametric ANOVA-type statistics (ATS) for repeated measures for skin conductance responses (SCRs), differential skin conductance responses (SCRdiff), valence, arousal, fear, US expectancy and stimulation side-effect ratings comparing cerebellar verum and sham groups.

| **Factor** | **Numerator Df** | ***F*** | ***P*** |
|---|---|---|---|
| | **Skin conductance responses** | | |
| | *Habituation* | | |
| Stimulus | 1 | 2.59 | 0.108 |
| Group | 1 | 0.02 | 0.882 |
| Stimulus × Group | 1 | <0.01 | 0.958 |
| | *Fear acquisition training* | | |
| Stimulus | 1 | 56.75 | **<.001*** |
| Block | 1 | 57.62 | **<.001*** |
| Group | 1 | 0.10 | 0.748 |
| Stimulus × Block | 1 | 0.04 | **0.030*** |
| Block × Group | 1 | 4.69 | 0.117 |
| Stimulus × Group | 1 | 2.45 | 0.841 |
| Stimulus × Block × Group | 1 | 0.11 | 0.738 |
| | *Extinction training* | | |
| Stimulus | 1 | 12.93 | **<.001*** |
| Block | 1 | 32.25 | **<.001*** |
| Group | 1 | 1.20 | 0.274 |
| Stimulus × Block | 1 | 0.01 | 0.278 |
| Block × Group | 1 | 1.18 | 0.931 |
| Stimulus × Group | 1 | 0.01 | 0.913 |
| Stimulus × Block × Group | 1 | 1.34 | 0.247 |
| | *Late acquisition training vs early extinction training* | | |
| Stimulus | 1 | 43.24 | **<.001*** |
| Block | 1 | 0.00 | 0.983 |
| Group | 1 | 0.65 | 0.422 |
| Stimulus × Block | 1 | 8.11 | **0.004*** |
| Block × Group | 1 | 0.00 | 0.998 |
| Stimulus × Group | 1 | 0.43 | 0.512 |
| Stimulus × Block × Group | 1 | 0.02 | 0.895 |
| | *Recall* | | |
| Stimulus | 1 | 10.20 | **0.001*** |
| Block | 1 | 74.03 | **<.001*** |
| Group | 1 | 0.35 | 0.551 |
| Stimulus × Block | 1 | 0.93 | 0.269 |
| Block × Group | 1 | 1.22 | **<.001*** |
| Stimulus × Group | 1 | 12.59 | 0.336 |
| Stimulus × Block × Group | 1 | 1.80 | 0.179 |
| | **Differential skin conductance responses (SCRdiff)** | | |
| | *Habituation* | | |
| Group | 1 | 0.06 | 0.806 |
| | *Fear acquisition training* | | |
| Block | 1 | 3.37 | 0.066 |
| Group | 1 | 0.05 | 0.816 |
| Block × Group | 1 | 0.75 | 0.387 |
| | *Extinction training* | | |
| Block | 1 | 3.65 | 0.056 |

| | | | |
|---|---|---|---|
| Group | 1 | 0.34 | 0.559 |
| Block × Group | 1 | 0.58 | 0.445 |
| *Late acquisition training vs early extinction training* | | | |
| Block | 1 | 8.97 | **0.003*** |
| Group | 1 | 0.30 | 0.582 |
| Block × Group | 1 | 0.20 | 0.654 |
| *Recall* | | | |
| Block | 1 | 11.30 | **<.001*** |
| Group | 1 | 2.31 | 0.129 |
| Block × Group | 1 | 4.82 | **0.028*** |
| **Valence** | | | |
| Group | 1 | 0.13 | 0.718 |
| Stimulus | 1 | 79.07 | **<.001*** |
| Time | 2.44 | 4.80 | **0.005*** |
| Group × Stimulus | 1 | 0.11 | 0.743 |
| Group × Time | 2.44 | 0.30 | 0.786 |
| Stimulus × Time | 2.39 | 43.57 | **<.001*** |
| Group × Stimulus × Time | 2.39 | 0.65 | 0.551 |
| **Arousal** | | | |
| Group | 1 | 0.14 | 0.713 |
| Stimulus | 1 | 124.18 | **<.001*** |
| Time | 2.58 | 11.18 | **<.001*** |
| Group × Stimulus | 1 | 0.28 | 0.600 |
| Group × Time | 2.58 | 0.91 | 0.424 |
| Stimulus × Time | 2.27 | 28.72 | **<.001*** |
| Group × Stimulus × Time | 2.27 | 1.10 | 0.339 |
| **Fear** | | | |
| Group | 1 | 0.18 | 0.667 |
| Stimulus | 1 | 79.29 | **<.001*** |
| Time | 2.76 | 17.39 | **<.001*** |
| Group × Stimulus | 1 | 0.36 | 0.548 |
| Group × Time | 2.76 | 0.56 | 0.628 |
| Stimulus × Time | 2.66 | 26.89 | **<.001*** |
| Group × Stimulus × Time | 2.66 | 1.29 | 0.278 |
| **US expectancy** | | | |
| Group | 1 | 0.08 | 0.776 |
| Stimulus | 1 | 109.09 | **<.001*** |
| Time | 2.73 | 27.73 | **<.001*** |
| Group × Stimulus | 1 | 1.36 | 0.244 |
| Group × Time | 2.73 | 1.15 | 0.324 |
| Stimulus × Time | 2.79 | 42.16 | **<.001*** |
| Group × Stimulus × Time | 2.79 | 0.53 | 0.647 |
| **Stimulation side-effects** | | | |
| **Headache** | | | |
| Group | 1 | 0.04 | 0.849 |
| Time | 1 | 6.28 | **0.012*** |
| Group × Time | 1 | 2.09 | 0.148 |
| **Neck pain** | | | |
| Group | 1 | 0.23 | 0.630 |
| Time | 1 | 0.13 | 0.723 |
| Group × Time | 1 | 1.37 | 0.241 |
| **Back pain** | | | |
| Group | 1 | 0.15 | 0.698 |
| Time | 1 | 1.23 | 0.268 |

| | | | |
|---|---|---|---|
| Group × Time | 1 | 1.05 | 0.305 |
| **Blurred vision** | | | |
| Group | 1 | 0.01 | 0.929 |
| Time | 1 | 2.60 | 0.107 |
| Group × Time | 1 | 0.67 | 0.411 |
| **Scalp irritation** | | | |
| Group | 1 | 0.70 | 0.403 |
| Time | 1 | 3.60 | 0.058 |
| Group × Time | 1 | 0.64 | 0.423 |
| **Scalp tingling** | | | |
| Group | 1 | 4.22 | **0.040*** |
| Time | 1 | 1.14 | 0.286 |
| Group × Time | 1 | 0.19 | 0.665 |
| **Scalp itching** | | | |
| Group | 1 | 0.22 | 0.637 |
| Time | 1 | 0.25 | 0.615 |
| Group × Time | 1 | 0.18 | 0.669 |
| **Increased heartbeat** | | | |
| Group | 1 | 0.20 | 0.659 |
| Time | 1 | 5.45 | **0.020*** |
| Group × Time | 1 | 0.06 | 0.799 |
| **Burning sensation** | | | |
| Group | 1 | 0.07 | 0.795 |
| Time | 1 | 0.08 | 0.778 |
| Group × Time | 1 | 0.41 | 0.523 |
| **Hot flashes** | | | |
| Group | 1 | 1.15 | 0.284 |
| Time | 1 | 0.68 | 0.409 |
| Group × Time | 1 | 0.01 | 0.934 |
| **Vertigo** | | | |
| Group | 1 | 0.46 | 0.497 |
| Time | 1 | 0.58 | 0.448 |
| Group × Time | 1 | 0.27 | 0.606 |
| **Sudden mood change** | | | |
| Group | 1 | 0.03 | 0.852 |
| Time | 1 | 7.73 | **0.005*** |
| Group × Time | 1 | 0.01 | 0.936 |
| **Fatigue** | | | |
| Group | 1 | 0.08 | 0.773 |
| Time | 1 | 6.22 | **0.013*** |
| Group × Time | 1 | 0.38 | 0.539 |
| **Phosphenes** | | | |
| Group | 1 | 0.18 | 0.672 |
| Time | 1 | 2.02 | 0.155 |
| Group × Time | 1 | 0.24 | 0.624 |

* Significant results at $p < 0.05$.

**Table 3-2.** Results of side-effects questionnaires - Self-reported median ratings and interquartile range (in brackets) prior and post ctACS administration.

| Questionnaire item | Prior stimulation | | Post stimulation | |
|---|---|---|---|---|
| | *Sham group* | *Verum group* | *Sham group* | *Verum group* |
| *Headache* | **1 (1 - 2)*** | **1 (1 - 2)*** | **1 (1 - 3.75)*** | **1.5 (1 - 4)*** |
| *Neck pain* | 1 (1 - 2) | 1 (1 - 2.75) | 1 (1 - 2) | 1 (1 - 2.75) |
| *Back pain* | 1 (1 - 2) | 1 (1 - 2) | 1 (1 - 1.75) | 1 (1 - 1.75) |
| *Blurred vision* | 1.5 (1 - 2) | 1 (1 - 3.75) | 2 (1 - 3.75) | 1.5 (1 - 3) |
| *Scalp irritation* | 1 (1 - 1) | 1 (1 - 1) | 1 (1 - 2) | 1 (1 - 1) |
| *Scalp tingling* | **1 (1 - 1)†** | **1 (1 - 1)†** | **1 (1 - 1)†** | **1 (1 - 1)†** |
| *Scalp itching* | 1 (1 - 1) | 1 (1 - 1) | 1 (1 - 1) | 1 (1 - 1) |
| *Increased heartbeat* | **2.5 (1.25 - 5)*** | **3 (2 - 5.75)*** | **2 (1.25 - 3.75)*** | **2 (2 - 4)*** |
| *Burning sensation* | 1 (1 - 1.75) | 1 (1 - 1.75) | 1 (1 - 1) | 1 (1 - 2) |
| *Hot flashes* | 1 (1 - 2) | 1 (1 - 3) | 1 (1 - 1.75) | 1 (1 - 3) |
| *Vertigo* | 1 (1 - 1) | 1 (1 - 1) | 1 (1 - 1.75) | 1 (1 - 1) |
| *Sudden mood change* | **1.5 (1 - 2)*** | **1.5 (1 - 4)*** | **1 (1 - 2)*** | **1 (1 - 2)*** |
| *Fatigue* | **2 (1 - 3)*** | **2 (1 - 3.75)*** | **2.5 (2 - 3)*** | **4 (1 - 6)*** |
| *Phosphenes* | 1 (1 - 1) | 1 (1 - 1) | 1 (1 - 1) | 1 (1 - 1) |

Statistical significances (least squares means tests, $p < 0.05$) are indicated in bold.
* Significant differences between prior- and post-ctACS administration.
† Significant differences between sham and verum group.

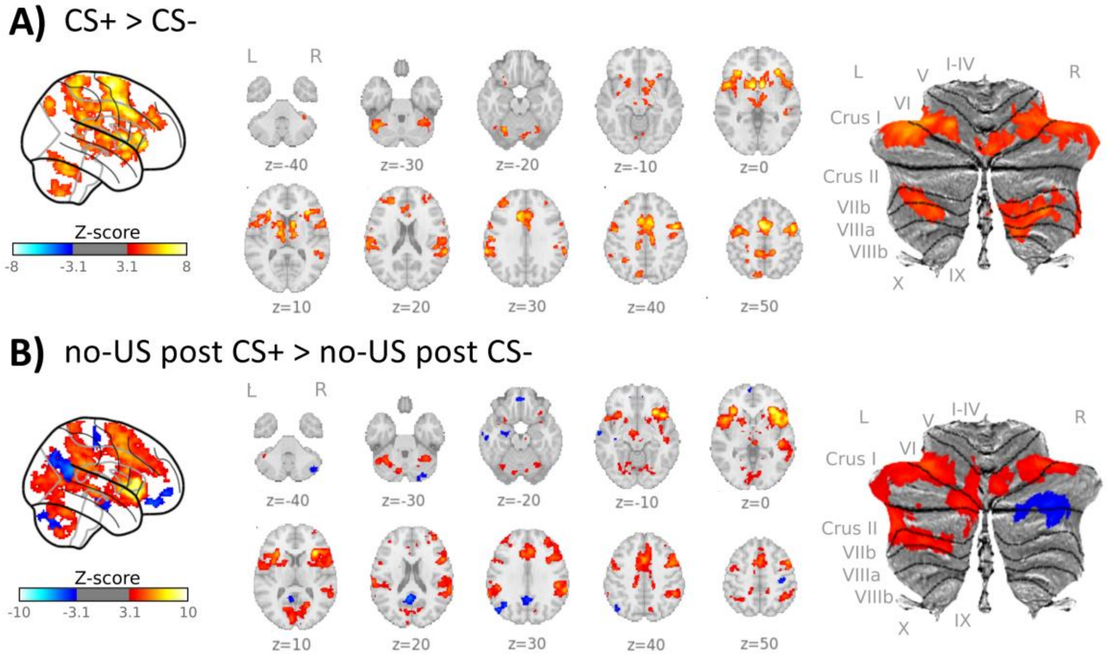


**Figure 4-1.** Cerebral and cerebellar activations during fear acquisition training related to the **A)** presentation of the CSs (contrast 'CS+ > CS-') and **B)** omission of the aversive US (contrast 'no-US post CS+ > no-US post CS-'). Data from both the verum and sham groups are analyzed together as a combined sample.

CS = conditioned stimulus; L = left; R = right; SUIT = spatially unbiased atlas template of the cerebellum; results of fMRI analysis are provided in **Table 4-1.**

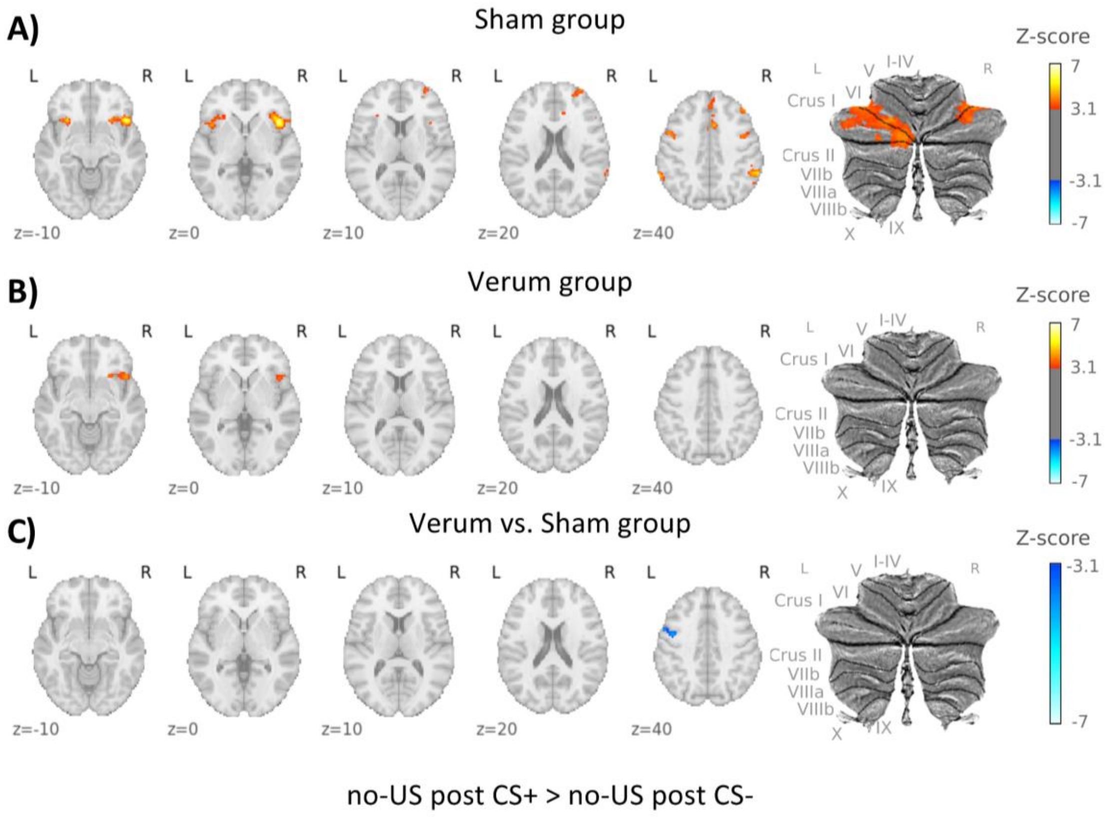


**Figure 4-2.** Cerebral and cerebellar activations during recall related to the omission of the aversive US (contrast 'no-US post CS+ > no-US post CS-') in **A)** the sham group, **B)** the verum group and **C)** for the comparison between verum and sham groups (contrast 'verum vs. sham'). In the individual groups, contrast increases are shown in red and decreases in blue.

In the differential contrast 'verum vs sham', shades of blue denote higher activation in the sham group, while shades of red denote higher activation in the verum group. CS = conditioned stimulus; L = left; R = right; SUIT = spatially unbiased atlas template of the cerebellum; Results of the fMRI analysis are provided in **Table 4-3.**

**Table 4-1.** Fear acquisition training. Activation clusters are reported. Listed below are all local maxima within significant clusters ($p < 0.05$, corrected for multiple comparisons using a cluster-forming threshold of $Z = 3.1$). The anatomical locations of these maxima were determined by referencing the Harvard-Oxford Cortical and Subcortical, and Cerebellar Atlases in MNI152 space following normalization with FLIRT. Only structures with a probability of 25% or higher are reported. For regions where the Harvard-Oxford atlas was insufficient for identification, manual identification was conducted. Brain areas manually identified using the Multi-Contrast Anatomical Subcortical Structures (MASSP) atlas (Bazin et al., 2020) are indicated in square brackets. Percentages represent the likelihood of anatomical localization.

| Index | Cluster size / voxel | *p* value | Z | Coordinates / mm | | | Name (Harvard-Oxford Atlas) |
|---|---|---|---|---|---|---|---|
| | | | | x | y | z | |
| *Both groups, acquisition training, CS+ > CS-* | | | | | | | |
| 1 | 7157 | < 0.001 | 8.08 | -4 | 6 | 48 | 51% Juxtapositional Lobule Cortex (formerly Supplementary Motor Cortex) |
| | | | 7.74 | 2 | -6 | 66 | 63% Juxtapositional Lobule Cortex (formerly Supplementary Motor Cortex) |
| | | | 7.63 | 6 | -6 | 66 | 53% Juxtapositional Lobule Cortex (formerly Supplementary Motor Cortex) |
| | | | 7.17 | -10 | 14 | 34 | 34% Cingulate Gyrus, anterior division |
| | | | 6.93 | -6 | -6 | 70 | 36% Juxtapositional Lobule Cortex (formerly Supplementary Motor Cortex) |
| | | | 6.82 | 10 | 6 | 44 | 31% Juxtapositional Lobule Cortex (formerly Supplementary Motor Cortex), 26% Cingulate Gyrus, anterior division |
| 2 | 4265 | < 0.001 | 7.72 | 8 | 10 | 0 | 81% Right Caudate, [Right Red Nucleus] |
| | | | 7.41 | 10 | 8 | 8 | 85% Right Caudate |
| | | | 6.99 | -8 | 8 | 2 | 77% Left Caudate |
| | | | 6.76 | -30 | 26 | 6 | 33% Insular Cortex |
| | | | 6.31 | -14 | 10 | -4 | 50% Left Cerebral White Matter, 31% Left Putamen, [Left Red Nucleus] |
| | | | 6.03 | -48 | 10 | 0 | 29% Frontal Operculum Cortex |
| 3 | 2227 | < 0.001 | 6.70 | 46 | 2 | 54 | 36% Middle Frontal Gyrus, 30% Precentral Gyrus |
| | | | 6.65 | 34 | 26 | 8 | 32% Frontal Operculum Cortex |
| | | | 6.46 | 50 | 2 | 46 | 70% Precentral Gyrus |
| | | | 6.40 | 50 | -2 | 48 | 66% Precentral Gyrus |
| | | | 6.20 | 42 | 28 | 6 | 27% Inferior Frontal Gyrus, pars triangularis |
| 4 | 997 | < 0.001 | 5.69 | 62 | -40 | 22 | 59% Supramarginal Gyrus, posterior division |
| | | | 4.96 | 58 | -44 | 8 | 50% Middle Temporal Gyrus, temporooccipital part |
| | | | 4.88 | 66 | -36 | 30 | 44% Supramarginal Gyrus, posterior division |
| | | | 4.88 | 48 | -30 | -4 | 51% Middle Temporal Gyrus, posterior division |
| | | | 4.68 | 60 | -26 | 24 | 35% Supramarginal Gyrus, anterior division, 26% Parietal Operculum Cortex |
| 5 | 936 | < 0.001 | 6.10 | -40 | -4 | 54 | 44% Precentral Gyrus, 25% Middle Frontal Gyrus |
| | | | 5.46 | -46 | -10 | 52 | 72% Precentral Gyrus |
| | | | 5.24 | -30 | -8 | 48 | 45% Precentral Gyrus |
| | | | 5.23 | -48 | -4 | 42 | 52% Precentral Gyrus |
| | | | 4.30 | -60 | 2 | 26 | 79% Precentral Gyrus |
| 6 | 892 | < 0.001 | 5.87 | -64 | -38 | 28 | 54% Supramarginal Gyrus, anterior division |
| | | | 5.16 | -64 | -28 | 22 | 54% Supramarginal Gyrus, anterior division |
| | | | 5.06 | -56 | -38 | 36 | 42% Supramarginal Gyrus, anterior division |
| | | | 4.94 | -56 | -42 | 36 | 39% Supramarginal Gyrus, posterior division |
| | | | 4.90 | -52 | -34 | 26 | 41% Parietal Operculum Cortex |
| | | | 4.89 | -56 | -20 | 22 | 29% Postcentral Gyrus |
| 7 | 673 | < 0.001 | 5.66 | -26 | -64 | -20 | 94% Left VI |
| | | | 5.62 | -34 | -56 | -26 | 92% Left VI |
| | | | 5.47 | -34 | -52 | -34 | 58% Left VI, 42% Left Crus I |
| | | | 5.46 | -40 | -58 | -32 | 100% Left Crus I |
| | | | 4.66 | -44 | -66 | -24 | 70% Left Crus I |
| | | | 3.89 | -50 | -54 | -34 | 85% Left Crus I |
| 8 | 634 | < 0.001 | 5.52 | -42 | 46 | 26 | 52% Frontal Pole |
| | | | 5.10 | -32 | 40 | 34 | 53% Frontal Pole, 26% Middle Frontal Gyrus |
| | | | 5.01 | -22 | 44 | 20 | 35% Frontal Pole |

| | | | | | | | |
|---|---|---|---|---|---|---|---|
| | | | 4.66 | -32 | 42 | 24 | 59% Frontal Pole |
| | | | 3.94 | -34 | 52 | 26 | 70% Frontal Pole |
| | | | 3.89 | -38 | 30 | 34 | 58% Middle Frontal Gyrus |
| 9 | 580 | < 0.001 | 5.01 | 42 | -52 | -34 | 100% Right Crus I |
| | | | 4.79 | 36 | -56 | -26 | 89% Right VI |
| | | | 4.66 | 36 | -42 | -36 | 86% Right VI |
| | | | 4.65 | 38 | -50 | -28 | 86% Right VI |
| | | | 4.63 | 34 | -52 | -32 | 72% Right VI |
| | | | 4.47 | 28 | -56 | -24 | 100% Right VI |
| 10 | 418 | < 0.001 | 5.38 | 34 | -64 | -58 | 49% Right VIIIa, 46% Right VIIb |
| | | | 4.50 | 10 | -64 | -54 | 35% Right VIIIb, 31% Right VIIIa, 29% Right IX |
| | | | 4.33 | 22 | -70 | -50 | 69% Right VIIb |
| | | | 4.20 | 16 | -64 | -56 | 58% Right VIIIa, 37% Right VIIIb |
| | | | 4.09 | 22 | -68 | -58 | 64% Right VIIIa |
| | | | 3.66 | 34 | -54 | -56 | 76% Right VIIIa |
| 11 | 293 | < 0.001 | 5.30 | -8 | -76 | 46 | 52% Precuneal Cortex |
| 12 | 252 | < 0.001 | 4.90 | -36 | -62 | -56 | 74% Left VIIb |
| | | | 4.79 | -26 | -66 | -56 | 66% Left VIIIa, 34% Left VIIb |
| | | | 4.22 | -30 | -58 | -50 | 50% Left VIIIa, 32% Left VIIb |
| | | | 3.25 | -40 | -54 | -52 | 46% Left VIIb, 29% Left Crus II, 25% Left VIIIa |
| 13 | 207 | 0.001 | 5.03 | -28 | -54 | 46 | 38% Superior Parietal Lobule |
| 14 | 138 | 0.012 | 4.04 | 40 | 50 | 18 | 85% Frontal Pole |
| | | | 3.97 | 32 | 52 | 24 | 87% Frontal Pole |
| | | | 3.75 | 34 | 42 | 30 | 62% Frontal Pole |
| | | | 3.68 | 32 | 44 | 22 | 60% Frontal Pole |
| | | | 3.64 | 28 | 48 | 20 | 69% Frontal Pole |
| 15 | 130 | 0.017 | 4.53 | 6 | -76 | -18 | 40% Vermis VI, 35% Right VI |
| | | | 3.85 | 8 | -70 | -12 | 46% Right VI |
| | | | 3.81 | 16 | -68 | -20 | 95% Right VI |
| | | | 3.76 | -4 | -74 | -12 | 29% Lingual Gyrus |
| *Both groups, acquisition training, no-US post CS+ > no-US post CS-* | | | | | | | |
| 1 | 8869 | < 0.001 | 9.53 | 44 | 20 | -6 | 40% Frontal Orbital Cortex |
| | | | 8.90 | 34 | 24 | 8 | 46% Frontal Operculum Cortex |
| | | | 8.68 | 34 | 24 | 4 | 32% Insular Cortex, 28% Frontal Operculum Cortex |
| | | | 8.67 | 36 | 26 | 0 | 34% Frontal Orbital Cortex, 26% Frontal Operculum Cortex |
| | | | 8.61 | 32 | 26 | -8 | 58% Frontal Orbital Cortex |
| | | | 7.67 | 58 | 12 | 4 | 41% Inferior Frontal Gyrus, pars opercularis |
| 2 | 4351 | < 0.001 | 6.41 | -2 | 14 | 38 | 56% Cingulate Gyrus, anterior division, 30% Paracingulate Gyrus, |
| | | | 6.38 | 2 | 8 | 56 | 30% Juxtapositional Lobule Cortex (formerly Supplementary Motor Cortex) |
| | | | 6.36 | 2 | 12 | 38 | 72% Cingulate Gyrus, anterior division, |
| | | | 5.85 | 6 | 26 | 32 | 46% Paracingulate Gyrus, 36% Cingulate Gyrus, anterior division |
| | | | 5.76 | -8 | 4 | 44 | 44% Juxtapositional Lobule Cortex (formerly Supplementary Motor Cortex) |
| | | | 5.76 | 4 | -6 | 68 | 64% Juxtapositional Lobule Cortex (formerly Supplementary Motor Cortex) |
| 3 | 3355 | < 0.001 | 7.85 | -46 | 14 | -4 | 33% Frontal Operculum Cortex |
| | | | 7.28 | -34 | 26 | 0 | 38% Frontal Orbital Cortex |
| | | | 6.32 | -38 | 16 | -4 | 76% Insular Cortex |
| | | | 5.82 | -58 | 6 | 4 | 51% Precentral Gyrus |
| | | | 5.74 | -36 | 10 | 10 | 44% Frontal Operculum Cortex, 28% Central Opercular Cortex |
| | | | 5.72 | -54 | 2 | 6 | 42% Central Opercular Cortex, 26% Precentral Gyrus |
| 4 | 2980 | < 0.001 | 5.98 | 22 | -66 | 8 | 46% Intracalcarine Cortex |
| | | | 5.71 | -32 | -58 | -28 | 89% Left VI |
| | | | 5.14 | -34 | -64 | -54 | 71% Left VIIb |
| | | | 5.13 | -6 | -86 | 10 | 28% Intracalcarine Cortex |
| 5 | 796 | < 0.001 | 6.07 | -48 | -36 | 28 | 37% Parietal Operculum Cortex |
| | | | 5.50 | -64 | -24 | 20 | 36% Supramarginal Gyrus, anterior division, 32% Postcentral Gyrus |
| | | | 5.24 | -58 | -36 | 26 | 37% Parietal Operculum Cortex |
| | | | 4.92 | -66 | -38 | 26 | 40% Supramarginal Gyrus, anterior division |
| | | | 4.67 | -64 | -44 | 30 | 48% Supramarginal Gyrus, posterior division |
| | | | 4.64 | -62 | -50 | 30 | 51% Supramarginal Gyrus, posterior division, 29% Angular Gyrus |
| 6 | 409 | < 0.001 | 5.00 | -8 | -8 | -10 | [Left Subthalamic Nucleus, Left Ventral Tegmental Area] |
| | | | 4.85 | 6 | -24 | -4 | [Right Red Nucleus, Third Ventricle] |

| | | | | | | | |
|---|---|---|---|---|---|---|---|
| | | | 4.78 | 20 | -14 | -6 | 70% Right Cerebral White Matter, 19% Right Pallidum |
| | | | 4.61 | 12 | -8 | 6 | 98% Right Thalamus |
| | | | 4.50 | 10 | -20 | -18 | [Right Substantia Nigra] |
| | | | 3.89 | 8 | -16 | -16 | [Right Substantia Nigra, right Ventral Tegmental Area] |
| 7 | 360 | < 0.001 | 5.55 | 36 | -56 | -26 | 89% Right VI |
| | | | 4.43 | 28 | -60 | -24 | 100% Right VI |
| | | | 4.06 | 34 | -64 | -22 | 67% Right VI |
| 8 | 212 | < 0.001 | 5.12 | 18 | -64 | -6 | 49% Lingual Gyrus |
| | | | 4.66 | 10 | -74 | -18 | 90% Right VI |
| | | | 4.12 | 10 | -72 | -12 | 37% Lingual Gyrus, 27% Right VI |
| | | | 3.46 | 28 | -78 | -10 | 59% Occipital Fusiform Gyrus |
| | | | 3.45 | 20 | -72 | -16 | 43% Right VI, 25% Occipital Fusiform Gyrus |
| | | | 3.38 | 26 | -70 | -8 | 59% Occipital Fusiform Gyrus |
| 9 | 201 | 0.001 | 4.59 | -42 | -54 | 52 | 28% Angular Gyrus |
| | | | 4.48 | -50 | -50 | 56 | 32% Supramarginal Gyrus, posterior division |
| | | | 4.04 | -28 | -54 | 48 | 38% Superior Parietal Lobule |
| | | | 3.76 | -46 | -48 | 46 | 33% Supramarginal Gyrus, posterior division |
| | | | 3.67 | -34 | -56 | 52 | 34% Superior Parietal Lobule |
| 10 | 195 | 0.001 | 4.30 | -34 | 46 | 32 | 83% Frontal Pole |
| | | | 4.26 | -40 | 44 | 26 | 82% Frontal Pole |
| | | | 4.10 | -34 | 54 | 26 | 49% Frontal Pole |
| | | | 3.71 | -28 | 36 | 30 | 31% Middle Frontal Gyrus |
| 11 | 186 | 0.001 | 5.40 | 18 | -52 | 70 | 50% Superior Parietal Lobule |
| | | | 4.77 | 18 | -46 | 70 | 37% Superior Parietal Lobule |
| 12 | 143 | 0.005 | 3.96 | 18 | -74 | 36 | 26% Precuneal Cortex |
| | | | 3.94 | 12 | -70 | 42 | 36% Precuneal Cortex |
| | | | 3.34 | 16 | -68 | 32 | 40% Precuneal Cortex |
| | | | 3.31 | 12 | -74 | 30 | 41% Cuneal Cortex |
| *Both groups, acquisition training, no-US post CS+ < no-US post CS-* | | | | | | | |
| 1 | 653 | < 0.001 | 6.11 | -4 | -52 | 22 | 62% Cingulate Gyrus, posterior division |
| | | | 3.64 | -8 | -52 | 6 | 39% Precuneal Cortex, 30% Cingulate Gyrus, posterior division |
| | | | 3.41 | -4 | -40 | 34 | 79% Cingulate Gyrus, posterior division |
| 2 | 428 | < 0.001 | 4.92 | -8 | 58 | -6 | 47% Frontal Pole |
| | | | 4.85 | 2 | 40 | -16 | 85% Frontal Medial Cortex Gyrus |
| | | | 4.84 | 6 | 48 | -16 | 56% Frontal Medial Cortex |
| | | | 4.69 | -2 | 46 | -18 | 93% Frontal Medial Cortex |
| | | | 4.26 | -4 | 36 | -18 | 73% Frontal Medial Cortex |
| 3 | 304 | < 0.001 | 4.95 | -48 | -72 | 32 | 85% Lateral Occipital Cortex, superior division |
| | | | 4.52 | -46 | -72 | 40 | 87% Lateral Occipital Cortex, superior division |
| | | | 3.93 | -44 | -62 | 34 | 43% Lateral Occipital Cortex, superior division |
| 4 | 292 | < 0.001 | 4.53 | 40 | -74 | -36 | 81% Right Crus I |
| | | | 4.43 | 44 | -74 | -46 | 60% Right Crus II |
| | | | 4.03 | 30 | -80 | -30 | 95% Right Crus I |
| | | | 3.81 | 24 | -88 | -32 | 46% Right Crus II, 33% Right Crus I |
| | | | 3.49 | 32 | -88 | -36 | 10% Right Crus II |
| 5 | 183 | 0.001 | 4.53 | 36 | -24 | 50 | 29% Postcentral Gyrus |
| | | | 3.90 | 42 | -24 | 66 | 44% Postcentral Gyrus |
| | | | 3.83 | 36 | -20 | 60 | 30% Precentral Gyrus |
| | | | 3.65 | 46 | -18 | 58 | 41% Postcentral Gyrus |
| 6 | 99 | 0.034 | 4.13 | -30 | -10 | -18 | 33% Left Amygdala, 30% Left Hippocampus |
| | | | 4.09 | -36 | -20 | -14 | 29% Left Hippocampus |
| | | | 4.03 | -24 | -12 | -20 | 93% Left Hippocampus |
| 7 | 95 | 0.040 | 4.29 | -64 | -10 | -12 | 33% Middle Temporal Gyrus, anterior division, 29% Middle Temporal Gyrus, posterior division |
| | | | 3.66 | -58 | -18 | -20 | 39% Middle Temporal Gyrus, posterior division |
| | | | 3.65 | -64 | -12 | -20 | 47% Middle Temporal Gyrus, posterior division, 31% Middle Temporal Gyrus, anterior division |
| | | | 3.62 | -62 | -10 | -24 | 44% Middle Temporal Gyrus, anterior division, 34% Middle Temporal Gyrus, posterior division |
| *Sham group, acquisition training, CS+ > CS-* | | | | | | | |
| 1 | 2818 | < 0.001 | 5.76 | -10 | 22 | 32 | 44% Paracingulate Gyrus |
| | | | 5.24 | -8 | 32 | 20 | 49% Cingulate Gyrus, anterior division |
| | | | 5.22 | -6 | 20 | 36 | 45% Paracingulate Gyrus, 34% Cingulate Gyrus, anterior division |
| | | | 5.12 | 6 | -8 | 64 | 51% Juxtapositional Lobule Cortex (formerly Supplementary Motor Cortex) |

| | | | | | | | |
|---|---|---|---|---|---|---|---|
| | | | 4.90 | 6 | -4 | 70 | 50% Juxtapositional Lobule Cortex (formerly Supplementary Motor Cortex) |
| | | | 4.87 | -10 | 12 | 34 | 28% Cingulate Gyrus, anterior division, |
| 2 | 953 | < 0.001 | 5.30 | -14 | 8 | -4 | 48% Left Cerebral White Matter, 30% Left Putamen |
| | | | 4.92 | -42 | 22 | 2 | 66% Frontal Operculum Cortex |
| | | | 4.75 | -14 | 6 | 10 | 60% Left Cerebral White Matter, 40% Left Caudate |
| | | | 4.72 | -30 | 24 | -12 | 52% Frontal Orbital Cortex |
| | | | 4.66 | -30 | 12 | -14 | 44% Insular Cortex |
| 3 | 836 | < 0.001 | 5.57 | 44 | 24 | -4 | 36% Frontal Operculum Cortex, 36% Frontal Orbital Cortex |
| | | | 5.15 | 38 | 28 | -4 | 51% Frontal Orbital Cortex |
| | | | 4.32 | 60 | 10 | 6 | 33% Precentral Gyrus, 25% Inferior Frontal Gyrus, pars opercularis |
| 4 | 639 | < 0.001 | 5.19 | 10 | -6 | 6 | 99% Right Thalamus |
| | | | 5.07 | 10 | 8 | 8 | 85% Right Caudate |
| | | | 4.80 | -12 | -10 | 8 | 99% Left Thalamus |
| | | | 4.69 | 10 | 18 | -4 | 70% Right Accumbens |
| | | | 4.46 | 8 | 10 | 0 | 81% Right Caudate |
| 5 | 596 | < 0.001 | 5.66 | 50 | 0 | 40 | 39% Precentral Gyrus |
| | | | 5.66 | 46 | 2 | 54 | 36% Middle Frontal Gyrus, 30% Precentral Gyrus |
| | | | 5.63 | 48 | 2 | 44 | 60% Precentral Gyrus |
| | | | 5.18 | 50 | -2 | 48 | 66% Precentral Gyrus |
| | | | 4.15 | 48 | -12 | 46 | 45% Precentral Gyrus, 25% Postcentral Gyrus |
| | | | 3.85 | 34 | -6 | 48 | 38% Precentral Gyrus |
| 6 | 319 | < 0.001 | 5.81 | -36 | -52 | -34 | 68% Left Crus I, 32% Left VI |
| | | | 4.07 | -46 | -60 | -32 | 100% Left Crus I |
| | | | 4.03 | -32 | -56 | -28 | 95% Left VI |
| | | | 3.92 | -48 | -64 | -34 | 95% Left Crus I |
| | | | 3.37 | -50 | -50 | -34 | 78% Left Crus I |
| 7 | 242 | < 0.001 | 4.25 | 16 | -64 | -52 | 58% Right VIIIa, 37% Right VIIIb |
| | | | 4.24 | 12 | -70 | -44 | 63% Right VIIb |
| | | | 4.20 | 18 | -68 | -48 | 58% Right VIIb, 32% Right VIIIa |
| | | | 4.08 | 10 | -52 | -58 | 68% Right IX, 32% Right VIIIb |
| | | | 3.41 | 10 | -64 | -54 | 35% Right VIIIb, 31% Right VIIIa, 29% Right IX |
| 8 | 219 | 0.001 | 4.09 | -36 | 0 | 0 | 99% Right Thalamus |
| | | | 3.95 | -34 | -6 | -10 | 85% Right Caudate |
| | | | 3.85 | -26 | -8 | 0 | 99% Left Thalamus |
| | | | 3.82 | -30 | -12 | -2 | 70% Right Accumbens |
| | | | 3.80 | -36 | -12 | -6 | 48% Insular Cortex |
| 9 | 217 | 0.001 | 4.23 | 26 | -56 | -24 | 98% Right VI |
| | | | 4.22 | 22 | -54 | -24 | 86% Right VI |
| | | | 3.92 | 32 | -52 | -30 | 91% Right VI |
| | | | 3.87 | 42 | -56 | -34 | 95% Right Crus I |
| | | | 3.80 | 16 | -54 | -24 | 76% Right V |
| | | | 3.79 | 42 | -52 | -34 | 100% Right Crus I |
| 10 | 212 | 0.001 | 4.52 | -38 | -60 | -56 | 60% Left VIIb, 25% Left VIIIa |
| | | | 4.37 | -32 | -60 | -50 | 65% Left VIIb |
| | | | 4.17 | -26 | -66 | -56 | 66% Left VIIIa, 34% Left VIIb |
| | | | 3.70 | -36 | -50 | -50 | 45% Left VIIb, 35% Left VIIIa |
| 11 | 197 | 0.002 | 4.62 | -50 | 10 | 2 | 33% Inferior Frontal Gyrus, pars opercularis |
| | | | 4.33 | -54 | 6 | 6 | 56% Precentral Gyrus |
| | | | 4.08 | -44 | 6 | 6 | 54% Central Opercular Cortex |
| | | | 3.63 | -40 | 10 | -2 | 73% Insular Cortex |
| | | | 3.49 | -56 | 2 | -2 | 47% Planum Polare |
| 12 | 177 | 0.003 | 4.93 | 62 | -38 | 20 | 49% Supramarginal Gyrus, posterior division |
| | | | 3.33 | 64 | -48 | 26 | 56% Angular Gyrus |
| 13 | 177 | 0.003 | 4.65 | 54 | -22 | 20 | 46% Parietal Operculum Cortex |
| | | | 3.73 | 48 | -32 | 22 | 60% Parietal Operculum Cortex |
| | | | 3.71 | 50 | -30 | 18 | 47% Parietal Operculum Cortex, 27% Planum Temporale |
| | | | 3.69 | 58 | -18 | 32 | 39% Postcentral Gyrus, 34% Supramarginal Gyrus, anterior division |
| 14 | 138 | 0.012 | 5.09 | -58 | -16 | 30 | 65% Postcentral Gyrus |
| | | | 4.03 | -40 | -26 | 22 | 42% Parietal Operculum Cortex |
| | | | 4.02 | -50 | -24 | 20 | 39% Parietal Operculum Cortex, 31% Central Opercular Cortex |
| | | | 3.98 | -46 | -22 | 20 | 50% Central Opercular Cortex, 27% Parietal Operculum Cortex |
| | | | 3.58 | -56 | -22 | 22 | 29% Postcentral Gyrus |
| 15 | 120 | 0.023 | 4.20 | 14 | -44 | 48 | 36% Precuneal Cortex |

| | | | | | | | |
|---|---|---|---|---|---|---|---|
| | | | 3.66 | 6 | -50 | 60 | 66% Precuneal Cortex |
| | | | 3.59 | 6 | -44 | 58 | 52% Precuneal Cortex |
| 16 | 100 | 0.050 | 4.24 | -42 | -14 | 64 | 59% Precentral Gyrus |
| | | | 3.88 | -48 | -10 | 52 | 72% Precentral Gyrus |
| | | | 3.66 | -42 | 0 | 52 | 32% Middle Frontal Gyrus, 28% Precentral Gyrus |
| | | | 3.64 | -36 | -14 | 60 | 38% Precentral Gyrus |
| | | | 3.59 | -42 | -8 | 50 | 68% Precentral Gyrus |
| | | | 3.40 | -34 | -6 | 60 | 38% Precentral Gyrus |
| *Sham group, acquisition training, no-US post CS+ > no-US post CS-* | | | | | | | |
| 1 | 3239 | < 0.001 | 7.65 | 44 | 22 | -6 | 43% Frontal Orbital Cortex |
| | | | 7.06 | 30 | 24 | -10 | 64% Frontal Orbital Cortex |
| | | | 6.73 | 46 | 14 | 0 | 43% Frontal Operculum Cortex |
| | | | 6.68 | 46 | 20 | 0 | 42% Frontal Operculum Cortex |
| | | | 6.29 | 34 | 24 | 10 | 44% Frontal Operculum Cortex |
| | | | 5.81 | 54 | 6 | 46 | 32% Precentral Gyrus |
| 2 | 2169 | < 0.001 | 5.73 | 0 | 8 | 60 | 33% Juxtapositional Lobule Cortex (formerly Supplementary Motor Cortex) |
| | | | 5.54 | -4 | 18 | 38 | 52% Paracingulate Gyrus, 36% Cingulate Gyrus, anterior division |
| | | | 5.36 | -2 | 10 | 52 | 45% Paracingulate Gyrus |
| | | | 5.00 | 8 | 22 | 36 | 37% Paracingulate Gyrus, 28% Cingulate Gyrus, anterior division |
| | | | 4.87 | -2 | 20 | 56 | 65% Superior Frontal Gyrus |
| | | | 4.73 | 4 | 14 | 64 | 54% Superior Frontal Gyrus |
| 3 | 1732 | < 0.001 | 5.61 | -46 | 14 | -4 | 33% Frontal Operculum Cortex |
| | | | 5.28 | -46 | 20 | 8 | 25% Inferior Frontal Gyrus, pars opercularis |
| | | | 4.96 | -40 | 6 | -2 | 68% Insular Cortex |
| | | | 4.96 | -18 | 4 | 0 | 65% Left Pallidum, 32% Left Putamen |
| | | | 4.83 | -36 | 20 | -8 | 41% Frontal Orbital Cortex, 30% Insular Cortex |
| | | | 4.76 | -28 | 18 | -10 | 35% Insular Cortex, |
| 4 | 1353 | < 0.001 | 5.59 | 66 | -38 | 26 | 57% Supramarginal Gyrus, posterior division |
| | | | 5.32 | 62 | -42 | 34 | 62% Supramarginal Gyrus, posterior division |
| | | | 5.21 | 60 | -52 | 4 | 65% Middle Temporal Gyrus, temporooccipital part |
| | | | 4.90 | 58 | -50 | 20 | 72% Angular Gyrus |
| | | | 4.89 | 58 | -50 | 24 | 65% Angular Gyrus |
| | | | 4.85 | 56 | -20 | 20 | 27% Parietal Operculum Cortex |
| 5 | 383 | < 0.001 | 4.90 | -56 | -48 | 28 | 44% Supramarginal Gyrus, posterior division |
| | | | 4.74 | -60 | -46 | 28 | 53% Supramarginal Gyrus, posterior division |
| | | | 4.67 | -64 | -42 | 32 | 42% Supramarginal Gyrus, posterior division, 28% Supramarginal Gyrus, anterior division |
| | | | 4.65 | -62 | -50 | 30 | 51% Supramarginal Gyrus, posterior division, 29% Angular Gyrus |
| 6 | 382 | < 0.001 | 4.42 | 4 | -24 | 0 | 45% Right Thalamus |
| | | | 4.32 | 6 | -16 | -16 | [Right Substantia Nigra] |
| | | | 4.27 | -6 | -16 | -10 | [Left Red Nucleus] |
| | | | 4.16 | -12 | -6 | -16 | 27% Left Amygdala |
| | | | 3.98 | 8 | -8 | -8 | [Right Subthalamic Nucleus] |
| | | | 3.90 | -10 | -10 | -10 | [Left Subthalamic Nucleus] |
| 7 | 263 | < 0.001 | 4.43 | 30 | -56 | -24 | 100% Right VI |
| | | | 4.38 | 34 | -56 | -26 | 95% Right VI |
| | | | 4.38 | 40 | -52 | -30 | 70% Right Crus I, 25% Right VI |
| | | | 4.21 | 24 | -60 | -22 | 96% Right VI |
| | | | 4.06 | 24 | -64 | -16 | 45% Right VI |
| | | | 3.73 | 24 | -78 | -10 | 59% Occipital Fusiform Gyrus |
| 8 | 261 | < 0.001 | 4.76 | -48 | -2 | 44 | 46% Precentral Gyrus |
| | | | 3.74 | -46 | 2 | 56 | 33% Middle Frontal Gyrus |
| 9 | 247 | < 0.001 | 4.54 | -14 | -30 | 42 | 39% Precentral Gyrus |
| | | | 4.36 | 14 | -42 | 50 | 33% Precuneal Cortex |
| | | | 4.12 | -12 | -38 | 54 | 27% Postcentral Gyrus |
| | | | 4.03 | 6 | -42 | 54 | 52% Precuneal Cortex |
| | | | 3.93 | -4 | -46 | 58 | 59% Precuneal Cortex |
| 10 | 216 | < 0.001 | 4.90 | 46 | -30 | 2 | 39% Superior Temporal Gyrus, posterior division |
| | | | 4.73 | 52 | -20 | -4 | 53% Superior Temporal Gyrus, posterior division |
| | | | 4.34 | 46 | -24 | -6 | 32% Middle Temporal Gyrus, posterior division, 31% Superior Temporal Gyrus, posterior division |
| 11 | 156 | 0.003 | 4.54 | -50 | -50 | 56 | 32% Supramarginal Gyrus, posterior division |
| | | | 4.52 | -28 | -52 | 44 | 41% Superior Parietal Lobule |
| | | | 3.41 | -36 | -52 | 52 | 45% Superior Parietal Lobule |

| | | | | | | | |
|---|---|---|---|---|---|---|---|
| 12 | 122 | 0.012 | 4.63 | -52 | -52 | 10 | 39% Middle Temporal Gyrus, temporooccipital part |
| | | | 4.08 | -58 | -60 | 14 | 37% Angular Gyrus, 28% Lateral Occipital Cortex, superior division |
| | | | 3.77 | -54 | -58 | 12 | 31% Middle Temporal Gyrus, temporooccipital part, 26% Angular Gyrus |
| *Verum group, acquisition training, CS+ > CS-* | | | | | | | |
| 1 | 2666 | < 0.001 | 7.09 | -4 | 6 | 48 | 51% Juxtapositional Lobule Cortex (formerly Supplementary Motor Cortex) |
| | | | 6.97 | 2 | -6 | 66 | 63% Juxtapositional Lobule Cortex (formerly Supplementary Motor Cortex) |
| | | | 6.30 | -14 | 2 | 72 | 44% Superior Frontal Gyrus |
| | | | 6.05 | -6 | -4 | 70 | 34% Juxtapositional Lobule Cortex (formerly Supplementary Motor Cortex) |
| | | | 5.74 | -16 | 4 | 66 | 49% Superior Frontal Gyrus |
| 2 | 1716 | < 0.001 | 6.23 | -20 | -44 | 64 | 38% Postcentral Gyrus |
| | | | 6.12 | 22 | -42 | 70 | 31% Postcentral Gyrus, 29% Superior Parietal Lobule |
| | | | 5.52 | 16 | -50 | 72 | 52% Superior Parietal Lobule |
| | | | 5.41 | 2 | -46 | 54 | 65% Precuneal Cortex |
| | | | 5.12 | -18 | -42 | 74 | 49% Postcentral Gyrus |
| | | | 5.07 | -8 | -42 | 52 | 47% Precuneal Cortex, 26% Postcentral Gyrus |
| 3 | 702 | < 0.001 | 6.52 | 8 | 10 | 0 | 81% Right Caudate |
| | | | 6.30 | -8 | 8 | 0 | 73% Left Caudate |
| | | | 5.48 | 10 | 8 | 8 | 91% Right Caudate |
| | | | 4.67 | -18 | 14 | -2 | 60% Left Putamen, 36% Left Cerebral White Matter |
| | | | 4.34 | -6 | 22 | 4 | 52% Left Lateral Ventricle, 47% Left Cerebral White Matter |
| | | | 4.33 | -8 | -2 | 10 | 57% Left Thalamus |
| 4 | 473 | < 0.001 | 6.19 | -30 | 24 | 8 | 35% Insular Cortex |
| | | | 4.43 | -48 | 10 | -2 | 27% Frontal Operculum Cortex |
| | | | 4.16 | -34 | 12 | 10 | 39% Frontal Operculum Cortex |
| | | | 4.10 | -54 | 8 | -2 | 25% Temporal Pole |
| | | | 3.96 | -50 | 2 | 8 | 35% Central Opercular Cortex |
| | | | 3.69 | -50 | 8 | 12 | 35% Inferior Frontal Gyrus, pars opercularis |
| 5 | 466 | < 0.001 | 5.12 | -64 | -36 | 30 | 65% Supramarginal Gyrus, anterior division |
| | | | 5.07 | -56 | -38 | 36 | 42% Supramarginal Gyrus, anterior division |
| | | | 4.50 | -66 | -26 | 22 | 48% Supramarginal Gyrus, anterior division |
| | | | 4.48 | -56 | -38 | 28 | 32% Parietal Operculum Cortex |
| | | | 4.30 | -50 | -36 | 26 | 50% Parietal Operculum Cortex |
| 6 | 376 | < 0.001 | 5.78 | -42 | 46 | 26 | 52% Frontal Pole |
| | | | 4.73 | -22 | 44 | 20 | 35% Frontal Pole |
| | | | 4.46 | -34 | 52 | 24 | 83% Frontal Pole |
| | | | 4.25 | -30 | 42 | 24 | 54% Frontal Pole |
| | | | 3.86 | -32 | 40 | 34 | 53% Frontal Pole, 26% Middle Frontal Gyrus |
| | | | 3.78 | -42 | 36 | 32 | 49% Middle Frontal Gyrus, 28% Frontal Pole |
| 7 | 333 | < 0.001 | 4.50 | 50 | 4 | 48 | 54% Precentral Gyrus |
| | | | 4.40 | 48 | 2 | 54 | 33% Precentral Gyrus |
| | | | 4.34 | 50 | -2 | 50 | 70% Precentral Gyrus |
| | | | 4.32 | 38 | -4 | 46 | 30% Precentral Gyrus |
| | | | 4.21 | 52 | 4 | 40 | 47% Precentral Gyrus |
| | | | 4.04 | 44 | -12 | 44 | 38% Precentral Gyrus |
| 8 | 325 | < 0.001 | 5.54 | -40 | -4 | 54 | 44% Precentral Gyrus, 25% Middle Frontal Gyrus |
| | | | 4.40 | -48 | -4 | 46 | 47% Precentral Gyrus |
| | | | 3.76 | -30 | -10 | 50 | 41% Precentral Gyrus |
| | | | 3.74 | -52 | 0 | 38 | 69% Precentral Gyrus |
| 9 | 242 | < 0.001 | 4.52 | -8 | -74 | 46 | 55% Precuneal Cortex |
| | | | 4.22 | -12 | -78 | 42 | 32% Lateral Occipital Cortex, superior division |
| | | | 4.18 | -10 | -78 | 36 | 39% Precuneal Cortex, 29% Cuneal Cortex |
| | | | 4.16 | -6 | -80 | 48 | 32% Precuneal Cortex |
| 10 | 205 | 0.001 | 5.37 | 36 | 26 | 8 | 34% Frontal Operculum Cortex |
| | | | 3.93 | 34 | 16 | 8 | 36% Insular Cortex, 31% Frontal Operculum Cortex |
| 11 | 116 | 0.027 | 4.10 | 16 | -70 | 36 | 40% Precuneal Cortex |
| | | | 3.84 | 16 | -70 | 44 | 29% Precuneal Cortex |
| | | | 3.76 | 12 | -76 | 48 | 46% Lateral Occipital Cortex, superior division |
| 12 | 115 | 0.028 | 4.17 | -32 | -54 | 44 | 35% Superior Parietal Lobule |
| 13 | 108 | 0.036 | 4.45 | -42 | -58 | -26 | 61% Left Crus I |
| | | | 4.30 | -34 | -56 | -26 | 92% Left VI |

| | | | | | | | |
|---|---|---|---|---|---|---|---|
| *Verum group, acquisition training, CS+ < CS-* | | | | | | | |
| 1 | 142 | 0.010 | 4.32 | 20 | 34 | 56 | 43% Superior Frontal Gyrus |
| | | | 4.32 | 24 | 30 | 52 | 48% Superior Frontal Gyrus |
| | | | 4.27 | 26 | 26 | 52 | 35% Superior Frontal Gyrus |
| | | | 4.05 | 20 | 38 | 48 | 44% Frontal Pole |
| 2 | 140 | 0.011 | 4.33 | 52 | -60 | 36 | 51% Lateral Occipital Cortex, superior division |
| | | | 4.26 | 48 | -56 | 36 | 43% Angular Gyrus |
| | | | 3.40 | 48 | -70 | 34 | 87% Lateral Occipital Cortex, superior division |
| 3 | 106 | 0.039 | 4.71 | 50 | 36 | -12 | 61% Frontal Pole |
| *Verum group, acquisition training, no-US post CS+ > no-US post CS-* | | | | | | | |
| 1 | 2695 | < 0.001 | 6.67 | 42 | 20 | 0 | 54% Frontal Operculum Cortex |
| | | | 5.88 | 58 | 14 | 2 | 35% Inferior Frontal Gyrus, pars opercularis |
| | | | 5.85 | 42 | 10 | 4 | 37% Central Opercular Cortex, 29% Frontal Operculum Cortex |
| | | | 5.22 | 52 | 12 | 24 | 43% Inferior Frontal Gyrus, pars opercularis, 26% Precentral Gyrus |
| 2 | 1093 | < 0.001 | 6.65 | -32 | 26 | 0 | 32% Frontal Orbital Cortex, 27% Insular Cortex |
| | | | 5.35 | -40 | 14 | -4 | 73% Insular Cortex |
| | | | 5.34 | -38 | 10 | 12 | 33% Frontal Operculum Cortex |
| | | | 5.12 | -50 | 0 | 2 | 69% Central Opercular Cortex |
| | | | 4.42 | -56 | 10 | 2 | 43% Inferior Frontal Gyrus, pars opercularis, 25% Precentral Gyrus |
| 3 | 659 | < 0.001 | 5.13 | -10 | 2 | 42 | 39% Juxtapositional Lobule Cortex (formerly Supplementary Motor Cortex), 28% Cingulate Gyrus, anterior division |
| | | | 4.99 | 4 | 10 | 38 | 77% Cingulate Gyrus, anterior division |
| | | | 4.44 | 8 | 26 | 30 | 36% Paracingulate Gyrus, 34% Cingulate Gyrus, anterior division |
| | | | 3.97 | 2 | 32 | 26 | 54% Cingulate Gyrus, anterior division, 28% Paracingulate Gyrus |
| | | | 3.84 | 12 | 18 | 30 | 26% Cingulate Gyrus, anterior division |
| | | | 3.72 | -10 | 4 | 52 | 38% Juxtapositional Lobule Cortex (formerly Supplementary Motor Cortex) |
| 4 | 590 | < 0.001 | 4.78 | -12 | -74 | 14 | 39% Intracalcarine Cortex |
| | | | 4.41 | 22 | -64 | 8 | 46% Intracalcarine Cortex |
| | | | 4.09 | -4 | -92 | 8 | 47% Occipital Pole |
| | | | 4.08 | -6 | -86 | 10 | 28% Intracalcarine Cortex |
| | | | 4.05 | -18 | -68 | -2 | 25% Lingual Gyrus |
| | | | 3.98 | -18 | -70 | 8 | 52% Intracalcarine Cortex |
| 5 | 447 | < 0.001 | 5.03 | 8 | -4 | 72 | 28% Juxtapositional Lobule Cortex (formerly Supplementary Motor Cortex), 28% Superior Frontal Gyrus |
| | | | 4.52 | 4 | 0 | 74 | 23% Juxtapositional Lobule Cortex (formerly Supplementary Motor Cortex) |
| | | | 4.47 | 2 | -6 | 66 | 63% Juxtapositional Lobule Cortex (formerly Supplementary Motor Cortex) |
| | | | 4.18 | 4 | 6 | 60 | 63% Juxtapositional Lobule Cortex (formerly Supplementary Motor Cortex) |
| | | | 4.05 | 14 | -8 | 72 | 38% Superior Frontal Gyrus |
| 6 | 322 | < 0.001 | 4.40 | 52 | -30 | 28 | 33% Parietal Operculum Cortex |
| | | | 3.94 | 66 | -32 | 22 | 30% Superior Temporal Gyrus, posterior division |
| | | | 3.92 | 64 | -26 | 24 | 52% Supramarginal Gyrus, anterior division |
| | | | 3.82 | 66 | -38 | 30 | 62% Supramarginal Gyrus, posterior division |
| | | | 3.67 | 66 | -42 | 20 | 57% Supramarginal Gyrus, posterior division |
| | | | 3.62 | 60 | -46 | 12 | 33% Middle Temporal Gyrus, temporooccipital part, 27% Angular Gyrus |
| 7 | 232 | < 0.001 | 5.24 | -30 | -58 | -28 | 100% Left VI |
| | | | 3.79 | -46 | -60 | -30 | 100% Left Crus I |
| | | | 3.57 | -40 | -62 | -26 | 95% Left Crus I |
| 8 | 229 | < 0.001 | 4.38 | 46 | -20 | -8 | 31% Superior Temporal Gyrus, posterior division |
| | | | 4.36 | 50 | -26 | 0 | 58% Superior Temporal Gyrus, posterior division |
| | | | 4.28 | 50 | -22 | -6 | 42% Superior Temporal Gyrus, posterior division, 41% Middle Temporal Gyrus, posterior division |
| | | | 4.25 | 54 | -22 | -6 | 46% Middle Temporal Gyrus, posterior division, 38% Superior Temporal Gyrus, posterior division |
| | | | 3.85 | 58 | -30 | 0 | 55% Superior Temporal Gyrus, posterior division |
| | | | 3.79 | 62 | -32 | -4 | 52% Middle Temporal Gyrus, posterior division |
| 9 | 212 | < 0.001 | 5.00 | 40 | -54 | 46 | 36% Angular Gyrus |
| | | | 3.79 | 42 | -42 | 40 | 36% Supramarginal Gyrus, posterior division |
| | | | 3.71 | 52 | -44 | 38 | 36% Supramarginal Gyrus, posterior division, 26% Angular Gyrus |
| | | | 3.67 | 38 | -48 | 46 | 33% Superior Parietal Lobule |

| | | | | | | | |
|---|---|---|---|---|---|---|---|
| 10 | 200 | 0.001 | 4.58 | -50 | -32 | 26 | 33% Parietal Operculum Cortex |
| | | | 4.18 | -58 | -30 | 22 | 45% Parietal Operculum Cortex |
| | | | 3.62 | -64 | -22 | 20 | 47% Postcentral Gyrus |
| | | | 3.36 | -66 | -36 | 24 | 42% Supramarginal Gyrus, anterior division |
| *Verum group, acquisition training, no-US post CS+ < no-US post CS-* | | | | | | | |
| 1 | 625 | < 0.001 | 5.25 | -4 | -50 | 22 | 67% Cingulate Gyrus, posterior division |
| | | | 4.84 | -6 | -56 | 26 | 37% Precuneal Cortex, 37% Cingulate Gyrus, posterior division |
| | | | 4.61 | -4 | -48 | 14 | 71% Cingulate Gyrus, posterior division, |
| | | | 4.52 | -8 | -58 | 14 | 53% Precuneal Cortex |
| | | | 4.07 | -2 | -62 | 24 | 86% Precuneal Cortex |
| 2 | 387 | < 0.001 | 4.53 | 4 | 38 | -16 | 72% Frontal Medial Cortex |
| | | | 4.51 | -6 | 58 | -6 | 64% Frontal Pole |
| | | | 4.07 | 8 | 48 | -16 | 30% Frontal Medial Cortex |
| | | | 4.00 | -4 | 50 | -18 | 73% Frontal Medial Cortex |
| | | | 3.62 | -2 | 64 | -14 | 70% Frontal Pole |
| 3 | 237 | < 0.001 | 5.05 | 38 | -74 | -36 | 81% Right Crus I |
| | | | 4.26 | 46 | -72 | -46 | 55% Right Crus II |
| | | | 4.05 | 34 | -84 | -28 | 88% Right Crus I |
| | | | 3.79 | 26 | -84 | -32 | 50% Right Crus I, 45% Right Crus II |
| | | | 3.74 | 48 | -70 | -40 | 61% Right Crus I, 29% Right Crus II |
| | | | 3.39 | 22 | -90 | -28 | 28% Right Crus II, 25% Right Crus I |
| 4 | 90 | 0.049 | 4.21 | -46 | -72 | 40 | 87% Lateral Occipital Cortex, superior division |
| | | | 4.13 | -46 | -72 | 32 | 77% Lateral Occipital Cortex, superior division |
| *Acquisition training, CS+ > CS-, verum > sham* | | | | | | | |
| | no surviving clusters | | | | | | |
| *Acquisition training, CS+ > CS-, verum < sham* | | | | | | | |
| | no surviving clusters | | | | | | |
| *Acquisition training, no-US post CS+ > no-US post CS-, verum > sham* | | | | | | | |
| | no surviving clusters | | | | | | |
| *Acquisition training, no-US post CS+ > no-US post CS-, verum < sham* | | | | | | | |
| | no surviving clusters | | | | | | |

**Table 4-2.** Fear extinction training. Activation clusters are reported. Listed below are all local maxima within significant clusters ($p < 0.05$, corrected for multiple comparisons using a cluster-forming threshold of $Z = 3.1$). The anatomical locations of these maxima were determined by referencing the Harvard-Oxford Cortical and Subcortical, and Cerebellar Atlases in MNI152 space following normalization with FLIRT. Only structures with a probability of 25% or higher are reported. For regions where the Harvard-Oxford atlas was insufficient for identification, manual identification was conducted. Brain areas manually identified using the Multi-Contrast Anatomical Subcortical Structures (MASSP) atlas (Bazin et al., 2020) are indicated in square brackets. Percentages represent the likelihood of anatomical localization.

| Index | Cluster size / voxel | *p* value | Z | Coordinates / mm | | | Name (Harvard-Oxford Atlas) |
|---|---|---|---|---|---|---|---|
| | | | | x | y | z | |
| *Sham group, extinction training, CS+ > CS-* | | | | | | | |
| 1 | 134 | 0.009 | 4.50 | -62 | -42 | 28 | 45% Supramarginal Gyrus, posterior division |
| | | | 4.40 | -62 | -44 | 32 | 58% Supramarginal Gyrus, posterior division |
| | | | 4.34 | -58 | -40 | 30 | 29% Supramarginal Gyrus, anterior division |
| *Sham group, extinction training, no-US post CS+ > no-US post CS-* | | | | | | | |
| 1 | 505 | < 0.001 | 4.54 | 48 | 28 | -4 | 29% Frontal Orbital Cortex |
| | | | 4.29 | 52 | 8 | 6 | 35% Precentral Gyrus |
| | | | 4.26 | 56 | 18 | 10 | 64% Inferior Frontal Gyrus, pars opercularis |
| | | | 4.22 | 58 | 24 | 4 | 38% Inferior Frontal Gyrus, pars triangularis |
| | | | 4.21 | 48 | 10 | 12 | 29% Inferior Frontal Gyrus, pars opercularis |
| 2 | 110 | 0.030 | 4.23 | 2 | 34 | 48 | 61% Superior Frontal Gyrus |
| *Verum group, extinction training, CS+ > CS-* | | | | | | | |
| 1 | 454 | < 0.001 | 4.55 | 10 | 2 | 48 | 57% Juxtapositional Lobule Cortex (formerly Supplementary Motor Cortex) |
| | | | 4.41 | 6 | 18 | 44 | 66% Paracingulate Gyrus |
| | | | 4.38 | -6 | 8 | 48 | 26% Paracingulate Gyrus, 26% Juxtapositional Lobule Cortex (formerly Supplementary Motor Cortex) |
| | | | 4.17 | 2 | 12 | 36 | 81% Cingulate Gyrus, anterior division |
| | | | 4.04 | 10 | 10 | 48 | 33% Paracingulate Gyrus |
| | | | 3.92 | 4 | 20 | 28 | 84% Cingulate Gyrus, anterior division |
| 2 | 300 | < 0.001 | 4.66 | 54 | 12 | 2 | 40% Inferior Frontal Gyrus, pars opercularis |
| | | | 4.15 | 46 | 12 | 6 | 34% Frontal Operculum Cortex |
| | | | 3.88 | 54 | 22 | 14 | 30% Inferior Frontal Gyrus, pars opercularis |
| | | | 3.78 | 56 | 4 | 18 | 46% Precentral Gyrus |
| | | | 3.51 | 48 | 12 | 12 | 30% Inferior Frontal Gyrus, pars opercularis |
| 3 | 215 | < 0.001 | 5.26 | 8 | 38 | 16 | 57% Cingulate Gyrus, anterior division |
| | | | 4.23 | 8 | 34 | 22 | 44% Cingulate Gyrus, anterior division, 32% Paracingulate Gyrus |
| | | | 4.00 | 0 | 32 | 34 | 68% Paracingulate Gyrus |
| | | | 3.73 | 0 | 42 | 42 | 41% Superior Frontal Gyrus |
| | | | 3.45 | 2 | 38 | 16 | 78% Cingulate Gyrus, anterior division |
| | | | 3.42 | 8 | 46 | 24 | 52% Paracingulate Gyrus |
| 4 | 171 | 0.002 | 4.40 | 4 | -26 | 44 | 78% Cingulate Gyrus, posterior division |
| | | | 4.00 | 14 | -30 | 46 | 41% Precentral Gyrus |
| | | | 3.97 | 4 | -20 | 44 | 65% Cingulate Gyrus, posterior division |
| | | | 3.78 | 10 | -20 | 40 | 52% Cingulate Gyrus, posterior division |
| 5 | 119 | 0.016 | 4.06 | -14 | -80 | 40 | 33% Lateral Occipital Cortex, superior division |
| | | | 3.87 | -20 | -76 | 42 | 53% Lateral Occipital Cortex, superior division |
| | | | 3.64 | -14 | -76 | 30 | 39% Cuneal Cortex, 26% Precuneal Cortex |
| | | | 3.49 | -16 | -86 | 36 | 36% Lateral Occipital Cortex, superior division |
| | | | 3.48 | -22 | -84 | 38 | 60% Lateral Occipital Cortex, superior division |
| | | | 3.34 | -10 | -84 | 34 | 39% Cuneal Cortex |
| 6 | 109 | 0.025 | 4.39 | -8 | -102 | 16 | 52% Occipital Pole |
| | | | 3.64 | -4 | -90 | 14 | 26% Occipital Pole |
| | | | 3.45 | 8 | -82 | 22 | 39% Cuneal Cortex |
| | | | 3.44 | -2 | -86 | 26 | 57% Cuneal Cortex |
| | | | 3.29 | 2 | -88 | 18 | 37% Cuneal Cortex |
| 7 | 100 | 0.036 | 4.45 | -20 | -62 | 68 | 44% Lateral Occipital Cortex, superior division |
| | | | 3.69 | -14 | -54 | 70 | 49% Superior Parietal Lobule |
| 8 | 99 | 0.038 | 4.30 | 26 | 22 | -14 | 60% Frontal Orbital Cortex |
| | | | 3.88 | 28 | 24 | -6 | 30% Frontal Orbital Cortex |

| | | | | | | | |
|---|---|---|---|---|---|---|---|
| | | | 3.39 | 38 | 28 | -6 | 56% Frontal Orbital Cortex |
| 9 | 94 | 0.047 | 4.43 | 22 | -48 | 72 | 64% Superior Parietal Lobule |
| *Verum group, extinction training, no-US post CS+ > no-US post CS-* | | | | | | | |
| | no surviving clusters | | | | | | |
| *Extinction training, CS+ > CS-, verum > sham* | | | | | | | |
| 1 | 602 | < 0.001 | 4.45 | 10 | -82 | 22 | 35% Cuneal Cortex |
| | | | 4.44 | -6 | -100 | 16 | 62% Occipital Pole |
| | | | 4.03 | -4 | -90 | 14 | 26% Occipital Pole |
| | | | 4.01 | -8 | -86 | 24 | 30% Cuneal Cortex |
| | | | 3.99 | 18 | -90 | 28 | 42% Occipital Pole |
| | | | 3.93 | -6 | -78 | 24 | 42% Cuneal Cortex |
| 2 | 94 | 0.047 | 4.79 | -6 | -74 | -4 | 78% Lingual Gyrus |
| | | | 3.72 | -16 | -78 | -6 | 26% Lingual Gyrus |
| *Extinction training, CS+ > CS-, verum < sham* | | | | | | | |
| | no surviving clusters | | | | | | |
| *Extinction training, no-US post CS+ > no-US post CS-, verum > sham* | | | | | | | |
| | no surviving clusters | | | | | | |
| *Extinction training, no-US post CS+ > no-US post CS-, verum < sham* | | | | | | | |
| | no surviving clusters | | | | | | |

**Table 4-3.** Recall. Activation clusters are reported. Listed below are all local maxima within significant clusters ($p < 0.05$, corrected for multiple comparisons using a cluster-forming threshold of $Z = 3.1$). The anatomical locations of these maxima were determined by referencing the Harvard-Oxford Cortical and Subcortical, and Cerebellar Atlases in MNI152 space following normalization with FLIRT. Only structures with a probability of 25% or higher are reported. For regions where the Harvard-Oxford atlas was insufficient for identification, manual identification was conducted. Brain areas manually identified using the Multi-Contrast Anatomical Subcortical Structures (MASSP) atlas (Bazin et al., 2020) are indicated in square brackets. Percentages represent the likelihood of anatomical localization.

| Index | Cluster size / voxel | *p* value | Z | Coordinates / mm | | | Name (Harvard-Oxford Atlas) |
|---|---|---|---|---|---|---|---|
| | | | | x | y | z | |
| *Sham group, recall, CS+ > CS-* | | | | | | | |
| 1 | 230 | < 0.001 | 4.43 | 46 | 18 | 2 | 47% Frontal Operculum Cortex |
| | | | 4.18 | 32 | 22 | 12 | 35% Frontal Operculum Cortex |
| | | | 3.38 | 40 | 18 | -4 | 65% Insular Cortex |
| *Sham group, recall, no-US post CS+ > no-US post CS-* | | | | | | | |
| 1 | 943 | < 0.001 | 5.12 | 4 | 28 | 30 | 52% Cingulate Gyrus, anterior division, 38% Paracingulate Gyrus |
| | | | 5.06 | 8 | 28 | 34 | 46% Paracingulate Gyrus, 27% Cingulate Gyrus, anterior division |
| | | | 4.57 | 6 | 16 | 42 | 61% Paracingulate Gyrus |
| | | | 4.51 | 8 | 34 | 28 | 52% Paracingulate Gyrus, 26% Cingulate Gyrus, anterior division |
| | | | 4.50 | 4 | 42 | 34 | 53% Paracingulate Gyrus, 26% Superior Frontal Gyrus |
| | | | 4.50 | 0 | 18 | 60 | 38% Superior Frontal Gyrus |
| 2 | 795 | < 0.001 | 5.60 | 62 | -46 | 38 | 42% Supramarginal Gyrus, posterior division, 31% Angular Gyrus |
| | | | 5.53 | 60 | -44 | 30 | 46% Supramarginal Gyrus, posterior division, 31% Angular Gyrus |
| | | | 5.38 | 64 | -44 | 32 | 51% Supramarginal Gyrus, posterior division, 26% Angular Gyrus |
| | | | 5.28 | 56 | -34 | 32 | 27% Parietal Operculum Cortex |
| | | | 5.20 | 64 | -36 | 30 | 44% Supramarginal Gyrus, posterior division |
| | | | 5.14 | 66 | -42 | 26 | 55% Supramarginal Gyrus, posterior division |
| 3 | 781 | < 0.001 | 7.27 | 42 | 24 | -4 | 42% Frontal Orbital Cortex, 31% Frontal Operculum Cortex |
| | | | 7.20 | 46 | 20 | -4 | 39% Frontal Operculum Cortex |
| | | | 4.10 | 28 | 20 | -12 | 51% Frontal Orbital Cortex |
| | | | 3.79 | 40 | 6 | 6 | 49% Central Opercular Cortex |
| | | | 3.55 | 38 | 18 | 12 | 31% Frontal Operculum Cortex |
| | | | 3.51 | 34 | 10 | 2 | 38% Insular Cortex |
| 4 | 566 | < 0.001 | 4.70 | -10 | -76 | -28 | 49% Left Crus I, 25% Left VI II |
| | | | 4.53 | -26 | -70 | -28 | 55% Left Crus I, 45% Left VI |
| | | | 4.36 | -14 | -82 | -30 | 57% Left Crus I, 38% Left Crus II |
| | | | 4.16 | -38 | -54 | -30 | 68% Left Crus I, 32% Left VI |
| | | | 4.12 | -18 | -66 | -26 | 95% Left VI |
| | | | 3.88 | -48 | -56 | -34 | 90% Left Crus I |
| 5 | 394 | < 0.001 | 5.54 | 52 | 10 | 46 | 40% Middle Frontal Gyrus |
| | | | 4.39 | 38 | 4 | 46 | 39% Middle Frontal Gyrus |
| | | | 4.37 | 44 | 8 | 58 | 36% Middle Frontal Gyrus |
| | | | 4.27 | 40 | 8 | 54 | 49% Middle Frontal Gyrus |
| | | | 4.23 | 42 | 0 | 38 | 36% Precentral Gyrus |
| | | | 4.13 | 36 | 6 | 52 | 37% Middle Frontal Gyrus |
| 6 | 352 | < 0.001 | 5.13 | -28 | 22 | -8 | 36% Frontal Orbital Cortex, 26% Insular Cortex |
| | | | 4.76 | -44 | 18 | -4 | 34% Frontal Operculum Cortex |
| | | | 3.72 | -34 | 24 | 6 | 49% Frontal Operculum Cortex |
| | | | 3.41 | -28 | 12 | -8 | [Left Putamen] |
| 7 | 248 | < 0.001 | 5.19 | -64 | -48 | 32 | 44% Supramarginal Gyrus, posterior division |
| | | | 5.17 | -62 | -50 | 36 | 43% Supramarginal Gyrus, posterior division |
| | | | 4.84 | -58 | -46 | 30 | 50% Supramarginal Gyrus, posterior division |
| 8 | 199 | 0.001 | 4.60 | -44 | 6 | 58 | 27% Middle Frontal Gyrus |
| | | | 4.57 | -46 | 2 | 42 | 41% Precentral Gyrus |
| | | | 4.51 | -52 | 6 | 42 | 37% Precentral Gyrus, 29% Middle Frontal Gyrus |
| 9 | 169 | 0.002 | 4.37 | 30 | 60 | 16 | 87% Frontal Pole |
| | | | 3.95 | 24 | 54 | 18 | 71% Frontal Pole |
| 10 | 154 | 0.004 | 4.84 | 46 | 34 | 36 | 34% Middle Frontal Gyrus |
| | | | 3.74 | 52 | 28 | 28 | 52% Middle Frontal Gyrus |

| | | | | | | | |
|---|---|---|---|---|---|---|---|
| | | | 3.18 | 50 | 28 | 18 | 34% Inferior Frontal Gyrus, pars triangularis |
| 11 | 111 | 0.022 | 4.16 | 46 | -58 | 50 | 38% Lateral Occipital Cortex, superior division, 29% Angular Gyrus |
| | | | 3.83 | 44 | -52 | 50 | 46% Angular Gyrus |
| 12 | 92 | 0.050 | 4.74 | 42 | -52 | -28 | 70% right crus I |
| | | | 4.06 | 42 | -66 | -26 | 100% right Crus I |
| | | | 3.24 | 34 | -68 | -24 | 62% right crus I, 38% right VI |
| *Verum group, recall, CS+ > CS-* | | | | | | | |
| | No surviving clusters | | | | | | |
| *Verum group, recall, no-US post CS+ > no-US post CS-* | | | | | | | |
| 1 | 314 | < 0.001 | 4.59 | 42 | 20 | -6 | 36% Frontal Orbital Cortex |
| | | | 4.55 | 48 | 20 | -8 | 26% Frontal Orbital Cortex |
| | | | 4.08 | 32 | 26 | -6 | 56% Frontal Orbital Cortex |
| | | | 3.98 | 28 | 18 | -12 | 32% Insular Cortex, 27% Frontal Orbital Cortex |
| | | | 3.55 | 34 | 24 | -14 | 68% Frontal Orbital Cortex |
| | | | 3.52 | 52 | 22 | 4 | 32% Inferior Frontal Gyrus, pars triangularis |
| *Recall, CS+ > CS-, verum > sham* | | | | | | | |
| | No surviving clusters | | | | | | |
| *Recall, CS+ > CS-, verum < sham* | | | | | | | |
| | no surviving clusters | | | | | | |
| *Recall, no-US post CS+ > no-US post CS-, verum > sham* | | | | | | | |
| | no surviving clusters | | | | | | |
| *Recall, no-US post CS+ > no-US post CS-, verum < sham* | | | | | | | |
| 1 | 126 | 0.012 | 4.58 | -54 | 6 | 42 | 30% Precentral Gyrus |
| | | | 4.39 | -44 | -2 | 38 | 47% Precentral Gyrus |
| | | | 4.36 | -44 | 0 | 42 | 40% Precentral Gyrus |

**Table 5-1.** Fear acquisition training. Parametric modulation of learning model-derived prediction parameters. Activation clusters are reported. Listed below are all local maxima within significant clusters ($p < 0.05$, corrected for multiple comparisons using a cluster-forming threshold of $Z$ = 3.1). The anatomical locations of these maxima were determined by referencing the Harvard-Oxford Cortical and Subcortical, and Cerebellar Atlases in MNI152 space following normalization with FLIRT. Only structures with a probability of 25% or higher are reported. For regions where the Harvard-Oxford atlas was insufficient for identification, manual identification was conducted. Brain areas manually identified using the Multi-Contrast Anatomical Subcortical Structures (MASSP) atlas (Bazin et al., 2020) are indicated in square brackets. Percentages represent the likelihood of anatomical localization.

| Index | Cluster size / voxel | *p* value | Z | Coordinates / mm | | | Name (Harvard-Oxford Atlas) |
|---|---|---|---|---|---|---|---|
| | | | | x | y | z | |
| *Both groups, acquisition training, CS x prediction* | | | | | | | |
| 1 | 7309 | < 0.001 | 7.08 | 6 | -8 | 66 | 47% Juxtapositional Lobule Cortex (formerly Supplementary Motor Cortex) |
| | | | 6.95 | -2 | 6 | 46 | 42% Juxtapositional Lobule Cortex (formerly Supplementary Motor Cortex) |
| | | | 6.80 | 22 | -42 | 70 | 31% Postcentral Gyrus, 29% Superior Parietal Lobule |
| | | | 6.69 | -10 | 18 | 32 | 38% Cingulate Gyrus, anterior division, 28% Paracingulate Gyrus |
| | | | 6.38 | 10 | 6 | 42 | 37% Cingulate Gyrus, anterior division, 25% Juxtapositional Lobule Cortex (formerly Supplementary Motor Cortex) |
| | | | 6.25 | 4 | -46 | 56 | 74% Precuneal Cortex |
| 2 | 5138 | < 0.001 | 6.91 | 10 | 8 | 8 | 85% Right Caudate |
| | | | 5.85 | -10 | 8 | 2 | 54% Left Caudate, 45% Left Cerebral White Matter |
| | | | 5.54 | 14 | -12 | 8 | 100% Right Thalamus |
| | | | 5.53 | 20 | 12 | -14 | [Right Striatum] |
| 3 | 2686 | < 0.001 | 6.64 | -26 | -64 | -20 | 94% Left VI |
| | | | 6.40 | -34 | -50 | -34 | 72% Left VI, 28% Left Crus I |
| | | | 5.53 | -38 | -58 | -26 | 56% Left Crus I, 34% Left VI |
| | | | 5.52 | -32 | -56 | -28 | 95% Left VI |
| | | | 5.43 | -46 | -60 | -32 | 100% Left Crus I |
| | | | 5.41 | 12 | -62 | -54 | 41% Right VIIIb, 31% Right IX |
| 4 | 1026 | < 0.001 | 5.71 | 64 | -40 | 22 | 68% Supramarginal Gyrus, posterior division |
| | | | 5.20 | 60 | -26 | 24 | 35% Supramarginal Gyrus, anterior division, 26% Parietal Operculum Cortex |
| | | | 5.08 | 54 | -26 | 24 | 41% Parietal Operculum Cortex |
| | | | 4.97 | 48 | -30 | 24 | 53% Parietal Operculum Cortex |
| | | | 4.86 | 58 | -44 | 8 | 50% Middle Temporal Gyrus, temporooccipital part |
| | | | 4.54 | 48 | -32 | -4 | 53% Middle Temporal Gyrus, posterior division |
| 5 | 954 | < 0.001 | 5.94 | -64 | -38 | 28 | 54% Supramarginal Gyrus, anterior division |
| | | | 5.65 | -56 | -40 | 34 | 33% Supramarginal Gyrus, anterior division, 28% Supramarginal Gyrus, posterior division |
| | | | 5.20 | -64 | -28 | 24 | 62% Supramarginal Gyrus, anterior division |
| | | | 5.09 | -50 | -32 | 24 | 46% Parietal Operculum Cortex |
| | | | 5.02 | -56 | -38 | 28 | 32% Parietal Operculum Cortex |
| | | | 5.01 | -52 | -20 | 28 | 28% Postcentral Gyrus |
| 6 | 878 | < 0.001 | 6.34 | 46 | 2 | 54 | 36% Middle Frontal Gyrus, 30% Precentral Gyrus |
| | | | 5.40 | 44 | -12 | 46 | 38% Precentral Gyrus |
| | | | 5.29 | 46 | 2 | 42 | 49% Precentral Gyrus |
| | | | 4.85 | 34 | -6 | 46 | 31% Precentral Gyrus |
| | | | 4.28 | 42 | 4 | 36 | 36% Precentral Gyrus, |
| 7 | 283 | < 0.001 | 4.94 | -36 | -8 | 56 | 48% Precentral Gyrus |
| | | | 4.58 | -46 | -10 | 52 | 72% Precentral Gyrus |
| | | | 3.95 | -42 | 2 | 60 | 33% Middle Frontal Gyrus |
| 8 | 217 | 0.001 | 4.89 | -38 | -60 | -56 | 60% Left VIIb, 25% Left VIIIa |
| | | | 4.68 | -24 | -66 | -56 | 79% Left VIIIa |
| | | | 3.71 | -38 | -48 | -50 | 49% Left VIIb, 35% Left VIIIa |
| | | | 3.54 | -30 | -58 | -50 | 50% Left VIIIa, 32% Left VIIb |
| | | | 3.51 | -14 | -64 | -50 | 56% Left VIIIa, 44% Left VIIIb |
| | | | 3.46 | -32 | -46 | -52 | 75% Left VIIIa |

| | | | | | | | |
|---|---|---|---|---|---|---|---|
| 9 | 217 | 0.001 | 4.19 | -10 | -78 | 46 | 39% Lateral Occipital Cortex, superior division, 25% Precuneal Cortex |
| | | | 4.12 | -8 | -72 | 46 | 52% Precuneal Cortex |
| | | | 4.00 | -8 | -78 | 36 | 38% Precuneal Cortex, 29% Cuneal Cortex |
| 10 | 134 | 0.010 | 4.64 | -22 | -44 | -56 | 86% Left VIIIb |
| | | | 4.35 | -6 | -44 | -52 | 92% Brainstem |
| | | | 4.26 | -12 | -52 | -56 | 49% Left IX, 45% Left VIIIb |
| | | | 4.03 | -10 | -46 | -52 | 45% Left IX, 25% Left VIIIb |
| | | | 3.94 | -14 | -46 | -54 | 43% Left VIIIb, 42% Left IX |
| *Both groups, acquisition training, no-US x prediction error* | | | | | | | |
| 1 | 5573 | < 0.001 | 8.66 | 44 | 20 | -6 | 40% Frontal Orbital Cortex |
| | | | 8.64 | 32 | 24 | -8 | 59% Frontal Orbital Cortex |
| | | | 8.32 | 36 | 26 | 0 | 34% Frontal Orbital Cortex, 26% Frontal Operculum Cortex |
| | | | 8.07 | 34 | 24 | 8 | 44% Frontal Operculum Cortex |
| | | | 7.99 | 34 | 22 | 4 | 47% Insular Cortex |
| | | | 7.93 | 44 | 20 | 2 | 56% Frontal Operculum Cortex |
| 2 | 4466 | < 0.001 | 6.54 | 2 | 8 | 56 | 30% Juxtapositional Lobule Cortex (formerly Supplementary Motor Cortex) |
| | | | 6.46 | 2 | 6 | 60 | 46% Juxtapositional Lobule Cortex (formerly Supplementary Motor Cortex) |
| | | | 6.15 | -2 | 14 | 38 | 56% Cingulate Gyrus, anterior division, 30% Paracingulate Gyrus |
| | | | 5.75 | 4 | 14 | 44 | 68% Paracingulate Gyrus |
| | | | 5.63 | -8 | 4 | 44 | 44% Juxtapositional Lobule Cortex (formerly Supplementary Motor Cortex) |
| | | | 5.56 | 4 | -6 | 68 | 64% Juxtapositional Lobule Cortex (formerly Supplementary Motor Cortex) |
| 3 | 3105 | < 0.001 | 7.44 | -46 | 14 | -4 | 33% Frontal Operculum Cortex |
| | | | 6.75 | -32 | 26 | 0 | 32% Frontal Orbital Cortex, 27% Insular Cortex |
| | | | 6.23 | -38 | 16 | -6 | 72% Insular Cortex |
| | | | 6.15 | -52 | 2 | 6 | 44% Central Opercular Cortex |
| | | | 5.88 | -36 | 12 | 8 | 48% Frontal Operculum Cortex |
| 4 | 2911 | < 0.001 | 6.19 | 52 | -30 | 30 | 25% Parietal Operculum Cortex |
| | | | 5.96 | 66 | -32 | 22 | 30% Superior Temporal Gyrus, posterior division |
| | | | 5.88 | 66 | -36 | 30 | 44% Supramarginal Gyrus, posterior division |
| | | | 5.77 | 50 | -28 | -2 | 62% Superior Temporal Gyrus, posterior division |
| | | | 5.53 | 50 | -22 | -6 | 42% Superior Temporal Gyrus, posterior division, 41% Middle Temporal Gyrus, posterior division |
| | | | 5.47 | 52 | -46 | 38 | 38% Angular Gyrus, 30% Supramarginal Gyrus, posterior division |
| 5 | 2438 | < 0.001 | 5.88 | -32 | -58 | -26 | 95% Left VI |
| | | | 5.58 | 22 | -66 | 8 | 46% Intracalcarine Cortex |
| | | | 5.01 | -16 | -70 | 10 | 50% Intracalcarine Cortex |
| | | | 4.90 | -12 | -76 | 16 | 25% Intracalcarine Cortex |
| 6 | 752 | < 0.001 | 5.52 | 8 | -22 | -18 | [Right Substantia Nigra] |
| | | | 5.51 | -8 | -10 | -12 | [Left Substantia Nigra] |
| | | | 5.42 | 20 | -16 | -6 | 75% Right Cerebral White Matter |
| | | | 5.22 | 6 | -24 | -2 | [Right Thalamus] |
| | | | 5.20 | 4 | -28 | -2 | 34% Brainstem, [Right Thalamus] |
| | | | 4.52 | 12 | -8 | 6 | 98% Right Thalamus |
| 7 | 690 | < 0.001 | 5.32 | -64 | -24 | 20 | 36% Supramarginal Gyrus, anterior division, 32% Postcentral Gyrus |
| | | | 4.76 | -58 | -32 | 22 | 53% Parietal Operculum Cortex |
| | | | 4.36 | -54 | -18 | 24 | 34% Postcentral Gyrus |
| | | | 4.27 | -64 | -44 | 30 | 48% Supramarginal Gyrus, posterior division |
| | | | 4.17 | -66 | -38 | 26 | 40% Supramarginal Gyrus, anterior division |
| 8 | 346 | < 0.001 | 4.89 | 36 | -56 | -26 | 89% Right VI |
| | | | 4.64 | 26 | -60 | -24 | 100% Right VI |
| | | | 4.63 | 40 | -52 | -32 | 90% Right Crus I |
| | | | 3.82 | 32 | -66 | -22 | 66% Right VI |
| 9 | 273 | < 0.001 | 4.79 | -32 | -64 | -52 | 75% Left VIIb |
| | | | 4.30 | -38 | -50 | -46 | 54% Left Crus II |
| | | | 4.22 | -22 | -70 | -48 | 80% Left VIIb |
| | | | 4.07 | -34 | -54 | -52 | 57% Left VIIIa, 32% Left VIIb |
| | | | 3.73 | -24 | -76 | -48 | 57% Left Crus II, 38% Left VIIb |
| 10 | 200 | < 0.001 | 5.21 | 20 | -46 | 74 | 35% Superior Parietal Lobule |
| | | | 5.15 | 18 | -52 | 72 | 49% Superior Parietal Lobule |
| | | | 5.14 | 18 | -46 | 70 | 37% Superior Parietal Lobule |

| | | | | | | | |
|---|---|---|---|---|---|---|---|
| 11 | 192 | 0.001 | 4.26 | -42 | -54 | 52 | 28% Angular Gyrus |
| | | | 4.14 | -50 | -50 | 56 | 32% Supramarginal Gyrus, posterior division |
| | | | 4.10 | -44 | -50 | 46 | 28% Supramarginal Gyrus, posterior division, 26% Angular Gyrus |
| | | | 4.05 | -28 | -54 | 48 | 38% Superior Parietal Lobule |
| 12 | 141 | 0.004 | 3.84 | 10 | -72 | 44 | 42% Precuneous Cortex |
| 13 | 105 | 0.019 | 4.60 | -2 | -50 | -22 | 88% Left I-IV |
| | | | 4.38 | 0 | -60 | -32 | 55% Vermis VIIIa |
| 14 | 253 | < 0.001 | -5.00 | -4 | -52 | 22 | 62% Cingulate Gyrus, posterior division |
| 15 | 124 | 0.008 | -4.22 | 36 | -24 | 50 | 29% Postcentral Gyrus |
| | | | -3.58 | 46 | -20 | 58 | 43% Postcentral Gyrus |
| | | | -3.42 | 40 | -24 | 66 | 41% Postcentral Gyrus, 30% Precentral Gyrus |
| 16 | 108 | 0.017 | -4.24 | 46 | -74 | -46 | 39% Right Crus II |
| | | | -3.82 | 44 | -76 | -36 | 80% Right Crus I |
| | | | -3.80 | 40 | -74 | -36 | 81% Right Crus I II |
| | | | -3.68 | 34 | -84 | -28 | 88% Right Crus I |
| *Sham group, acquisition training, CS x prediction* | | | | | | | |
| 1 | 2621 | < 0.001 | 5.78 | -10 | 20 | 32 | 37% Paracingulate Gyrus, 34% Cingulate Gyrus, anterior division |
| | | | 5.61 | 6 | -8 | 64 | 51% Juxtapositional Lobule Cortex (formerly Supplementary Motor Cortex) |
| | | | 4.97 | 12 | -22 | 40 | 57% Cingulate Gyrus, posterior division |
| | | | 4.85 | 2 | 2 | 50 | 55% Juxtapositional Lobule Cortex (formerly Supplementary Motor Cortex) |
| | | | 4.82 | 2 | 20 | 30 | 90% Cingulate Gyrus, anterior division |
| | | | 4.77 | 10 | 16 | 34 | 44% Cingulate Gyrus, anterior division |
| 2 | 1596 | < 0.001 | 4.85 | -58 | -16 | 30 | 65% Postcentral Gyrus |
| | | | 4.79 | -22 | 8 | 8 | 76% Left Putamen |
| | | | 4.61 | -34 | -6 | -12 | 96% Left Cerebral White Matter |
| | | | 4.57 | -52 | 8 | 4 | 36% Precentral Gyrus, 29% Inferior Frontal Gyrus, pars opercularis |
| | | | 4.38 | -40 | 2 | -4 | 72% Insular Cortex |
| 3 | 806 | < 0.001 | 4.57 | 44 | 22 | -4 | 37% Frontal Operculum Cortex, 28% Frontal Orbital Cortex |
| | | | 4.51 | 42 | 26 | -2 | 40% Frontal Orbital Cortex, 35% Frontal Operculum Cortex |
| | | | 4.41 | 60 | 8 | 6 | 45% Precentral Gyrus |
| | | | 4.40 | 32 | 22 | -8 | 43% Frontal Orbital Cortex, 35% Insular Cortex |
| 4 | 609 | < 0.001 | 5.29 | 54 | -22 | 20 | 46% Parietal Operculum Cortex |
| | | | 4.30 | 66 | -38 | 28 | 62% Supramarginal Gyrus, posterior division |
| | | | 4.28 | 62 | -38 | 22 | 51% Supramarginal Gyrus, posterior division |
| | | | 3.94 | 48 | -30 | 24 | 53% Parietal Operculum Cortex |
| | | | 3.62 | 58 | -18 | 32 | 39% Postcentral Gyrus, 34% Supramarginal Gyrus, anterior division |
| | | | 3.57 | 58 | -32 | 34 | 37% Supramarginal Gyrus, anterior division |
| 5 | 566 | < 0.001 | 6.07 | -34 | -50 | -34 | 72% Left VI, 28% Left Crus I |
| | | | 4.85 | -46 | -60 | -32 | 100% Left Crus I |
| | | | 4.72 | -26 | -64 | -20 | 94% Left VI |
| | | | 4.67 | -44 | -66 | -28 | 98% Left Crus I |
| | | | 4.17 | -30 | -56 | -28 | 100% Left VI |
| | | | 3.87 | -36 | -58 | -26 | 49% Left VI, 41% Left Crus I |
| 6 | 536 | < 0.001 | 5.87 | 46 | 0 | 54 | 44% Precentral Gyrus |
| | | | 5.46 | 48 | 0 | 42 | 40% Precentral Gyrus |
| | | | 5.06 | 50 | -2 | 48 | 66% Precentral Gyrus |
| | | | 3.96 | 58 | 6 | 40 | 43% Precentral Gyrus |
| | | | 3.90 | 46 | -12 | 48 | 41% Precentral Gyrus |
| | | | 3.81 | 60 | 0 | 42 | 36% Precentral Gyrus |
| 7 | 472 | < 0.001 | 4.69 | 40 | -50 | -30 | 61% Right Crus I, 39% Right VI |
| | | | 4.65 | 42 | -54 | -34 | 100% Right Crus I |
| | | | 4.25 | 36 | -42 | -36 | 86% Right VI |
| | | | 4.20 | 28 | -60 | -18 | 47% Right VI, 28% Temporal Occipital Fusiform Cortex |
| | | | 4.20 | 28 | -56 | -24 | 100% Right VI |
| | | | 4.20 | 22 | -52 | -24 | 71% Right VI, 29% Right V |
| 8 | 428 | < 0.001 | 5.17 | 16 | -62 | -52 | 65% Right VIIIb, 35% Right VIIIa |
| | | | 4.49 | 10 | -66 | -44 | 43% Right VIIIa |
| | | | 4.01 | 10 | -52 | -58 | 68% Right IX, 32% Right VIIIb |
| | | | 3.95 | 34 | -60 | -58 | 60% Right VIIIa, 35% Right VIIb |
| | | | 3.82 | -2 | -70 | -38 | 66% Vermis VIIIa |
| 9 | 371 | < 0.001 | 4.86 | 12 | -12 | 6 | 100% Right Thalamus |

| | | | | | | | |
|---|---|---|---|---|---|---|---|
| | | | 4.73 | 8 | -4 | 6 | 98% Right Thalamus |
| | | | 4.69 | 10 | 8 | 8 | 85% Right Caudate |
| | | | 4.57 | 12 | 0 | 12 | 31% Right Caudate |
| | | | 3.97 | 8 | -14 | -8 | [Right Red Nucleus] |
| | | | 3.81 | 12 | -12 | -2 | 67% Right Cerebral White Matter, [Right Thalamus] |
| 10 | 328 | < 0.001 | 4.35 | 14 | -40 | 50 | 29% Precuneal Cortex, 26% Postcentral Gyrus |
| | | | 4.04 | 12 | -46 | 54 | 39% Precuneal Cortex |
| | | | 3.96 | 6 | -52 | 62 | 62% Precuneal Cortex |
| | | | 3.81 | 6 | -44 | 56 | 57% Precuneal Cortex |
| 11 | 132 | 0.010 | 4.30 | -38 | -60 | -56 | 60% Left VIIb, 25% Left VIIIa |
| | | | 4.21 | -26 | -66 | -56 | 66% Left VIIIa, 34% Left VIIb |
| | | | 3.89 | -38 | -50 | -48 | 39% Left Crus II, 32% Left VIIb |
| 12 | 102 | 0.035 | 4.61 | -4 | -10 | 12 | 97% Left Thalamus |
| *Sham group, acquisition training, no-US x prediction error* | | | | | | | |
| 1 | 2624 | < 0.001 | 7.01 | 30 | 24 | -10 | 64% Frontal Orbital Cortex |
| | | | 6.80 | 44 | 22 | -6 | 43% Frontal Orbital Cortex |
| | | | 6.12 | 40 | 28 | -2 | 45% Frontal Orbital Cortex |
| | | | 6.00 | 48 | 20 | 8 | 39% Inferior Frontal Gyrus, pars opercularis, 26% Inferior Frontal Gyrus, pars triangularis |
| | | | 5.91 | 46 | 14 | 0 | 43% Frontal Operculum Cortex |
| | | | 5.70 | 46 | 20 | 0 | 42% Frontal Operculum Cortex |
| 2 | 1217 | < 0.001 | 5.44 | 2 | 8 | 60 | 34% Juxtapositional Lobule Cortex (formerly Supplementary Motor Cortex) |
| | | | 4.74 | -2 | 10 | 54 | 33% Paracingulate Gyrus |
| | | | 4.69 | 4 | 14 | 64 | 54% Superior Frontal Gyrus |
| | | | 4.56 | -4 | 18 | 38 | 52% Paracingulate Gyrus, 36% Cingulate Gyrus, anterior division |
| | | | 4.36 | -2 | 20 | 56 | 65% Superior Frontal Gyrus |
| | | | 4.28 | 2 | 24 | 56 | 47% Superior Frontal Gyrus |
| 3 | 1169 | < 0.001 | 5.31 | -44 | 12 | -2 | 39% Frontal Operculum Cortex |
| | | | 4.70 | -34 | 18 | -10 | 46% Insular Cortex, 28% Frontal Orbital Cortex |
| | | | 4.52 | -54 | 2 | 6 | 42% Central Opercular Cortex, 26% Precentral Gyrus |
| | | | 4.47 | -40 | 6 | -2 | 68% Insular Cortex |
| | | | 4.41 | -52 | 12 | -6 | 50% Temporal Pole |
| 4 | 582 | < 0.001 | 4.53 | 66 | -38 | 26 | 57% Supramarginal Gyrus, posterior division |
| | | | 4.42 | 48 | -48 | 30 | 36% Angular Gyrus |
| | | | 4.38 | 58 | -50 | 24 | 65% Angular Gyrus |
| | | | 4.29 | 64 | -40 | 34 | 68% Supramarginal Gyrus, posterior division |
| | | | 4.20 | 52 | -46 | 38 | 38% Angular Gyrus, 30% Supramarginal Gyrus, posterior division |
| | | | 4.12 | 56 | -44 | 46 | 57% Supramarginal Gyrus, posterior division |
| 5 | 426 | < 0.001 | 4.94 | 20 | -14 | -6 | 70% Right Cerebral White Matter |
| | | | 4.76 | 6 | -24 | -16 | 36% Brainstem, [Right VTA] |
| | | | 4.55 | -6 | -14 | -10 | [Right VTA] |
| | | | 4.43 | -12 | -6 | -16 | 27% Left Amygdala |
| | | | 4.38 | 6 | -24 | -2 | [Right Thalamus] |
| | | | 4.09 | -4 | -28 | 0 | 29% Left Thalamus, [Third Ventricle] |
| 6 | 168 | 0.001 | 4.71 | 48 | -30 | 2 | 52% Superior Temporal Gyrus, posterior division |
| | | | 4.31 | 54 | -18 | -6 | 54% Superior Temporal Gyrus, posterior division |
| | | | 3.50 | 58 | -26 | -6 | 56% Middle Temporal Gyrus, posterior division |
| 7 | 95 | 0.031 | 3.85 | -48 | -2 | 44 | 46% Precentral Gyrus |
| | | | 3.79 | -54 | 0 | 50 | 26% Precentral Gyrus |
| 8 | 90 | 0.039 | 4.77 | -18 | 4 | 0 | 65% Left Pallidum, 32% Left Putamen |
| | | | 4.09 | -20 | 4 | 10 | 68% Left Cerebral White Matter, 32% Left Putamen, [Left Internal Capsule] |
| | | | 3.96 | -26 | 2 | 10 | 96% Left Putamen |
| *Verum group, acquisition training, CS x Prediction* | | | | | | | |
| 1 | 1837 | < 0.001 | 5.90 | -4 | 6 | 48 | 51% Juxtapositional Lobule Cortex (formerly Supplementary Motor Cortex) |
| | | | 5.64 | -14 | 2 | 72 | 44% Superior Frontal Gyrus |
| | | | 5.30 | 4 | -8 | 66 | 61% Juxtapositional Lobule Cortex (formerly Supplementary Motor Cortex) |
| | | | 5.21 | -6 | -2 | 72 | 42% Superior Frontal Gyrus |
| | | | 4.97 | 6 | 10 | 40 | 51% Cingulate Gyrus, anterior division |
| | | | 4.95 | 10 | 4 | 44 | 39% Juxtapositional Lobule Cortex (formerly Supplementary Motor Cortex), 31% Cingulate Gyrus, anterior division |
| 2 | 1250 | < 0.001 | 6.62 | 22 | -42 | 72 | 29% Postcentral Gyrus |

| | | | | | | | |
|---|---|---|---|---|---|---|---|
| | | | 5.61 | 2 | -46 | 54 | 65% Precuneal Cortex |
| | | | 5.48 | -18 | -44 | 62 | 38% Postcentral Gyrus |
| | | | 5.17 | 18 | -50 | 72 | 50% Superior Parietal Lobule |
| | | | 5.05 | -20 | -50 | 68 | 47% Superior Parietal Lobule |
| | | | 4.32 | -6 | -50 | 68 | 37% Precuneal Cortex |
| 3 | 435 | < 0.001 | 5.07 | -56 | -38 | 36 | 42% Supramarginal Gyrus, anterior division |
| | | | 4.77 | -64 | -38 | 28 | 54% Supramarginal Gyrus, anterior division |
| | | | 4.44 | -50 | -36 | 26 | 50% Parietal Operculum Cortex |
| | | | 4.18 | -66 | -28 | 24 | 58% Supramarginal Gyrus, anterior division |
| | | | 3.71 | -62 | -22 | 22 | 53% Postcentral Gyrus |
| | | | 3.49 | -60 | -48 | 26 | 55% Supramarginal Gyrus, posterior division |
| 4 | 317 | < 0.001 | 5.20 | -10 | 8 | 2 | 54% Left Caudate |
| | | | 4.01 | -8 | 18 | 0 | 51% Left Caudate |
| | | | 4.01 | -18 | 12 | 0 | 57% Left Putamen, 43% Left Cerebral White Matter |
| | | | 3.65 | -8 | -6 | 10 | 98% Left Thalamus |
| | | | 3.18 | -2 | 2 | 8 | 55% Left Lateral Ventricle, 44% Left Cerebral White Matter |
| 5 | 226 | < 0.001 | 4.22 | 36 | 26 | 8 | 34% Frontal Operculum Cortex |
| | | | 4.18 | 34 | 28 | 2 | 30% Frontal Orbital Cortex |
| | | | 4.07 | 26 | 14 | -4 | 75% Right Putamen, 25% Right Cerebral White Matter |
| | | | 3.41 | 32 | 14 | 8 | 62% Insular Cortex |
| | | | 3.32 | 22 | 8 | 0 | 99% Right Putamen |
| 6 | 176 | 0.002 | 5.65 | 8 | 10 | 0 | 81% Right Caudate |
| | | | 5.15 | 10 | 8 | 8 | 85% Right Caudate, [Right Internal Capsule] |
| | | | 3.66 | 6 | -6 | 4 | 100% Right Thalamus |
| 7 | 155 | 0.004 | 4.35 | -42 | -60 | -26 | 84% Left Crus I |
| | | | 4.25 | -38 | -56 | -26 | 41% Left VI, 39% Left Crus I |
| | | | 3.72 | -50 | -52 | -28 | 37% Left Crus I |
| | | | 3.38 | -42 | -66 | -22 | 28% Left Crus I |
| 8 | 130 | 0.011 | 4.44 | -42 | 46 | 26 | 52% Frontal Pole |
| | | | 3.86 | -30 | 40 | 26 | 43% Frontal Pole |
| | | | 3.82 | -34 | 52 | 24 | 83% Frontal Pole |
| | | | 3.80 | -22 | 46 | 20 | 45% Frontal Pole |
| | | | 3.78 | -32 | 40 | 34 | 53% Frontal Pole, 26% Middle Frontal Gyrus |
| 9 | 118 | 0.018 | 4.49 | -32 | 24 | 8 | 46% Frontal Operculum Cortex |
| 10 | 106 | 0.030 | 4.13 | -44 | -6 | 54 | 70% Precentral Gyrus |
| | | | 3.99 | -36 | -8 | 56 | 48% Precentral Gyrus |
| *Verum group, acquisition training, no-US x prediction error* | | | | | | | |
| 1 | 3291 | < 0.001 | 7.29 | 34 | 26 | 0 | 28% Frontal Orbital Cortex, 26% Insular Cortex |
| | | | 6.46 | 42 | 20 | -2 | 38% Frontal Operculum Cortex |
| | | | 6.36 | 32 | 26 | -6 | 56% Frontal Orbital Cortex |
| | | | 5.58 | 42 | 8 | 4 | 52% Central Opercular Cortex |
| | | | 5.49 | 56 | 16 | 0 | 33% Inferior Frontal Gyrus, pars opercularis |
| 2 | 1965 | < 0.001 | 5.57 | -10 | 2 | 42 | 39% Juxtapositional Lobule Cortex (formerly Supplementary Motor Cortex), 28% Cingulate Gyrus, anterior division |
| | | | 5.55 | -10 | -2 | 42 | 31% Juxtapositional Lobule Cortex (formerly Supplementary Motor Cortex) |
| | | | 5.33 | 2 | 10 | 38 | 78% Cingulate Gyrus, anterior division |
| | | | 5.04 | 2 | -8 | 44 | 56% Cingulate Gyrus, anterior division |
| | | | 4.89 | 4 | 6 | 56 | 67% Juxtapositional Lobule Cortex (formerly Supplementary Motor Cortex) |
| | | | 4.86 | 8 | -4 | 72 | 28% Juxtapositional Lobule Cortex (formerly Supplementary Motor Cortex), 28% Superior Frontal Gyrus |
| 3 | 1071 | < 0.001 | 6.20 | -30 | 26 | 2 | 42% Insular Cortex |
| | | | 5.51 | -46 | 14 | -2 | 41% Frontal Operculum Cortex |
| | | | 5.50 | -40 | 16 | -4 | 63% Insular Cortex |
| | | | 5.25 | -36 | 16 | 8 | 70% Frontal Operculum Cortex |
| | | | 5.22 | -36 | 10 | 12 | 42% Frontal Operculum Cortex, 25% Central Opercular Cortex |
| | | | 4.98 | -50 | 0 | 2 | 69% Central Opercular Cortex |
| 4 | 762 | < 0.001 | 4.74 | -10 | -76 | 16 | 37% Intracalcarine Cortex |
| | | | 4.64 | -18 | -70 | 10 | 39% Intracalcarine Cortex |
| | | | 4.35 | -4 | -90 | 8 | 26% Occipital Pole |
| | | | 4.05 | -6 | -80 | 10 | 58% Intracalcarine Cortex |
| | | | 3.90 | -18 | -68 | -2 | 25% Lingual Gyrus |
| 5 | 669 | < 0.001 | 5.14 | 52 | -30 | 28 | 33% Parietal Operculum Cortex |
| | | | 4.33 | 64 | -38 | 8 | 33% Supramarginal Gyrus, posterior division |

| | | | | | | | |
|---|---|---|---|---|---|---|---|
| | | | 4.30 | 64 | -26 | 24 | 52% Supramarginal Gyrus, anterior division |
| | | | 4.11 | 66 | -42 | 20 | 57% Supramarginal Gyrus, posterior division |
| | | | 4.08 | 66 | -36 | 30 | 44% Supramarginal Gyrus, posterior division |
| 6 | 402 | < 0.001 | 5.10 | 40 | -54 | 46 | 36% Angular Gyrus |
| | | | 4.60 | 44 | -50 | 48 | 40% Angular Gyrus |
| | | | 4.48 | 38 | -48 | 46 | 33% Superior Parietal Lobule |
| | | | 4.17 | 42 | -42 | 40 | 36% Supramarginal Gyrus, posterior division |
| | | | 3.85 | 52 | -46 | 40 | 44% Angular Gyrus, 26% Supramarginal Gyrus, posterior division |
| | | | 3.62 | 54 | -42 | 48 | 61% Supramarginal Gyrus, posterior division |
| 7 | 295 | < 0.001 | 5.48 | -30 | -58 | -28 | 100% Left VI |
| | | | 3.70 | -48 | -56 | -32 | 99% Left Crus I |
| | | | 3.36 | -34 | -64 | -36 | 75% Left Crus I |
| 8 | 199 | < 0.001 | 4.60 | -52 | -34 | 28 | 28% Parietal Operculum Cortex, 18% Supramarginal Gyrus, anterior division |
| | | | 4.42 | -56 | -30 | 20 | 63% Parietal Operculum Cortex |
| | | | 3.85 | -66 | -24 | 20 | 33% Supramarginal Gyrus, anterior division |
| 9 | 175 | 0.001 | 4.36 | -26 | -76 | -48 | 67% Left Crus II |
| | | | 4.14 | -32 | -64 | -50 | 67% Left VIIb |
| | | | 4.09 | -26 | -70 | -50 | 79% Left VIIb |
| | | | 3.95 | -36 | -62 | -54 | 66% Left VIIb |
| | | | 3.75 | -36 | -56 | -54 | 49% Left VIIb, 47% Left VIIIa |
| 10 | 171 | 0.001 | 4.50 | 46 | -20 | -8 | 31% Superior Temporal Gyrus, posterior division |
| | | | 4.03 | 50 | -26 | 0 | 58% Superior Temporal Gyrus, posterior division |
| | | | 3.30 | 62 | -32 | -6 | 38% Middle Temporal Gyrus, posterior division |
| 11 | 142 | 0.004 | 4.41 | -8 | -76 | -36 | 62% Left Crus II |
| | | | 3.91 | -6 | -74 | -28 | 28% Left VI, 27% Left Crus I |
| | | | 3.67 | -14 | -74 | -34 | 56% Left Crus II, 34% Left Crus I |
| 12 | 123 | 0.009 | 4.04 | 28 | -54 | -32 | 85% Right VI |
| | | | 3.82 | 38 | -58 | -26 | 50% Right Crus I, 50% Right VI |
| | | | 3.60 | 24 | -64 | -24 | 90% Right VI |
| | | | 3.39 | 34 | -66 | -24 | 51% Right Crus I, 49% Right VI |
| | | | 3.15 | 22 | -64 | -32 | 51% Right VI |
| 13 | 122 | 0.009 | 4.98 | 18 | -52 | 72 | 49% Superior Parietal Lobule |
| 14 | 97 | 0.028 | 4.63 | -18 | 10 | -6 | 99% Left Putamen |
| | | | 3.87 | -14 | 14 | 4 | 79% Left Caudate, [Left Internal Capsule] |
| 15 | 89 | 0.041 | 4.69 | -18 | -44 | -54 | 85% Left VIIIb |
| | | | 3.74 | -14 | -52 | -56 | 74% Left VIIIb |
| 16 | 103 | 0.021 | -4.27 | -4 | -50 | 22 | 67% Cingulate Gyrus, posterior division |
| | | | -3.98 | -6 | -56 | 26 | 37% Precuneal Cortex, 37% Cingulate Gyrus, posterior division |
| *Acquisition training, CS x prediction, verum > sham* | | | | | | | |
| | No surviving clusters | | | | | | |
| *Acquisition training, CS x prediction, verum < sham* | | | | | | | |
| | No surviving clusters | | | | | | |
| *Acquisition training, noUS x prediction error, verum > sham* | | | | | | | |
| | No surviving clusters | | | | | | |
| *Acquisition training, noUS x prediction error, verum < sham* | | | | | | | |
| | No surviving clusters | | | | | | |

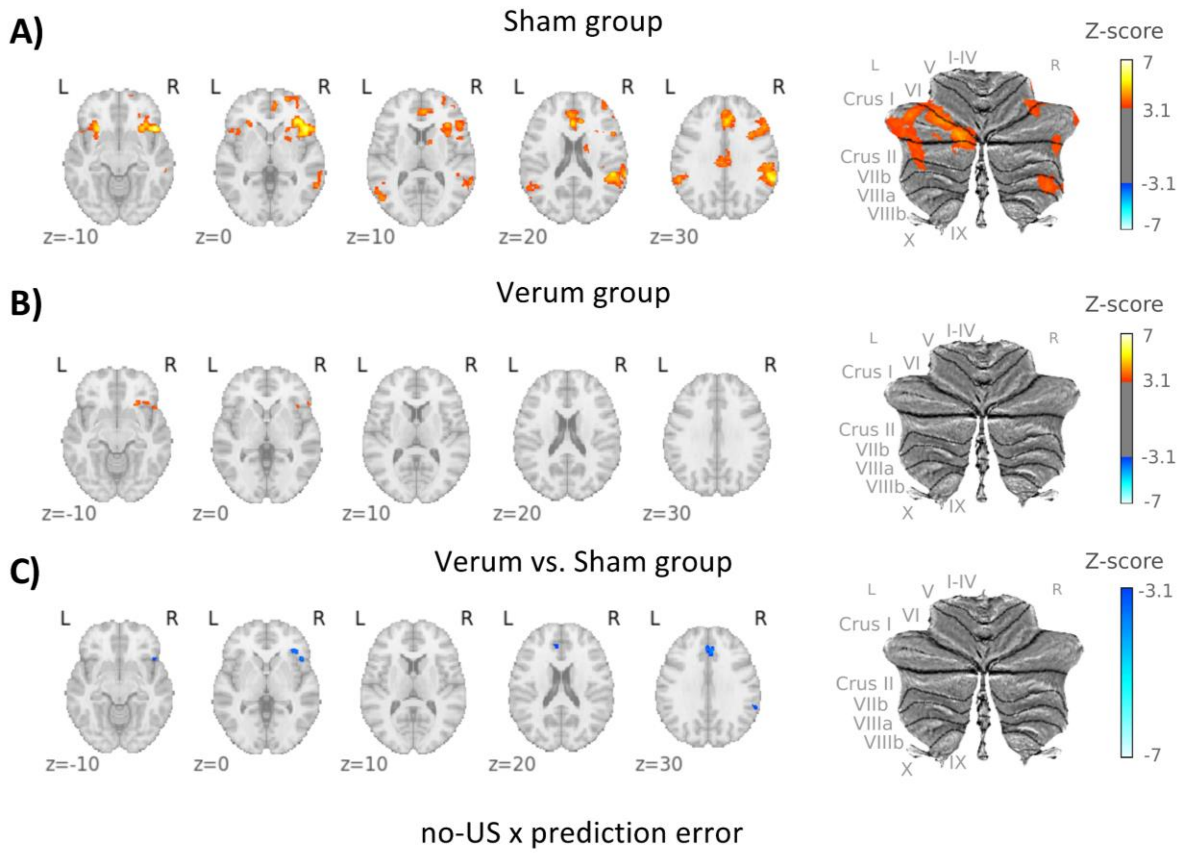


**Figure 6-1.** Parametric modulation with learning model-derived omission of US events at CS termination with individual absolute mean prediction error values (no-US x prediction error) during recall in **A)** the sham group, **B**) the verum group and **C)** for the comparison between verum and sham groups (contrast 'verum vs. sham'). In the individual groups, contrast increases are shown in red and decreases in blue.

In the differential contrast 'verum vs sham', shades of blue denote higher activation in the sham group, while shades of red denote higher activation in the verum group. CS = conditioned stimulus; US = unconditioned stimulus; L = left; R = right; SUIT = spatially unbiased atlas template of the cerebellum; Results of fMRI analysis are provided in **Table 6-2.**

**Table 6-1.** Fear extinction training. Parametric modulation of learning model-derived prediction parameters. Activation clusters are reported. Listed below are all local maxima within significant clusters ($p < 0.05$, corrected for multiple comparisons using a cluster-forming threshold of $Z = 3.1$). The anatomical locations of these maxima were determined by referencing the Harvard-Oxford Cortical and Subcortical, and Cerebellar Atlases in MNI152 space following normalization with FLIRT. Only structures with a probability of 25% or higher are reported. For regions where the Harvard-Oxford atlas was insufficient for identification, manual identification was conducted. Brain areas manually identified using the Multi-Contrast Anatomical Subcortical Structures (MASSP) atlas (Bazin et al., 2020) are indicated in square brackets. Percentages represent the likelihood of anatomical localization.

| Index | Cluster size / voxel | *p* value | Z | Coordinates / mm | | | Name (Harvard-Oxford Atlas) |
|---|---|---|---|---|---|---|---|
| | | | | x | y | z | |
| *Sham group, extinction training, CS x prediction* | | | | | | | |
| | No surviving clusters | | | | | | |
| *Sham group, extinction training, no-US x prediction error* | | | | | | | |
| 1 | 640 | < 0.001 | 5.31 | 48 | 24 | -2 | 33% Frontal Operculum Cortex |
| | | | 5.08 | 28 | 20 | -12 | 51% Frontal Orbital Cortex |
| | | | 4.91 | 44 | 20 | -6 | 40% Frontal Orbital Cortex |
| | | | 4.83 | 40 | 30 | -6 | 30% Frontal Orbital Cortex |
| | | | 4.71 | 38 | 24 | -8 | 64% Frontal Orbital Cortex |
| | | | 4.61 | 60 | 18 | 6 | 31% Inferior Frontal Gyrus, pars opercularis |
| 2 | 362 | < 0.001 | 5.77 | 40 | 14 | 36 | 45% Middle Frontal Gyrus |
| | | | 4.19 | 52 | 20 | 40 | 53% Middle Frontal Gyrus |
| | | | 4.18 | 54 | 16 | 40 | 30% Middle Frontal Gyrus |
| | | | 4.08 | 46 | 4 | 40 | 38% Precentral Gyrus |
| | | | 4.07 | 48 | 6 | 44 | 33% Middle Frontal Gyrus |
| | | | 4.06 | 46 | 8 | 40 | 35% Middle Frontal Gyrus |
| 3 | 277 | < 0.001 | 4.66 | 56 | -46 | 44 | 41% Supramarginal Gyrus, posterior division, 39% Angular Gyrus |
| | | | 4.43 | 54 | -48 | 34 | 54% Angular Gyrus |
| | | | 4.27 | 58 | -48 | 34 | 52% Angular Gyrus, 28% Supramarginal Gyrus, posterior division |
| 4 | 243 | < 0.001 | 5.56 | 8 | 34 | 58 | 45% Superior Frontal Gyrus |
| | | | 4.91 | 8 | 34 | 54 | 45% Superior Frontal Gyrus |
| *Verum group, extinction training, CS x Prediction* | | | | | | | |
| 1 | 3033 | < 0.001 | 5.94 | -4 | 10 | 48 | 53% Paracingulate Gyrus |
| | | | 5.83 | 10 | 40 | 16 | 48% Cingulate Gyrus, anterior division, 31% Paracingulate Gyrus |
| | | | 5.42 | 6 | 4 | 64 | 62% Juxtapositional Lobule Cortex (formerly Supplementary Motor Cortex) |
| | | | 5.35 | 10 | 10 | 48 | 33% Paracingulate Gyrus |
| | | | 5.34 | 10 | 2 | 48 | 57% Juxtapositional Lobule Cortex (formerly Supplementary Motor Cortex) |
| | | | 5.30 | 10 | 34 | 22 | 41% Cingulate Gyrus, anterior division, 33% Paracingulate Gyrus |
| 2 | 1247 | < 0.001 | 5.13 | 30 | 22 | -4 | 36% Insular Cortex |
| | | | 4.54 | 56 | 6 | 14 | 51% Precentral Gyrus |
| | | | 4.42 | 34 | 14 | 6 | 50% Insular Cortex |
| 3 | 609 | < 0.001 | 5.53 | -30 | 24 | -6 | 45% Frontal Orbital Cortex |
| | | | 5.38 | -30 | 26 | 2 | 42% Insular Cortex |
| | | | 4.36 | -44 | 20 | 4 | 44% Frontal Operculum Cortex |
| | | | 4.29 | -38 | 20 | 8 | 55% Frontal Operculum Cortex |
| | | | 4.18 | -36 | 2 | 12 | 49% Central Opercular Cortex |
| 4 | 540 | < 0.001 | 4.72 | -16 | -74 | 30 | 34% Cuneal Cortex, 27% Precuneal Cortex |
| | | | 4.38 | -14 | -76 | 38 | 27% Precuneal Cortex |
| | | | 3.94 | -8 | -78 | -6 | 54% Lingual Gyrus |
| | | | 3.77 | -14 | -66 | 10 | 53% Intracalcarine Cortex |
| 5 | 499 | < 0.001 | 5.08 | 8 | -92 | 14 | 58% Occipital Pole |
| | | | 4.75 | -6 | -98 | 14 | 59% Occipital Pole |
| | | | 4.70 | -4 | -90 | 14 | 26% Occipital Pole |
| | | | 4.09 | 10 | -82 | 0 | 34% Intracalcarine Cortex, 26% Lingual Gyrus |
| | | | 3.95 | 2 | -88 | 18 | 37% Cuneal Cortex |
| | | | 3.84 | 8 | -82 | 22 | 39% Cuneal Cortex |
| 6 | 313 | < 0.001 | 4.52 | 68 | -38 | 28 | 35% Supramarginal Gyrus, posterior division |

| | | | | | | | |
|---|---|---|---|---|---|---|---|
| | | | 4.12 | 60 | -40 | 24 | 51% Supramarginal Gyrus, posterior division |
| | | | 4.03 | 50 | -42 | 16 | 36% Supramarginal Gyrus, posterior division |
| | | | 3.93 | 58 | -44 | 12 | 26% Supramarginal Gyrus, posterior division, 26% Middle Temporal Gyrus, temporooccipital part |
| | | | 3.81 | 64 | -46 | 24 | 48% Angular Gyrus, 36% Supramarginal Gyrus, posterior division |
| 7 | 286 | < 0.001 | 9.31 | -58 | -46 | 34 | 51% Supramarginal Gyrus, posterior division |
| | | | 4.47 | -54 | -36 | 28 | 32% Parietal Operculum Cortex |
| | | | 4.14 | -60 | -50 | 12 | 32% Supramarginal Gyrus, posterior division, 30% Angular Gyrus |
| | | | 4.05 | -64 | -48 | 14 | 45% Supramarginal Gyrus, posterior division |
| | | | 4.04 | -62 | -50 | 6 | 51% Middle Temporal Gyrus, temporooccipital part |
| | | | 4.03 | -62 | -46 | 30 | 59% Supramarginal Gyrus, posterior division |
| 8 | 234 | < 0.001 | 4.51 | 4 | -18 | 42 | 69% Cingulate Gyrus, posterior division |
| | | | 4.40 | 12 | -22 | 42 | 53% Cingulate Gyrus, posterior division, 25% Precentral Gyrus |
| | | | 4.34 | 16 | -30 | 42 | 31% Precentral Gyrus |
| | | | 4.33 | 14 | -30 | 46 | 41% Precentral Gyrus |
| 9 | 184 | 0.001 | 4.53 | 34 | 44 | 34 | 83% Frontal Pole |
| | | | 4.42 | 32 | 44 | 30 | 76% Frontal Pole |
| | | | 3.84 | 30 | 38 | 28 | 41% Frontal Pole |
| | | | 3.75 | 28 | 56 | 32 | 45% Frontal Pole |
| | | | 3.48 | 30 | 52 | 28 | 85% Frontal Pole |
| | | | 3.45 | 34 | 44 | 24 | 67% Frontal Pole |
| 10 | 184 | 0.001 | 4.29 | -10 | -8 | 10 | 100% Left Thalamus |
| | | | 4.16 | -8 | 14 | 4 | 90% Left Caudate |
| | | | 3.95 | -8 | 6 | 4 | 65% Left Caudate, [Left Internal Capsule] |
| | | | 3.78 | -16 | 16 | -2 | 49% Left Caudate, 36% Left Cerebral White Matter, [Left Internal Capsule] |
| | | | 3.76 | -12 | 2 | 12 | 61% Left Caudate, 36% Left Cerebral White Matter, [Left Internal Capsule] |
| | | | 3.67 | -16 | -16 | 12 | 92% Left Thalamus |
| 11 | 146 | 0.005 | 4.90 | 20 | -48 | 74 | 39% Superior Parietal Lobule |
| | | | 3.38 | 32 | -48 | 68 | 66% Superior Parietal Lobule |
| 12 | 110 | 0.022 | 4.13 | -32 | 36 | 28 | 34% Middle Frontal Gyrus, 25% Frontal Pole |
| | | | 4.04 | -28 | 44 | 32 | 78% Frontal Pole |
| | | | 3.91 | -24 | 52 | 36 | 62% Frontal Pole |
| 13 | 99 | 0.036 | 4.31 | 20 | -58 | 4 | 35% Lingual Gyrus |
| | | | 3.93 | 16 | -62 | 6 | 47% Intracalcarine Cortex |
| | | | 3.25 | 18 | -60 | 12 | 41% Precuneal Cortex |
| 14 | 97 | 0.039 | 4.36 | 40 | -16 | 2 | 47% Insular Cortex |
| | | | 4.10 | 50 | -18 | -10 | 45% Middle Temporal Gyrus, posterior division, 37% Superior Temporal Gyrus, posterior division |
| | | | 3.38 | 58 | -26 | -8 | 58% Middle Temporal Gyrus, posterior division |
| 15 | 94 | 0.045 | 4.14 | 10 | -8 | 10 | 100% Right Thalamus |
| | | | 3.63 | 10 | -18 | 12 | 100% Right Thalamus |
| 16 | 93 | 0.047 | 4.09 | 50 | 0 | 52 | 60% Precentral Gyrus |
| | | | 4.00 | 48 | 4 | 52 | 33% Precentral Gyrus, 31% Middle Frontal Gyrus |
| | | | 3.87 | 38 | -4 | 46 | 30% Precentral Gyrus |
| | | | 3.46 | 44 | -6 | 50 | 54% Precentral Gyrus |
| *Verum group, extinction training, no-US x prediction error* | | | | | | | |
| 1 | 707 | < 0.001 | 4.88 | 32 | 24 | -2 | 51% Insular Cortex |
| | | | 4.73 | 28 | 22 | -14 | 74% Frontal Orbital Cortex |
| | | | 4.68 | 54 | 24 | 4 | 50% Inferior Frontal Gyrus, pars triangularis |
| | | | 4.28 | 56 | 24 | 12 | 44% Inferior Frontal Gyrus, pars triangularis, 29% Inferior Frontal Gyrus, pars opercularis |
| | | | 4.27 | 46 | 24 | -16 | 43% Frontal Orbital Cortex |
| 2 | 191 | 0.001 | 5.26 | -14 | -78 | -32 | 46% Left Crus II, 44% Left Crus I |
| | | | 3.58 | -22 | -76 | -42 | 80% Left Crus II |
| | | | 3.38 | -24 | -72 | -38 | 52% Left Crus II, 33% Left Crus I |
| | | | 3.25 | -14 | -76 | -44 | 47% Left Crus II, 43% Left VIIb |
| 3 | 186 | 0.001 | 4.48 | 48 | -18 | -8 | 47% Superior Temporal Gyrus, posterior division |
| | | | 4.38 | 54 | -28 | -2 | 58% Superior Temporal Gyrus, posterior division, 28% Middle Temporal Gyrus, posterior division |
| 4 | 162 | 0.002 | 4.37 | 4 | -28 | 32 | 81% Cingulate Gyrus, posterior division |
| | | | 4.11 | 4 | -18 | 30 | 57% Cingulate Gyrus, posterior division |
| | | | 4.08 | 0 | -22 | 30 | 75% Cingulate Gyrus, posterior division |
| 5 | 136 | 0.006 | 5.40 | 10 | 32 | 28 | 52% Paracingulate Gyrus |
| | | | 3.61 | 6 | 24 | 30 | 62% Cingulate Gyrus, anterior division |

| | | | | | | | |
|---|---|---|---|---|---|---|---|
| | | | 3.60 | 10 | 24 | 34 | 36% Paracingulate Gyrus |
| 6 | 127 | 0.008 | 4.06 | 54 | -46 | 24 | 42% Angular Gyrus |
| | | | 4.04 | 62 | -54 | 22 | 67% Angular Gyrus |
| | | | 4.01 | 68 | -42 | 20 | 29% Supramarginal Gyrus, posterior division |
| | | | 3.77 | 58 | -58 | 18 | 49% Angular Gyrus |
| | | | 3.47 | 68 | -42 | 26 | 25% Supramarginal Gyrus, posterior division |
| | | | 3.40 | 68 | -38 | 30 | 33% Supramarginal Gyrus, posterior division |
| 7 | 124 | 0.009 | 5.15 | 4 | 8 | 62 | 48% Juxtapositional Lobule Cortex (formerly Supplementary Motor Cortex) |
| *Extinction training, CS x prediction, verum > sham* | | | | | | | |
| | No surviving clusters | | | | | | |
| *Extinction training, CS x prediction, verum < sham* | | | | | | | |
| | No surviving clusters | | | | | | |
| *Extinction training, noUS x prediction error, verum > sham* | | | | | | | |
| | No surviving clusters | | | | | | |
| *Extinction training, noUS x prediction error, verum < sham* | | | | | | | |
| | No surviving clusters | | | | | | |

**Table 6-2.** Recall. Parametric modulation of learning model-derived prediction parameters. Activation clusters are reported. Listed below are all local maxima within significant clusters ($p < 0.05$, corrected for multiple comparisons using a cluster-forming threshold of $Z = 3.1$). The anatomical locations of these maxima were determined by referencing the Harvard-Oxford Cortical and Subcortical, and Cerebellar Atlases in MNI152 space following normalization with FLIRT. Only structures with a probability of 25% or higher are reported. For regions where the Harvard-Oxford atlas was insufficient for identification, manual identification was conducted. Brain areas manually identified using the Multi-Contrast Anatomical Subcortical Structures (MASSP) atlas (Bazin et al., 2020) are indicated in square brackets. Percentages represent the likelihood of anatomical localization.

| Index | Cluster size / voxel | *p* value | Z | Coordinates / mm | | | Name (Harvard-Oxford Atlas) |
|---|---|---|---|---|---|---|---|
| | | | | x | y | z | |
| *Sham group, recall, CS x prediction* | | | | | | | |
| 1 | 113 | 0.007 | 4.03 | 32 | 22 | 12 | 35% Frontal Operculum Cortex |
| 2 | 98 | 0.015 | 4.33 | 48 | 18 | 0 | 34% Frontal Operculum Cortex |
| | | | 3.86 | 56 | 16 | 2 | 47% Inferior Frontal Gyrus, pars opercularis |
| 3 | 88 | 0.025 | 3.75 | -52 | 18 | 10 | 55% Inferior Frontal Gyrus, pars opercularis |
| | | | 3.69 | -54 | 14 | 2 | 67% Inferior Frontal Gyrus, pars opercularis |
| | | | 3.45 | -52 | 10 | 6 | 53% Inferior Frontal Gyrus, pars opercularis |
| 4 | 86 | 0.028 | 4.24 | 10 | -10 | 72 | 25% Superior Frontal Gyrus |
| | | | 3.47 | 14 | -20 | 72 | 53% Precentral Gyrus |
| | | | 3.46 | 16 | -8 | 70 | 36% Superior Frontal Gyrus |
| *Sham group, recall, no-US x prediction error* | | | | | | | |
| 1 | 2988 | < 0.001 | 7.20 | 48 | 20 | -4 | 26% Frontal Operculum Cortex |
| | | | 5.86 | 36 | 30 | -2 | 33% Frontal Orbital Cortex |
| | | | 5.84 | 40 | 2 | 40 | 34% Precentral Gyrus |
| | | | 5.48 | 32 | 24 | -4 | 44% Insular Cortex |
| | | | 5.09 | 30 | 22 | -12 | 71% Frontal Orbital Cortex |
| | | | 4.94 | 38 | 4 | 52 | 36% Middle Frontal Gyrus |
| 2 | 2555 | < 0.001 | 5.82 | 62 | -44 | 30 | 51% Supramarginal Gyrus, posterior division, 29% Angular Gyrus |
| | | | 5.75 | 68 | -40 | 26 | 34% Supramarginal Gyrus, posterior division |
| | | | 5.28 | 62 | -46 | 38 | 42% Supramarginal Gyrus, posterior division, 31% Angular Gyrus |
| | | | 4.92 | 68 | -40 | 20 | 37% Supramarginal Gyrus, posterior division |
| 3 | 2027 | < 0.001 | 5.86 | 10 | 30 | 22 | 48% Cingulate Gyrus, anterior division |
| | | | 5.48 | 4 | 32 | 28 | 49% Paracingulate Gyrus, 45% Cingulate Gyrus, anterior division |
| | | | 5.16 | 0 | 28 | 20 | 85% Cingulate Gyrus, anterior division |
| | | | 5.07 | 4 | 42 | 44 | 66% Superior Frontal Gyrus |
| | | | 4.86 | 4 | 44 | 6 | 64% Cingulate Gyrus, anterior division, 33% Paracingulate Gyrus |
| 4 | 579 | < 0.001 | 4.81 | -60 | -44 | 34 | 56% Supramarginal Gyrus, posterior division |
| | | | 4.63 | -56 | -60 | 16 | 41% Angular Gyrus, 29% Lateral Occipital Cortex, superior division |
| | | | 4.28 | -46 | -54 | 16 | 34% Angular Gyrus |
| | | | 4.20 | -50 | -70 | 10 | 70% Lateral Occipital Cortex, inferior division |
| | | | 4.11 | -62 | -44 | 44 | 33% Supramarginal Gyrus, posterior division |
| 5 | 475 | < 0.001 | 5.80 | -28 | 20 | -8 | 36% Insular Cortex |
| | | | 4.78 | -30 | 10 | -8 | [Left Putamen] |
| | | | 4.46 | -40 | 20 | -6 | 38% Frontal Orbital Cortex |
| 6 | 435 | < 0.001 | 4.73 | -28 | -68 | -30 | 89% Left Crus I |
| | | | 4.33 | -36 | -66 | -26 | 81% Left Crus I |
| | | | 4.30 | -26 | -74 | -26 | 89% Left Crus I |
| | | | 4.17 | -46 | -56 | -32 | 100% Left Crus I |
| | | | 3.96 | -38 | -56 | -46 | 58% Left Crus II I |
| | | | 3.68 | -42 | -62 | -40 | 81% Left Crus I |
| 7 | 361 | < 0.001 | 4.87 | 22 | 58 | -2 | 64% Frontal Pole |
| | | | 3.97 | 42 | 54 | 6 | 94% Frontal Pole |
| | | | 3.84 | 40 | 52 | 20 | 82% Frontal Pole |
| | | | 3.77 | 36 | 60 | -2 | 75% Frontal Pole |
| | | | 3.74 | 28 | 62 | 4 | 81% Frontal Pole |
| | | | 3.72 | 36 | 44 | 4 | 44% Frontal Pole |
| 8 | 193 | < 0.001 | 5.12 | -12 | -76 | -30 | 65% Left Crus I, 25% Left Crus II |

| | | | | | | | |
|---|---|---|---|---|---|---|---|
| 9 | 186 | < 0.001 | 4.04 | -4 | -24 | 30 | 56% Cingulate Gyrus, posterior division |
| | | | 4.02 | -2 | -20 | 32 | 74% Cingulate Gyrus, posterior division |
| | | | 4.01 | 4 | -18 | 30 | 57% Cingulate Gyrus, posterior division |
| | | | 3.65 | 4 | -8 | 30 | 46% Cingulate Gyrus, anterior division |
| 10 | 136 | 0.004 | 4.05 | 36 | -50 | -40 | 49% Right Crus I |
| | | | 3.91 | 32 | -50 | -34 | 80% Right VI |
| | | | 3.89 | 32 | -48 | -48 | 65% Right VIIIa |
| | | | 3.64 | 34 | -54 | -54 | 70% Right VIIIa, 25% Right VIIb |
| | | | 3.45 | 40 | -62 | -26 | 94% Right Crus I |
| | | | 3.28 | 36 | -42 | -38 | 55% Right VI, 40% Right Crus I |
| 11 | 99 | 0.021 | 4.24 | 8 | -28 | 46 | 40% Cingulate Gyrus, posterior division, 38% Precentral Gyrus |
| 12 | 90 | 0.032 | 3.91 | 26 | 50 | 40 | 58% Frontal Pole |
| | | | 3.76 | 22 | 54 | 38 | 48% Frontal Pole |
| | | | 3.46 | 32 | 42 | 40 | 69% Frontal Pole |
| | | | 3.11 | 26 | 36 | 46 | 38% Frontal Pole |
| *Verum group, recall, CS x prediction* | | | | | | | |
| | No surviving clusters | | | | | | |
| *Verum group, recall, no-US x prediction error* | | | | | | | |
| 1 | 228 | < 0.001 | 4.76 | 30 | 26 | -4 | 34% Frontal Orbital Cortex |
| | | | 4.15 | 52 | 18 | -8 | 28% Temporal Pole |
| | | | 3.83 | 40 | 24 | -8 | 64% Frontal Orbital Cortex |
| | | | 3.81 | 56 | 24 | 0 | 47% Inferior Frontal Gyrus, pars triangularis |
| | | | 3.75 | 46 | 18 | -4 | 33% Frontal Operculum Cortex |
| | | | 3.28 | 38 | 26 | -16 | 85% Frontal Orbital Cortex |
| *Recall, CS x prediction, verum > sham* | | | | | | | |
| | No surviving clusters | | | | | | |
| *Recall, CS x prediction, verum < sham* | | | | | | | |
| | No surviving clusters | | | | | | |
| *Recall, noUS x prediction error, verum > sham* | | | | | | | |
| | No surviving clusters | | | | | | |
| *Recall, noUS x prediction error, verum < sham* | | | | | | | |
| 1 | 241 | < 0.001 | 4.29 | 4 | 32 | 28 | 49% Paracingulate Gyrus, 45% Cingulate Gyrus, anterior division |
| | | | 4.15 | 0 | 34 | 28 | 55% Paracingulate Gyrus, 26% Cingulate Gyrus, anterior division |
| | | | 3.92 | -10 | 40 | 22 | 48% Paracingulate Gyrus |
| | | | 3.80 | -6 | 34 | 44 | 41% Superior Frontal Gyrus |
| | | | 3.40 | -4 | 28 | 36 | 80% Paracingulate Gyrus |
| | | | 3.33 | 0 | 24 | 34 | 44% Paracingulate Gyrus, 35% Cingulate Gyrus, anterior division |
| 2 | 122 | 0.007 | 4.30 | 48 | 22 | -2 | 29% Frontal Operculum Cortex |
| 3 | 99 | 0.021 | 3.81 | 64 | -42 | 26 | 70% Supramarginal Gyrus, posterior division |
| | | | 3.77 | 68 | -40 | 26 | 34% Supramarginal Gyrus, posterior division |
| | | | 3.67 | 62 | -46 | 38 | 42% Supramarginal Gyrus, posterior division, 31% Angular Gyrus |
| | | | 3.52 | 58 | -46 | 48 | 35% Angular Gyrus, 30% Supramarginal Gyrus, posterior division |
